# Shining a light on camouflage evolution: using genetic algorithms to determine the effects of geometry and lighting on optimal camouflage

**DOI:** 10.1101/2025.03.01.640995

**Authors:** George R.A. Hancock, Innes C. Cuthill, Jolyon Troscianko

## Abstract

Visual camouflage evolves within the bounds of light’s interaction with the surroundings and the sensory limits of its observers. Rapid temporal variation in lighting from weather and its interaction with objects within the visual scene alters the contrast of the spatio-chromatic features of both backgrounds and animals, the latter through self-shading and received shadows from their surroundings. Despite the apparent effect of lighting on object appearance, the enormity of interactions and the diversity of animal phenotypic solutions present challenges to investigating the combined effects of lighting and habitat structure on camouflage effectiveness and design. Genetic algorithms and mathematical animal pattern generation provide a potential solution to investigating this high-dimensional feature space. Here, an online artificial evolution experiment was used to examine the effect of lighting and habitat geometry on camouflage. Lighting and geometry changed which prey phenotypes evolved, and the predictive power of common camouflage metrics. Crucially, lighting condition systematically altered prey-targets’ internal contrast and interacted with habitat geometry, affecting the evolved patterning, colour, and countershading. Our work demonstrates the importance of considering the relative geometry and lighting of an environment when determining the function of animal colouration and the adaptive value of camouflage.

## Introduction

Animal camouflage has long served as an example of evolution by natural selection, as the colours and patterns used by animals reflect the characteristics of their local environment as seen by the visual systems of their observers, be they predators or prey (Endler 1987; Cuthill 2019). Effective camouflage, however, does not simply rely on matching the colour and pattern of the local background (Endler 1987; Merilaita 2003; Stevens and Ruxton 2019) (Stevens et al. 2015; Troscianko et al. 2016). Recognition, saliency, edge disruption and shape destruction all play a role in camouflage (Skelhorn et al. 2010; Webster et al. 2013; Webster 2015; Skelhorn and Rowe 2016; Pike 2018; Kelley et al. 2022). Background matching itself must function within the variable appearances of an animal’s background through space and time. Most animals move and so interact with – and are observed against – different surfaces and backgrounds (Merilaita et al. 2001; Briolat et al. 2021). Meanwhile, the range, frequency and pattern of colours in natural environments shift with the seasonality of their biotic (e.g., plant phenology) and abiotic (e.g., surface moisture) components (Endler 1984; Jones et al. 2018). Even if animals and their surrounding surfaces were somehow locked in time, their visual appearance would still vary from changes in lighting conditions caused by weather and the sun’s position (Cuthill et al. 2016, 2019). Weather in particular poses a challenge to camouflage as passing clouds and wind-swept foliage can rapidly change the lighting of scenes (Matchette et al., 2018, 2019). This instability of lighting necessitates behavioural, morphological and life history specialisations to maintain effective camouflage and conversely signalling (Stevens and Ruxton 2019). In this paper, we combine artificial animal pattern generation, genetic algorithms and mass-participation online games to explore the evolution of camouflage in environments with different lighting conditions.

Firstly, what should camouflage specialised to fixed or variable lighting environments look like, and how does it function? To understand these questions, we must first summarise how lighting changes the visual scene. On a cloudless day, in open habitat, the lighting of the environment is intense and directional. Surfaces facing the source of light are illuminated more intensely, while surfaces obstructed by their own or surrounding structures are in shadow. Shadows cast onto an object’s surface by itself (self-shadows) and onto surrounding surfaces (cast shadows) can provide cues to their shape and location (Casati 2004; Rensink and Cavanagh 2004). Obstruction of light does not simply change the luminance contrast; it also changes the ratio of longwave to shortwave light, because objects in shadow are illuminated by the sky, not the sun, and Rayleigh scattering is wavelength-dependent (Endler 1993; Narasimhan and Nayar 2002). For human trichromat vision, this results in an increase in the blue-yellow contrast of the scene (Troscianko et al. 2004), with shaded regions appearing bluer compared to regions in direct sunlight (Endler, 1993). Meanwhile, atmospheric conditions (e.g., cloud/fog) or the sun’s elevation (e.g., dusk/dawn) can scatter the light illuminating the scene, resulting in a diffuse lighting environment. Scenes with diffuse lighting still possess depth cues from self-shading, although they areless contrasting (Penacchio et al., 2015; Cuthill et al., 2016), and lack salient cast shadows (Mavrovouna et al. 2021; Kelley et al. 2022). How these changes influence the detectability of animals is poorly understood. The increased internal contrast and the presence of cast shadows under direct lighting may allow animals to be more easily distinguished from their background while increased visual complexity of the background from shadows and disruption of the animal’s shape from received shadows may instead hamper detection.

The disparities between visual scenes under different atmospheric conditions are influenced by the surrounding geometry of the scene. Flat surfaces allow 3-dimensional (3D) objects to cast unobstructed shadows, providing salient cues for their depth and location within the scene (Hancock et al. 2025).Within geometrically complex environments, light reflects from objects and scatters, becoming more diffuse, before reaching the scene or target object (Endler, 1993). Environments with tall vegetation – relative to the observer or target – therefore have less intense lighting, but can also feature dynamic dappled lighting and shadows from the movement of leaves and branches (Endler 1993; Matchette et al. 2018; Cuthill et al. 2019). This, in turn, can affect the detectability of objects (Matchette et al. 2018, 2019). Even shorter ground cover can influence the conspicuousness of cast shadows and self-shadows. Coarse surfaces, such as vegetation, distort the shape of and interfere with shadows (Dee and Santos 2011; Hancock et al. 2025). Tall thin branches and grasses have the added effect of creating thin directional shadows on surfaces, changing the uniformity of direction of patterns within the scene (directionality/anisotropy) under direct lighting. These sharp contrasting shadows disappear under diffuse lighting, though geometry will still affect and create soft shadows from differences in depth. Thus, how lighting influences camouflage should depend on the frequency of different atmospheric conditions and the geometry of an animal’s habitat across spatial scales relative to its size.

Existing research on how changes in the directionality and intensity of light from weather and geometry influence animal camouflage has largely been observational for all but a few mechanisms. Flat habitats and surfaces are thought to select for flatter postures and body plans in marine (flat-fish), terrestrial (bearded dragons) and trunk-dwelling species (leaf-tailed geckos, moths) (Stevens and Ruxton 2019). Flattening reduces the conspicuousness of cast and self-shadows and other pictorial depth cues. In the inverse setting, shadows of closed 3D complex habitats are thought to select for darker and or more complex and contrasting luminance patterns based on phylogenetic and ecological comparisons of felid coat patterns (Ortolani 1999; Caro 2005; Allen et al. 2011; Cheng et al. 2018; Marcondes et al. 2021). Experimentation on pattern adaptation to light environments has predominantly focused on countershading and edge-enhancement camouflage. Comparisons of the patterns of ungulates have shown that the intensities of their countershading gradients (the downward increase in luminance from their dorsal to ventral surface) are higher in species that occupy open environments at lower latitudes, where the intensity and directionality of the lighting environment are expected to be greater and more consistent (Allen et al. 2012). These observations are supported by experiments showing that counter-shaded models survived for longer if they had countershading gradients optimised to their self-shading gradient (Cuthill et al. 2016). While edge enhancement patterns which pictorially mimic shadows that form at the edges of objects from direct lighting (Osorio and Srinivasan 1991) are harder to find under direct rather than diffuse illumination (Egan et al. 2016).

How shadows should influence the orientation and colour of animal patterns is less clear. Cuttlefish can adaptively adjust the size and shape of their pattern to match the geometric and pictorial properties of their surrounding and produce shadow-mimicking patterns within directly illuminated 3D scenes (Barbosa et al. 2008; El Nagar et al. 2021). While a number of other invertebrates, fish, amphibians and reptiles can adjust their luminance, albeit at a slower rate, most terrestrial animals cannot perform rapid adaptive colouration and must make trade-offs or specialise to their environment (Duarte et al. 2017). The reduced stability of blue-yellow compared to green-red in neural opponent processing under variable lighting is thought to one factor that drives the use of red colouration in animal signalling and for the detection of camouflaged predators by primates (Lovell et al. 2005; Arenas et al. 2014; Pessoa et al. 2014). But does the stability of information change selection for background matching within these channels? Do long shadows from 3D complex habitats select for more contrasting and directional patterns and how might changes in light paths from local geometry affect camouflage strategies such as countershading, edge enhancement and glossiness (Allen et al. 2011; Cuthill et al. 2016; Egan et al. 2016; Penacchio et al. 2018)? How does the directionality of light interact with features of an animal’s texture that affect camouflage, e.g. glossiness (Thomas et al. 2023)? Moreover, how camouflage colours and patterns should be generalised to changes in background scene statistics from the weather is unknown. To simplify this host of potential phenotypic interactions with the directionality of light and habitat geometry, we have provided a table of predictions (Table 1).

**Table 1.**
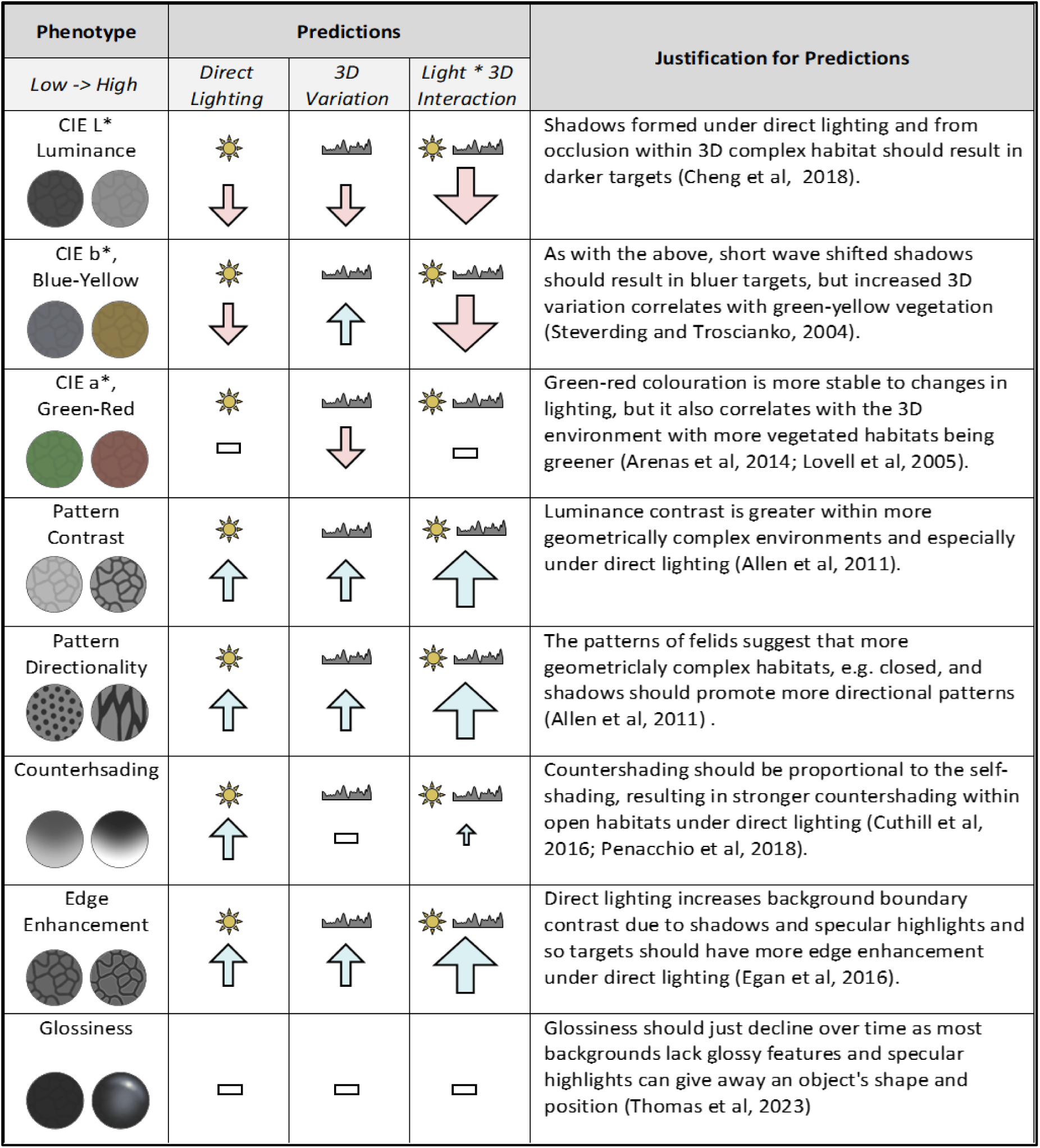
A summary of the predicted effects of lighting condition, habitat 3D variation and the interaction between direct lighting and 3D variation on the appearance of animal patterns. Upwards arrows indicate a positive effect on the phenotypic feature, e.g., increased luminance or directionality, downwards arrows indicate a negative effect, and a horizontal line indicates no effect. Larger arrows indicate that increased variation increases the effect size of direct lighting, while smaller arrows indicate a reduction in effect size. For each prediction, the justification and relevant references are provided. CIE L*, a* and b* refer to the axes widely used CIELAB perceptual colour space, namely luminance (L*), red-green (a*) and yellow-blue (b*) (CIE, 2007).

The vast array of interactions between lighting and geometry present fundamental challenges to examining their effects on camouflage (Fennell et al. 2019). The scene’s 3D geometry must be measured at multiple spatial scales and its visual appearance imaged under different weather conditions. Simultaneously, a large gamut of both intersecting and non-overlapping camouflage strategies must be compared. Here we aimed to provide a cohesive investigation of the broad effects of lighting and geometry on optimal animal camouflage colouration and patterns. We opted for a heuristic search approach, using an online citizen science game derived from the CamoEvo toolbox to disentangle camouflage optimisation within habitats of varying 3D and spatio-chromatic variability and under direct and diffuse lighting conditions (Hancock and Troscianko 2022). CamoEvo combines genetic algorithms (GAs), Gray-Scott reaction-diffusion patterns (aka Turing patterns), and human visual search experiments to compare the effects of background features on camouflage optimisation (Allen et al. 2011; Fennell et al. 2019, 2021). CamoEvo and similar natural selection-inspired mechanisms, such as other GAs and generative adversarial networks, have proved successful in creating functional animal-like camouflage patterns against different experimental backgrounds (Bond and Kamil 2002; Talas et al. 2020; Fennell et al. 2021). By using a GA to compare camouflage optimisation between lighting conditions, we can determine whether and how lighting influences the pattern shape, colours and specular reflection of targets, while also validating the circumstances where characteristics known to be influenced by lighting, e.g., countershading and edge enhancement, are under selection (Cuthill et al. 2016; Egan et al. 2016). A full table of predictions is provided below.

## Methods

### a Visual Scenes

For our backgrounds, we used a set of 28 temperate habitats photographed across the UK as part of a previous study on the effects of lighting and geometry on the patterns of visual scenes and objects (Hancock et al. 2025). These backgrounds included grassland, heathland, woodland and estuarine habitats. All photographs were taken from a height of 1.2 m with an ASUS Zenfone (ZenFone AR ZS571KL, AsusTek Computer Inc., Taipei, Taiwan) and calibrated with the MICA image analysis toolbox (Moher Alsady et al. 2016). For each habitat, the same 24 scenes were photographed under direct and diffuse lighting conditions. Direct lighting conditions were days with <10% cloud and no less than 2 h from sunrise or sunset. The diffuse lighting photographs of the same background were taken under a 1.5 m^3^ photography tent placed over the scene (NEEWER., Bee Block Inc, 15 Cotters Lane, East Brunswick, NJ 08816, USA), simulating the isotropic lighting conditions of cloudy days (Figure 1). Within each scene, a 3D-printed 30 mm diameter target painted grey with a matt 8% reflectance paint was placed at the centre. This target was used to normalise each scene using the MICA toolbox, as well as for rendering the evolved pattern. Additionally, for each lighting condition, a repeat photograph was taken with a black unpainted target soaked with acetone to give it a ‘glossy’ texture. 3D scans of the scenes were taken from the same camera position using the phone’s built-in Tango-enabled 3D scanner (Tango being Google’s 3D mapping technology) (Hancock et al. 2023). All scenes were limited to having some unobstructed lighting present (e.g., dappling in woodland), though the target itself could be placed within shadow or direct light within the scene. Habitat 3D variation was measured as the standard deviation of height across the scene averaged across all 24 scenes. Backgrounds with higher 3D variation featured vegetation and other tall objects.

**Figure 1.**
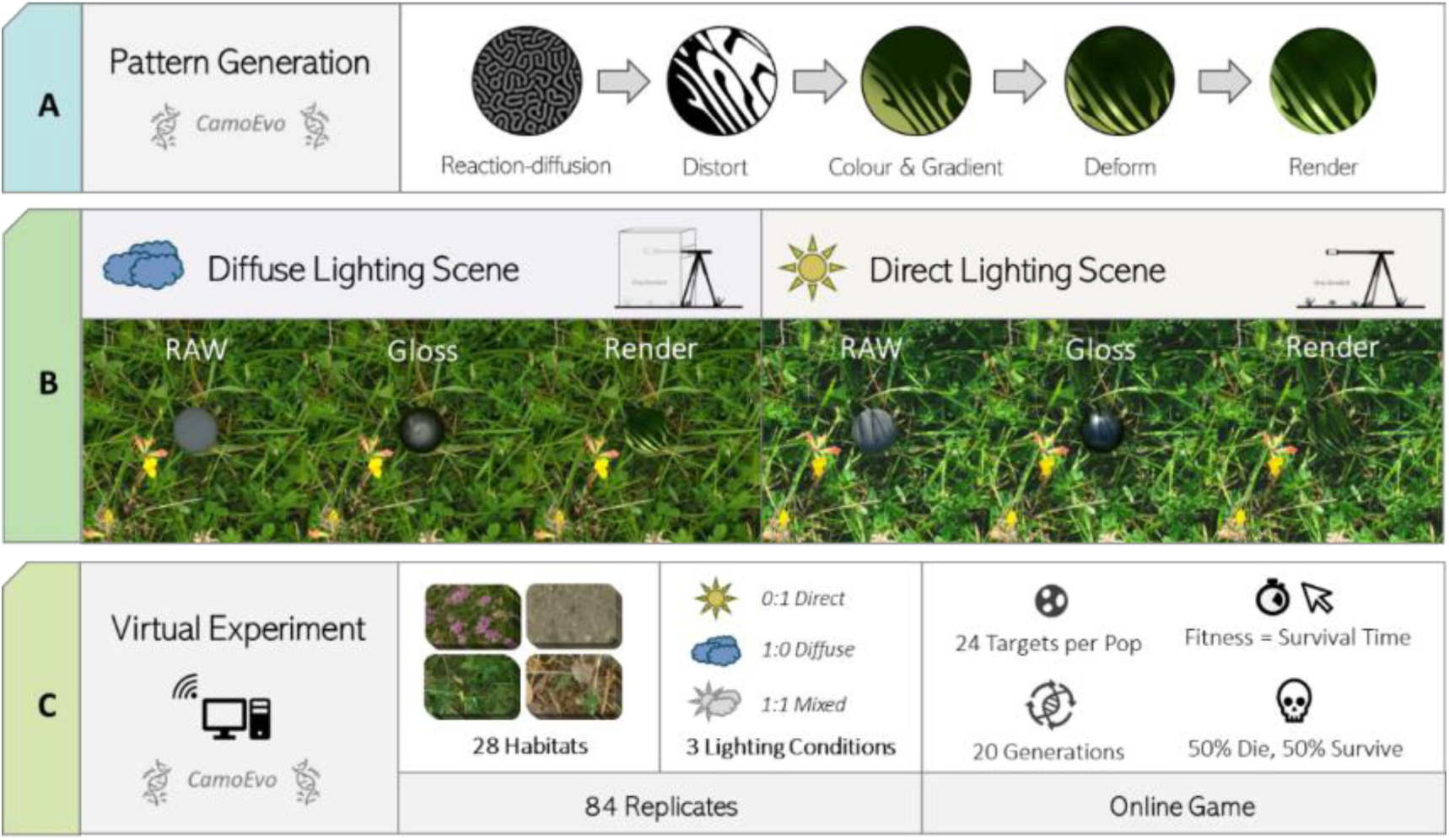
Schematic for experimental design using a meadow habitat as a demo. (A) Prey pattern generation using CamoEvo patterns, spherical deformation and rendering. (B) Calibrated raw, gloss and rendered targets under diffuse (left) and direct (right) lighting conditions. (C) Virtual online predation experiment using 28 different habitats and 3 lighting conditions. Each treatment evolved for 20 generations with 24 targets per population. The selection of targets was based on the time taken to click on the targets. The target pattern used was a sample pattern shown from the last generation (gen = 20) of the direct lighting-only evolved condition for the chalk wildflower meadow habitat.

### b Pattern Creation

To create our evolving targets, we used a modified version of CamoEvo’s pattern generator (Hancock and Troscianko 2022). CamoEvo uses Gray-Scott reaction-diffusion generated maculations, CIELAB colours (CIE 2007) and additional image filters to provide edge enhancement and countershading, creating biologically relevant patterns (Figure 1) These phenotypic traits of the target patterns were all controlled by 34 decimal genes on a haploid chromosome. These genes in turn were passed to CamoEvo’s genetic algorithm to mutate, recombine and select individuals. A spherical pattern deformation function was applied to the patterns for our experiment to create realistic 3D wrapping, and two additional genes/traits were added for controlling glossiness of the maculation and glossiness the background pattern. For full details of this system, see our CamoEvo paper and the supplementary material.

### c Stimuli

To show our evolving patterns under the lighting conditions of the scene, the patterns were rendered onto the calibrated grey target by first subtracting the target by its known reflectance and then multiplying it by the RGB values of the CamoEvo pattern. To change the target from matt to glossy, the RGB values of the glossy targets were pasted with addition (see supplementary material A). Background images with rendered patterns were then randomly cropped to create the stimuli for our online experiment. Each image was 1904 ×1488 px, and the target button had a diameter of 80 px. The target positions were never less than 2x their diameter from the centre or border of the image. These no-go zones were to prevent biases to fitness within my visual search experiment from being too close to the periphery of the observer’s vision or immediately fixated at the centre of the screen.

### d Online Experiment Design

For our game the 28 habitats were used for three different lighting treatments: two specialist lighting treatments, where all background images were from direct (DIRECT) or diffuse (DIFFUSE) and a mixed treatment, where half of the backgrounds were under diffuse and direct lighting (MIXED). This gave a total of 84 replicates, 28 per lighting treatment. For each replicate a starting population of 24 prey targets were generated using CamoEvo (Hancock and Troscianko 2022). The genes of the starting population were set with a uniform distribution from 0-1 across all individuals to prevent initial clustering of genotypes within populations. These targets were rendered onto the target button and backgrounds to create the stimuli above. An online computer-based search game was created from existing JavaScript to be played on internet browsers: https://camoevo.visual-ecology.com/ (Troscianko et al. 2021). The goal being to use humans as a surrogate predator to drive selection for the artificial prey for 20 generations (0-20).

Whenever a player played the game, they were randomly assigned one of the 84 populations and were tasked with finding and capturing the targets as fast as they could. Each population was shown to one participant and once it had been played it could not be played again until all other populations at the same concurrent generation had been played to prevent one population from being ahead of the others. All 24 stimuli and their respective targets were displayed in random order. At the start of the game, instructions were provided on what the game was for and what the players needed to do to play the game. As a control for starting viewpoint, participants were tasked with moving their cursor to the centre of the screen before initialising each slide or clicking in the centre for touchscreens. A ring at the centre of the screen marked where the participant needed to move their cursor to, with the instruction to not move their cursor until they had spotted the target. To capture the targets, participants needed to click on them within a time window of 15 s. The capture time was coded as the time taken to click on the target, while response time was encoded as the time taken to leave the centre circle. Fitness was encoded as the response time unless a delay of 600 ms was exceeded or the response time was faster than the stimulus display, in which case the capture time was used in place of response time, dubbed survival time, following the same protocol as used for the CamoEvo toolbox (Hancock and Troscianko 2022). Response and capture time were equal for touch screen users. Using response time and a centre fixation point minimises fitness noise caused by cursor travel time (Troscianko et al., 2017b).

Before any member of the target population was shown, a demo target was displayed. This demo consisted of a randomly generated pattern for the first generation (Gen) and then a deceased pattern from the previous generation for all subsequent generations. The purpose of this demo-target was to allow the participant to create an initial search image for the targets, which might otherwise make the first target harder to find and so have a disproportionately high fitness (Troscianko et al. 2021). Once a population had been completed, it was removed from the server, patterns were ranked by fitness, and CamoEvo’s genetic algorithm was used to create the next generation. For each round of selection, the 12 lowest (50%) ranking members of the population died, and new hybrid offspring were created by random pairing and recombination of the survivors from the previous generations. The new population for the next generation was then uploaded to the server to be played. This process was repeated until all populations had undergone 20 generations of selection and evolution. As an additional control for noise, the top 4 highest ranking targets from the generation prior to selection were given a lifeline against deletion, again following the same method used for CamoEvo (Hancock and Troscianko 2022).

### e Colour Analyses

For each scene, the 3D scans and direct and diffuse lighting images were rescaled to the minimum number of pixels per mm of all target images and converted to human CIELAB colour space (CIE 2007). For each channel (L*, a* and b*), the contrast (standard deviation) at 6 different spatial scales (spatial frequencies relative to target diameter, [1/64, 1/32, 1/16, 1/4, 1/1, 1/2, 1/1] x) and 4 different orientations [0, 45, 90, 135] were measured with Gabor filters with custom plugins for ImageJ v 1.53 (Talas et al. 2017; Van Den Berg et al. 2020; Barnett et al. 2021). These measures were taken for the target pattern before rendering (target-skin), the target pattern post rendering (rendered-target), the local surround (circle radius x2.5 target radius) and the global surround (circle radius x15 target radius). The ‘directionality’ of patterns was measured for each spatial scale by dividing the orientation with the maximum contrast by the mean contrast. Likewise, the ‘verticalness’ of patterning was calculated by subtracting the contrast filtered at 90 degrees from the contrast at 0 degrees, such that negative values indicated horizontal patterns and positive values vertical patterns. For the background, we also measured the 3D variability as the standard deviation of depth for the global surround. As a measure of habitat spatial variation in luminance, green-red, blue-yellow and depth we measured the mean contrast of each channel across all spatial scales, L_variation_, a_variation_, b_variation_, 3D_variation_ respectively.

To measure camouflage, we used metrics previously used for computer visual search and field experiments (Troscianko et al. 2016, 2017; Price et al. 2019). All metrics used have been shown to be able to predict the conspicuousness of prey. Luminance match was measured as the absolute difference between the mean of the L* channel of the local background and the rendered-target (Luminance_Difference_), while colour match was the Euclidean distance in the green-red (a*) and blue-yellow (b*) plane (Colour_Difference_) (Troscianko et al. 2021). Pattern difference was measured as the sum of absolute difference in contrast at each spatial scale for the luminance channel (Pattern_Difference_) (Troscianko et al. 2016). Target edge disruption was measured using GabRat of the target’s surrounding in the luminance channel, with the default sigma of 3, against its original background (Troscianko et al. 2017). Novel pattern measures were also used. To measure countershading and self-shading, the linear gradient from the top to the bottom of the target was measured for the CIELAB channels. Target gradients were measured for the target-skins, averaged direct lighting raw-targets for each habitat (see supplementary material A) and the targets rendered onto the averaged raw-target of its corresponding habitat. A positive gradient indicated that the top of the target was lighter than the bottom. Target glossiness was measured by calculating the difference in luminance value (mean and Std Dev) with and without the gloss genes active for each rendered target, as well as with the maximum gloss. Maximum glossiness was measured as the luminance and pattern of the targets could also affect glossiness; darker targets appeared glossier than lighter targets. Edge enhancement was measured as the difference in luminance pattern contrast at 4 spatial frequencies [1/64, 1/32, 1/16, 1/8] between the target with and without (no edge enhancement) the edge enhancement genes active.

### f Statistical Analyses

Statistics were performed with R, version 4.3.0 (R Core Team 2023). All metrics for target phenotype, camouflage and background structure were continuous and were re-scaled to a mean=0 and Std Dev=1, lighting treatment was given as a categorical variable with three-factor levels (DIRECT, DIFFUSE and MIXED). To test the effects of lighting and habitat 3D variation (3D_variation_) on our proxy for fitness (survival time) we used linear mixed models with the lme4 package (Douglas Bates et al. 2015). The rescaled log survival time was given as our predictor variable with generation number (N-Gen), lighting and 3D_variation_ as our fixed effects with the interaction between the three variables. We also compared the effect of lighting and 3D_variation_ on the predictive power of our camouflage metrics (Luminance_Difference,_ Colour_Difference,_ Pattern_Difference,_ GabRat_Difference_) for fitness, using each camouflage metric, instead of N-Gen, as the fixed effect. For all analyses, the habitat and population were used as random effects to account for changes caused by lighting between populations within the same habitat and individual population effects. For example, *lmm(log(SurvivalTime) ∼ CamouflageMetric * Light * 3D + (1|Habitat) + (1|Population) … where the * indicates interaction between fixed effects*.

To compare the effects of lighting on the evolution of phenotypes predicted to be affected by direct lighting and its interaction with geometry (pattern contrast, countershading, pattern directionality, pattern orientation, edge enhancement and glossiness) we first compiled the pattern metrics of the target-skin (contrast, directionality, verticality and gradient in the L*, a* and b* channels). These phenotype metrics were then used in a principal component analysis to account for multicollinearity between variables and whether treatment explained any of the principal components. We then used a similar model structure to the above to compare how lighting and geometry affected the evolution of principal components and our phenotype metrics. The phenotype metrics and principal components were used as the response variables, while N-Gen, lighting and 3D_variation_ were used as the fixed effects with population and habitat as random factors. For example, lmm (PhenotypeMetric ∼ N-Gen * Light * 3D + (1|Habitat) + (1|Population) …To ensure model assumptions were met, all residuals were checked for homogeneity of variance and homoscedasticity. Square root and log transformations were used as required to meet data distribution criteria.

## Results

For brevity, unless stated otherwise, all effects are significant at p<0.001, the conservative threshold being necessitated by the large number of tests conducted. Full model output tables are provided in supplementary material B.

### 1. Camouflage: Fitness, Match and Edge Disruption

We received the total of 1,764 individual players necessary for the experiment over 12 months, totalling 42,336 target presentations of which only 312 were timeouts. To assess whether targets became more camouflaged, the change in survival time, i.e., our fitness proxy, with generation number was measured (Figure 2). On average, fitness improved with n-generations (model not using lighting treatment or 3D complexity as predictors; N-Gen, β = 0.035, t_42250_ = 40.37) with a starting median survival time of 649 (IQR = 667.5) milliseconds (ms) and an ending median of 1026 (IQR = 1130.25) ms. Of the 84 treatment populations, 65 improved, and the remaining 19 (DIRECT=4, DIFFUSE=6, MIXED=10) either did not significantly increase in fitness or decreased in fitness with n-generations. The evolved fitness was highest for the population evolved against the gravel background under mixed lighting, with a median survival time of 1321 (IQR ± 3119) ms. Populations, where n-generations lowered survival time (N=12), were excluded from subsequent analyses, barring the habitat appearance analysis below.

**Figure 2.**
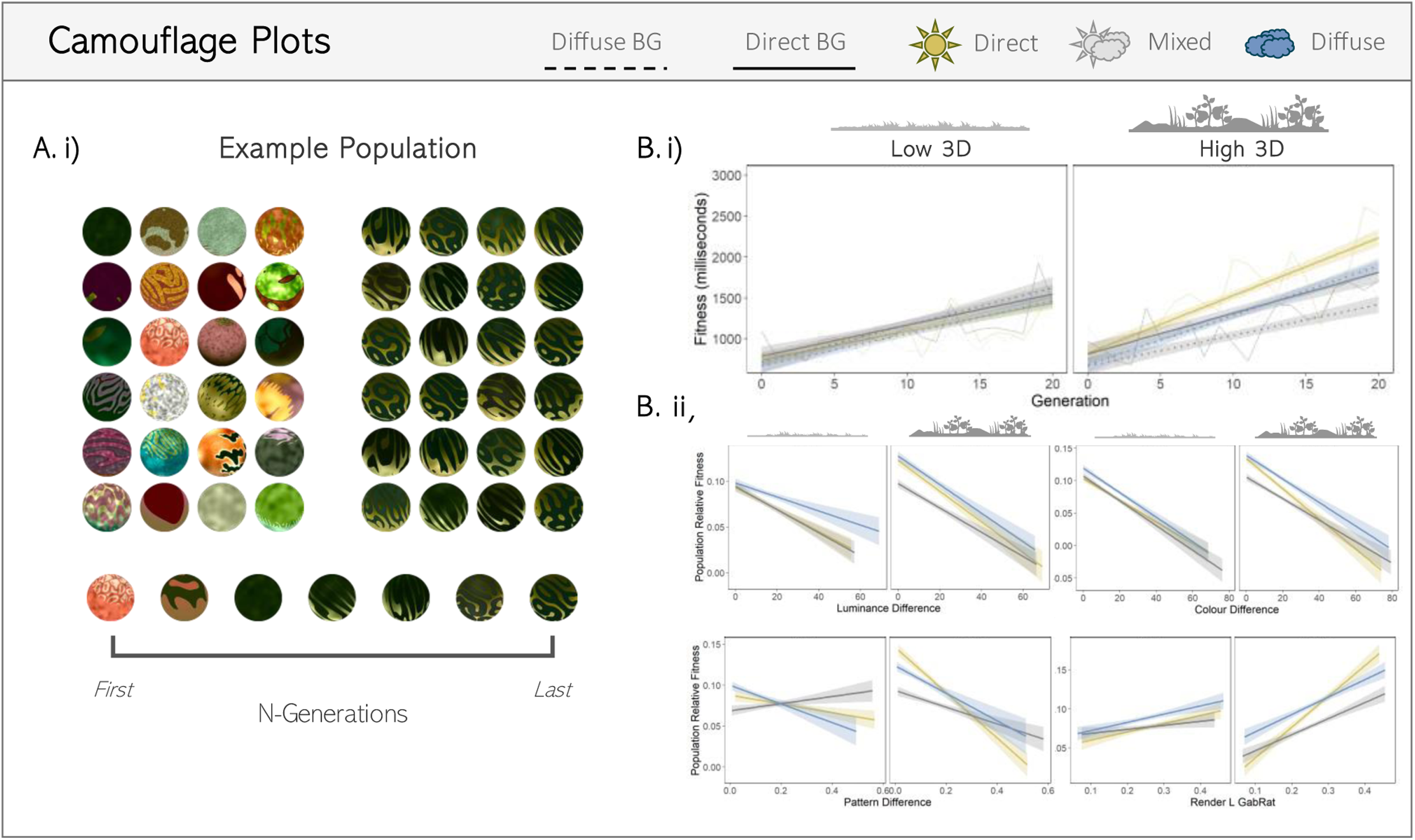
Evolution of Camouflage. A.i) Example change in phenotype for a population that improved in fitness. B.i) Average capture time increased over successive generations. B.ii) Panes showing the log predictive value of common camouflage metrics for survival time, luminance difference (absolute CIE L*difference), colour difference (Euclidean CIE a* and b* difference), pattern match (sum CIE L* contrast difference across scales) and edge disruption (GabRat L*). Lower values indicate a greater match to the background, while higher GabRat indicates greater edge disruption. Habitat 3D variation is split by the quantiles for the habitat’s mean depth variation across spatial scales.

We tested the effects of spatio-chromatic variation/complexity in the L*, a* and b* channels as well as 3D variation on the slope of fitness with n-generations to determine whether properties of the background influenced the evolution of camouflage. In doing so we found that the slope of fitness was significantly higher for populations on more variable backgrounds for all 4-background metrics (L, a, b and 3D variation, ‘*’ removed from models to avoid confusion with interaction symbology). Whether the population had a negative slope for survival time with n-generations was used as a random factor. The greatest effect size of any habitat metric for fitness and fitness with n-generations was observed for high variance in the combined L* and a* channels (La_variation_, β = 0.034, t_42190_ = 35.10; La_variation_ * N-Gen, β= 0.0127, t_,42330_ = 13.5). The highest fitness effect of any individual background metric was from background L variance (L_variation_, β = 0.0273, t_42330_ = 15.00; L _variation_* N-Gen, β= 0.0009, t_42330_ = 5.767), while simultaneously having the lowest effect on fitness with n-generations. Whereas, for improvement in fitness with n-generations it was background B variance which (B _variation_, β = 0.0020, t_42330_ = 11.35, p =0.276 | B * N-Gen, β = 0.0022, t_42330_ = 14.24). Most populations that failed to significantly improve in survival time were evolved on less variable backgrounds.

As direct lighting increases background luminance and blue-yellow variation, both of which increased fitness (reduced detectability), it is unsurprising that direct-lighting-only treatments evolved targets with a greater fitness then in mixed and diffuse treatments (model using lighting and 3D as factors, as described in methods; N-Gen, β = 0.050, t_36210_ = 31.08; N-Gen * MIXED, β = -0.009, t_36210_ = -4.17; N-Gen * DIFFUSE, β = -0.004, t_36210_ = -1.90). As 3D variation correlates with spatio-chromatic variation, targets evolved to be significantly harder to find when their habitat’s 3D variation was higher (N-Gen * 3D_variation_, β = 0.009, t_36210_ = 5.90), but had lower survival times for <u>diffuse</u> and <u>mixed</u> lighting treatments (DIFFUSE * 3D_variation_, β= -0.025, t_2,41_ = -2.43, p= 0.020; MIXED * 3D_variation_, β = -0.022, t_2,41_ = -2.10, p= 0.042). For populations evolved under mixed lighting, targets had lower survival times when against diffuse lighting backgrounds (Effect of background type on survival times for MIXED treatments; Diffuse_Background_, β = -0.014, t_12070_ = -4.33) especially when habitat 3D variation was high (Diffuse_Background_ * 3D_variation_, β= -0.011, t_12070_ = -3.39).

Luminance difference, colour difference, pattern difference and edge disruption were all significant predictors of survival time. Colour difference (β = -0.043, t_35500_ = -23.53) was the best predictor of survival time across habitats, followed by edge disruption (β = 0.040, t_33860_ = 19.93), pattern difference (β = -0.036, t_34420_ = -18.66) and lastly luminance difference (β= - 0.036, t_35780_ = -20.21). All camouflage metrics had significantly steeper slopes for 3D complex habitats. The effect size of camouflage metrics was impacted by lighting treatment. Diffuse only evolved targets had shallower slopes for luminance match (Luminance_Difference_ * Diffuse, β = 0.007, t_35910_ = 2.75, p = 0.006), colour match (Colour_Difference_ * Diffuse, β = 0.005, t_35780_ = 2.19, p= 0.028), and edge disruption (GabRat * Diffuse, β = -0.014, t_34050_ = -5.17). Whereas mixed lighting populations had a shallower slope for pattern match (Pattern_Difference_ * MIXED, β = 0.023, t_34240_ = 8.58) and edge disruption (GabRat * MIXED, β = -0.009, t_31400_ = -3.32, p= 0.001). For mixed lighting evolved targets, colour and pattern difference had an even lower effect on survival time when 3D complexity was high (Colour_Difference_ * MIXED * 3D_variation_, β = 0.008, t_35910_ = 3.12, p= 0.002; Pattern_Difference_ * MIXED * 3D_Variation_, β = 0.011, t_35480_ = 3.70), where the difference in colour and pattern between diffuse and direct lighting was greater.

### 2. Colour: Mean and Contrast (Std Dev)

Following the observed decrease in absolute colour difference between targets and their background, the colour space of targets in the final populations reflected the colour space of their habitat and lighting treatment (Figure 3). Notably, within 3D complex habitats, targets evolved to be, and were fitter when, darker than their local background, but less so for DIFFUSE populations (see supplementary material A). The principal component analysis of the target-skin features (Contrast, directionality and verticality in the L*, a* and b* channels) found PC1 (15.66% variance explained) to consist predominantly of pattern contrast in b*, L*, then a* greater than or equal to 1/4x the target’s spatial frequency and a decrease in verticality at those same scales, i.e., increase in horizontal contrast. PC2 (9.76% var) was predominantly colour verticality at intermediate scales, PC3 was higher green-red (a*) contrast across spatial scales (7.16% var) and PC4 was higher luminance verticality and directionality at small spatial scales (6.77% var). PC1 decreased with n-generations (N-Gen, β = -0.042, t_36210_ = -4.97), with PC1 being higher for DIRECT populations (DIFFUSE * N-Gen, β = -0.201, t_36210_ = - 16.63; MIXED * N-Gen, β = -0.096, t_36210_ = -8.04) and both DIRECT and MIXED populations against more 3D complex habitats but not diffuse (3D * N-Gen, β = 0.026, t_36210_ = 3.04, p= 0.002; MIXED * 3D * N-Gen, β = 0.015, t_36210_ = 1.24, p= 0.214; DIFFUSE * 3D * N-Gen, β = - 0.072, t_36210_ = -5.99). As the pattern contrast and PC1 increases were for the un-rendered targets, differences in target luminance and blue-yellow contrast observed between lighting conditions was not from the self or received shadows on the target and were entirely driven by pattern phenotype evolution.

**Figure 3.**
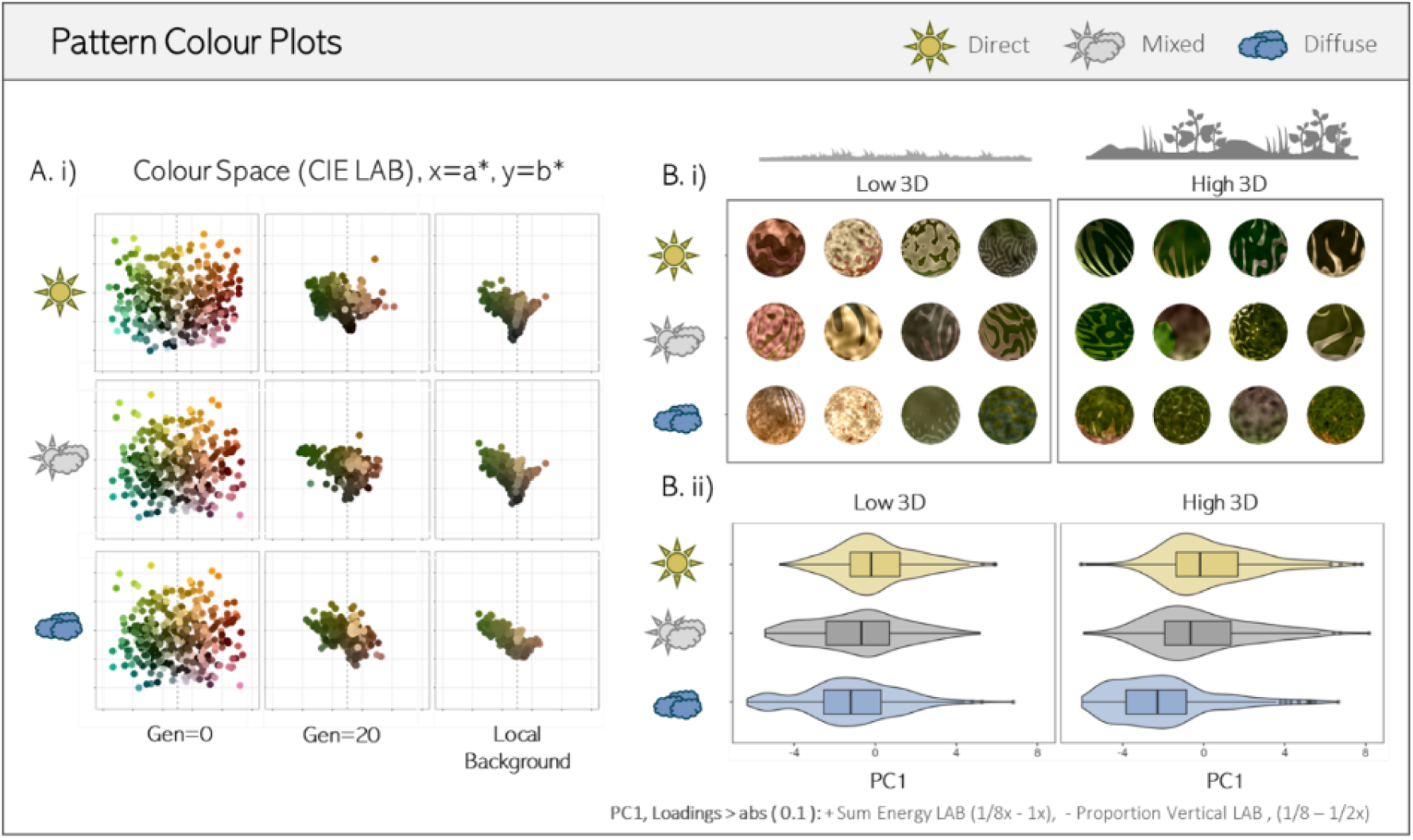
Change in colour space and colour patterns. A.i) Populations become increasingly similar to the mean colours of their local backgrounds (2.5x target radius) starting from a dispersed colour space. B.i) Example evolved targets from populations in low and high 3D variation habitats selected for under the different lighting treatments. B.ii) Hybrid violin & boxplots of PC1 for target patterns, where PC1 is predominantly large-scale luminance and colour contrast, and large horizontal (- vertical) patterns. Habitat 3D variation is split by the quantiles for the habitats’ mean depth variation across spatial scales.

### 3. Pattern: Gradient and Shape

As suggested by the above PC1, targets evolve larger scale horizontal patterns for direct lighting populations. Countershading requires a dark-to-light dorso-ventral gradient that flattens the internal contrast from self-shading (Rowland 2009). This gradient should in turn be proportional to the intensity of the raw luminance gradient from self-shading with a steeper gradient required for flatter, more open habitats where the targets have more intense self-shading gradient. Target populations in direct-lighting-only treatments evolved a more positive gradient (i.e., stronger countershading) (N-Gen, β = 0.022, t_36210_ = 20.99) than those in mixed lighting (N-Gen * MIXED, β = -0.008, t_36210_ = -5.36), and particularly compared to those under diffuse only lighting (N-Gen * DIFFUSE, β = -0.018, t_36210_ = -12.00) when rendered onto the raw direct lighting target of the corresponding habitat (Figure 4). So, direct lighting targets evolved to be more effective at countershading with mixed lighting as an intermediate. However, the gradient increase was steeper for more 3D variable habitats (3D_variation_ * N-Gen, β = 0.011, t_36210_ = 10.11; 3D_variation_ * DIFFUSE * N-Gen, β = -0.008, t_36210_ = -5.18), i.e., when the raw target’s luminance gradient was lower rather than higher (Allen et al. 2012; Cuthill et al. 2016). This reduction of gradient corresponded with the target-skin having a positive luminance gradient and a darker colouration. Although differences in background and target luminance alone could not explain the reduced gradient for the targets within flatter habitats. Comparison of countershading when splitting the habitats by their dominant substrate (bare, gravel, leaflitter, grass and misc-vegetation) found countershading did not affect survival time for gravel habitats and had a lower effect for closed-leaflitter habitats (Figure 4, Table 2). So, lighting environment and geometry both affected and interacted to affect the adaptive value of countershading.

**Figure 4.**
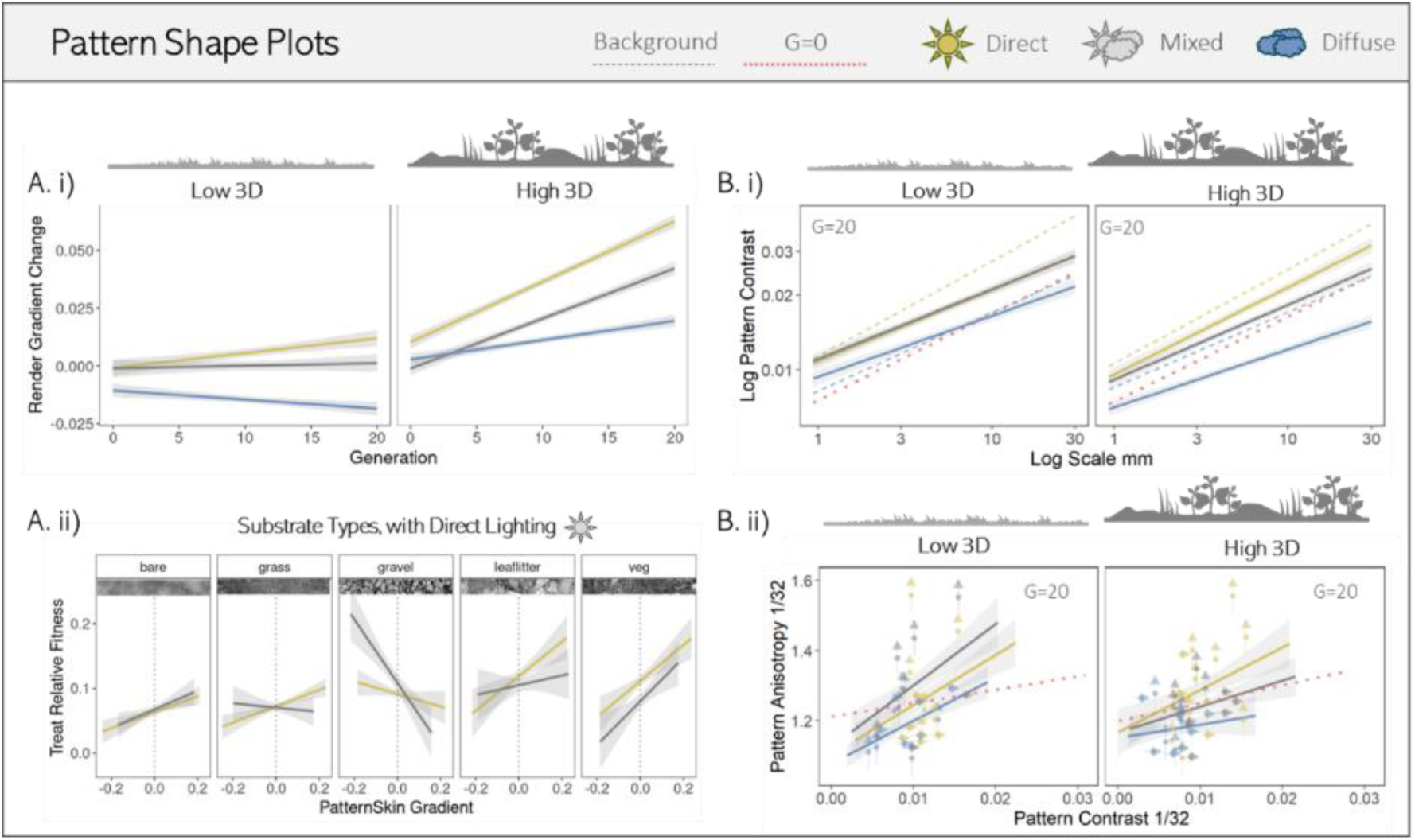
Phenotypic change in countershading and pattern shape. A.i) Targets under direct and mixed lighting became increasingly positive in their luminance gradient. A.ii) Having a more positive render target gradient increases survival time for all substrates barring gravel when the background has direct lighting, for both mixed and direct only conditions B.i) Targets evolved under direct and mixed lighting conditions evolve a greater pattern contrast at larger spatial scales and across all scales when background 3D complexity is hgih. Red dotted line shows the targets in generation zero, dashed lines show the pattern energy of the diffuse and direct backgrounds. B.ii) Under direct lighting contrasting patterns are more directional and orient vertically against 3D complex backgrounds, arrows indicate whether the population average orientation is vertical or horizontal. Habitat 3D variation is split by the quantiles for the habitats’ mean depth variation across spatial scales.

**Table 2.** Linear mixed model output effects of dominant substrate on the relationship between countershading gradient of the target-skin (CS) and substrate type on target survival time. Positive estimates and t values are shown in blue while negative values are shown in red. Model formula:

| | Predictor | $\beta$ | SE | DF | t value | p value |
| --- | --- | --- | --- | --- | --- | --- |
| Fitness | Intercept | 0.462 | 0.02 | 25 | 30.54 | <0.001 |
|  | CS_Gradient:bare | 0.275 | 0.05 | 19120 | 5.46 | <0.001 |
|  | CS_Gradient:grass | 0.367 | 0.05 | 18930 | 7.51 | <0.001 |
|  | CS_Gradient:gravel | -0.122 | 0.06 | 19140 | -1.92 | 0.054 |
|  | CS_Gradient:leaflitter | 0.174 | 0.06 | 19140 | 2.79 | 0.005 |
|  | CS_Gradient:veg | 0.345 | 0.06 | 19100 | 5.73 | <0.001 |

Compared with diffuse lighting treatments, targets under direct and mixed lighting evolved greater pattern contrasts then diffuse at larger spatial scales, especially when 3D complexity was high, but less so for mixed populations (Figure 4, Table 3). At the scale where the maculation genes for the target patterns best predicted pattern contrast for the unrendered target (1/32), targets became more directional in their patterning for direct and mixed lighting (N-Gen, β = 0.023, t_36210_ = 2.81, p=0.005; N-Gen * MIXED, β = -0.004, t_36210_ = -0.30, p=0.762) but less directional for diffuse (N-Gen * DIFFUSE, β = -0.126, t_36210_ = -10.46). Directionality further increased with number of generations when 3D complexity was high, but only for direct lighting populations (N-Gen * 3D, β = 0.095, t_36210_ = 11.10) and otherwise declined for mixed and diffuse populations (N-Gen * 3D * MIXED, β = -0.176, t_36210_ = -14.35; N-Gen * 3D * DIFFUSE, β = -0.126, t_36210_ = -10.26), supporting the hypothesis that direct lighting selects for more directional patterns then diffuse lighting. Patterns also became increasingly vertical for direct (N-Gen, β = 0.043_36210_, t = 5.27) compared to mixed and diffuse (N-Gen * MIXED, β = -0.031, t_36210_ = -2.58; N-Gen * DIFFUSE, β = -0.073, t_36210_ = -5.94). As with directionality, this observed increase in verticality was enhanced for 3D complex habitats for direct but not diffuse or mixed lighting populations (N-Gen * 3D_variation_, β = 0.095, t_36210_ = 11.10; N-Gen * MIXED * 3D_variation_, β = -0.176, t_36210_ = -14.35; N-Gen * DIFFUSE * 3D_variation_, β = -0.126, t_36210_ = -10.26).

**Table 3.** Linear mixed model output effects of lighting treatment and 3D complexity on the unrendered target-skin’s contrast across spatial scales. Positive estimates and t values are shown in blue while negative values are shown in red. Model formula lmm (PatternEnergy ∼ Scale*Light*3D+(1|Treatment)+(1|Habitat) +(1|UniquePhoto))…

| Log Pattern Energy | Predictor | $\beta$ | SE | DF | t value | p value |
| --- | --- | --- | --- | --- | --- | --- |
|  | Intercept | -0.766 | 0.13 | 76 | -5.84 | <0.001 |
|  | Scale | 0.257 | 0.01 | 9040 | 23.33 | <0.001 |
|  | Mixed | 0.029 | 0.17 | 56 | 0.18 | 0.861 |
|  | Diffuse | -0.085 | 0.17 | 62 | -0.52 | 0.607 |
|  | Mean 3D Var | -0.169 | 0.18 | 75 | -0.96 | 0.341 |
|  | Scale : Mixed | -0.002 | 0.02 | 9040 | -0.15 | 0.885 |
|  | Scale : Diffuse | -0.033 | 0.02 | 9040 | -2.19 | 0.029 |
|  | Scale : 3D | 0.055 | 0.01 | 9040 | 3.73 | <0.001 |
|  | Mixed : 3D | 0.010 | 0.22 | 55 | 0.04 | 0.965 |
|  | Diffuse : 3D | 0.026 | 0.23 | 60 | 0.11 | 0.910 |
|  | Scale : Mixed : 3D | -0.053 | 0.02 | 9040 | -2.50 | 0.012 |
|  | Scale : Diffuse : 3D | -0.111 | 0.02 | 9040 | -5.24 | <0.001 |

### 4. Additional Features: Edge enhancement and glossiness

The positive shift in luminance contrast from edge enhancement increased with n-generations (N-Gen, β = 0.043, t_36210_ = 5.31) but less so for diffuse lighting targets (DIFFUSE * N-Gen, β = -0.030, t_36210_ = -2.54, p = 0.011). Increased 3D complexity also resulted in increased edge enhancement (N-Gen * 3D_variation_, β = 0.027, t_36210_ = 3.283.10, p = 0.001), though unexpectedly this observed increase was greater for mixed lighting treatments for 3D variable habitats (N-Gen * MIXED * 3D_variation_, β = 0.077, t_36210_ = 6.59) and irrespective of the 3D environment (N-Gen * MIXED, β = 0.024, t_36210_ = 2.09, p=0.037). Whether the increased selection for edge enhancement within 3D complex mixed lighting regimes was due to it functioning as a generalist strategy or 3D environments increasing the survival time under direct lighting is difficult to disentangle.

The ‘potential’ glossiness (change in target luminance value when gloss genes are maximised) was negatively affected by direct lighting and the target-skin’s luminance, darker targets inherently appear glossier (See Supplementary Material). The relative glossiness of the evolved targets was significantly lower for 3D complex habitats except for those evolved under mixed lighting conditions (Table 4). Targets evolved significantly less-glossy phenotypes overtime but only within 3D complex backgrounds. Notably, diffuse lighting increased selection against glossiness while mixed lighting did not select against glossiness. Some level of glossiness may help compensate for changes in target and background appearance under variable lighting conditions. Most surviving targets, however, did not appear ‘glossy’, with extreme glossiness values dying out early and high internal contrast from patterning masking the specular highlights from gloss

**Table 4.**
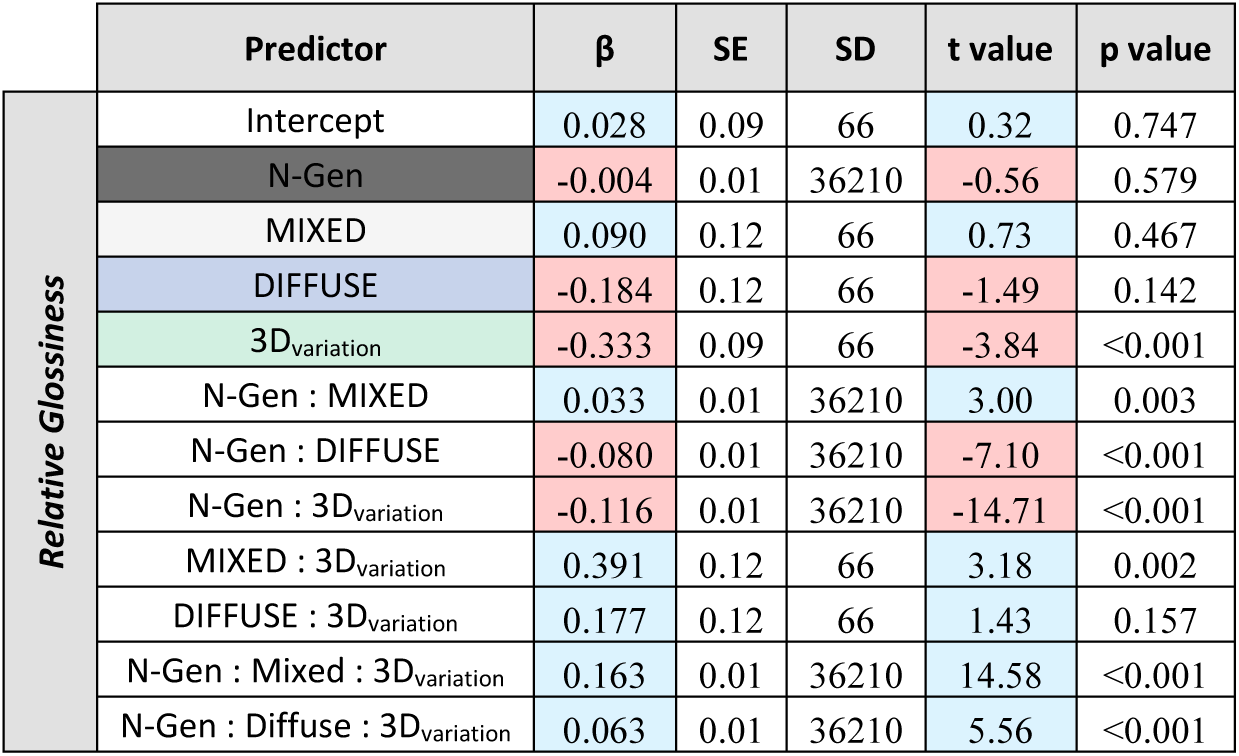
Linear mixed model output effects of lighting treatment and background geometry on the evolved glossiness of the targets, relative to the maximum glossiness value they could have. Model formula lmm (RelativeGlossiness ∼ N-Gen * Light * 3D_variation_ + (1|Treatment) + (1|Habitat))…

| | Predictor | $\beta$ | SE | SD | t value | p value |
| --- | --- | --- | --- | --- | --- | --- |
| Relative Glossiness | Intercept | 0.028 | 0.09 | 66 | 0.32 | 0.747 |
|  | N-Gen | -0.004 | 0.01 | 36210 | -0.56 | 0.579 |
|  | MIXED | 0.090 | 0.12 | 66 | 0.73 | 0.467 |
|  | DIFFUSE | -0.184 | 0.12 | 66 | -1.49 | 0.142 |
|  | 3D<sub>variation</sub> | -0.333 | 0.09 | 66 | -3.84 | <0.001 |
|  | N-Gen : MIXED | 0.033 | 0.01 | 36210 | 3.00 | 0.003 |
|  | N-Gen : DIFFUSE | -0.080 | 0.01 | 36210 | -7.10 | <0.001 |
|  | N-Gen : 3D<sub>variation</sub> | -0.116 | 0.01 | 36210 | -14.71 | <0.001 |
|  | MIXED : 3D<sub>variation</sub> | 0.391 | 0.12 | 66 | 3.18 | 0.002 |
|  | DIFFUSE : 3D<sub>variation</sub> | 0.177 | 0.12 | 66 | 1.43 | 0.157 |
|  | N-Gen : Mixed : 3D<sub>variation</sub> | 0.163 | 0.01 | 36210 | 14.58 | <0.001 |
|  | N-Gen : Diffuse : 3D<sub>variation</sub> | 0.063 | 0.01 | 36210 | 5.56 | <0.001 |

## Discussion

We assessed the effects of three different lighting treatments on the evolution of artificial animal colouration. In doing so we confirmed that direct lighting on average increased rather than decreased the difficulty of target detection due to higher background complexity and in doing so facilitated faster and more reliable evolution (fewer failures to improve the fitness proxy, survival time) of camouflage. This provides the first empirical demonstration of Merilaita’s oft-cited model for the importance of background complexity in the evolution of camouflage and shows shadows to be a powerful source of complexity (Merilaita 2003). In line with our predictions, there were clear differences in the colouration, contrast and orientations of evolved target patterns between lighting treatments. Specifically, selection under direct lighting leads to more contrasting, darker-bluer patterns, countershading, increased edge enhancement and more directional patterns compared to diffuse lighting environments. These phenotypic features emerged in just 20 generations, demonstrating how alternate camouflage strategies are selected for rapidly under different habitat structures and lighting regimes. The phenotypic differences between evolved populations also corresponded with differences in the predictive power of common camouflage metrics, with edge disruption being more effective against backgrounds with direct lighting.

Our results highlight the critical interactions between lighting and habitat geometry as most of the relationships between lighting and camouflage were influenced by geometry. Habitat geometries that produced larger shadows in the presence of direct lighting altered the evolved target patterns, increasing pattern contrast and directionality for both mixed and direct-lighting-only treatments. The increased scale of background spatio-chromatic variation under direct lighting against more 3D complex habitats increased the survival time benefits of camouflage under direct relative to diffuse lighting. This increased disparity between the lighting conditions may have driven the observed increased overlap in phenotype between the mixed and direct-only lighting treatments within more 3D complex habitats. As targets within the mixed lighting treatments were randomly assigned to either a diffuse or direct background, habitats where one lighting condition favours survival over another will have skewed selection of traits to the lighting with the highest benefit to survival (Penacchio et al. 2018; Hughes et al. 2023). Some components of the mixed lighting evolved targets did differ from the direct within the more 3D complex habitats. For example, pattern contrast at larger spatial scales evolved to be more intermediate between direct and diffuse. Mixed lighting targets also continued to have low levels of glossiness in later generations despite diffuse lighting and 3D complexity selecting against glossiness for the other treatments. As the luminance of glossy surfaces changes in response to both lighting and geometry, glossiness may serve as a mechanism of maintaining contrast match across different environments. The overall selection against glossiness however was unsurprising, with previous experiments, both military and ecological, showing gloss to negatively impact survival (Thomas et al. 2023).

Countershading has long been proposed as an adaptation for camouflage by improving background match and/or by destroying shape-from-shading cues generated by the interaction of the illuminant with geometry (Rowland 2009; Penacchio et al. 2015). We add to the mounting body of evidence that countershading does indeed improve camouflage under direct lighting and that, although mixed lighting selected for reduced countershading, some level of shading gradient persisted, indicating that it can be robust in the face of variable lighting conditions (Penacchio et al. 2018). We also found correlations between countershading and striped patterns as well as reduced internal contrast under direct lighting for darker targets irrespective of countershading. Phylogenetic comparisons of ungulate patterns also found a correlation between more intense countershading and stripes (Allen et al. 2012), suggesting that these two features may interact to disrupt shape.

We also found exceptions to the prediction that the gradient of countershading should perfectly counter self-shading, known as ‘optimal countershading’ (Cuthill et al. 2016). Against backgrounds where the substrate contained objects that resembled the target, e.g., pebbles and stones, countershading was not selected for and in some cases the inverse evolved, despite the intensity of self-shading being higher within these habitats. The presence of objects that match geometric features of the target could negate the need for self-shadow destruction and by extension make masquerade conflict with countershading as a camouflage strategy (Troscianko et al. 2009). Indeed, many of the targets observed possessed patterns or colourations that resembled one singular object, particularly stones. Patterns which instead distort the shape of the self-shadow such as inverse countershading or shadow-like maculation could provide alternative mechanisms for masking shape recognition (Kelley et al. 2022). However, it is worth noting that our experiment only considers prey viewed from one orientation in plain view, whereas countershading must function against more than one detection angle and in the presence or absence of obstruction. It is possible that countershading is still beneficial when viewed from the side where the background appearance differs in visible structure. Field experiments are required to further validate the phenotypic effects found within our experiments, both for countershading and other patterning. Likewise, the effect of body shape and the shape of background distractors on selection for countershading has yest to be explored.

Many animals have directional striped maculation which can be orientated to match different background features (Kang et al. 2012). Animals which occupy mixed lighting conditions may need to change their orientation and position as clouds pass over, not just for thermoregulation, but to maintain orientation match to the pictorial features of their background (Penacchio et al. 2015; Mavrovouna et al. 2021). The orientation of maculation may reflect the typical orientation of the animal relative to the light when basking or navigating its environment, in a similar way to how orientation affects countershading in arboreal animals (Angilletta 2009; Kamilar and Bradley 2011; Mavrovouna et al. 2021). The increased level of pattern directionality and shifts in pattern orientation observed between our lighting treatments may have been the result of how we orientated the targets. Targets were only orientated with respect to background features for backgrounds with direct lighting. The targets were always aligned parallel to their self-shading gradient so that their countershading patterns would in turn be align with them. As a result, targets against direct backgrounds could evolve patterns parallel to the light, matching the orientation of vertical received shadows from vegetation found within 3D complex habitats. Meanwhile, for diffuse backgrounds, the target orientation was random with respect to the background patterns. Other aspects of pattern shape and orientation may have also differed if the targets were allowed to make behavioural decisions on where to position and orientate themselves based on the global and/or local statistics of the scene, e.g. moths that align with the patterns of background bark (Kang et al. 2012; Stevens and Ruxton 2019). These behavioural decisions are likely to have large impact on camouflage phenotypes (Kang et al., 2012; Stevens and Ruxton, 2019).

Overall, our work is the most comprehensive endeavour to link animal camouflage to the physical structure and lighting regimes of habitats. We have been able to show that while there are trends for optimising phenotypes under different conditions, there are also numerous exceptions and alternative strategies that provided effective protection by camouflage. These exceptions potentially act as important drivers of evolutionary diversity, particularly when linked with predator learning and frequency-dependent effects. Several camouflage adaptations such as masquerade, disruptive colouration, countershading and even colour match were affected by geometry and lighting environment in ways that were unexpected from previous work (Cuthill et al. 2005, 2016; Skelhorn et al. 2010). Patterns that are more contrasting than the background and superficially disruptive can also be pattern matching to the background contrast under direct lighting, as shown by the increased pattern contrast of mixed and direct lighting only evolved patterns. Simple ecological and behavioural differences between species in how they interact with their background’s geometry under different lighting conditions can, by extension, affect the mechanisms with which they camouflage themselves. Furthermore, any alterations to habitat geometry and lighting from human activity, whether it be shifts in atmospheric condition from climate change or habitat alteration from land-management and urbanisation, could affect the adaptive value of camouflaged phenotypes and the predation risk of vulnerable species. Finally, the methods used here can also be used to explore the effects of and interactions between lighting, geometry and a wide range of alternative drivers of colour evolution, such as phylogenetic constraints, object shape, thermoregulation, signalling and observer condition (visual system and field of view) (Stuart-Fox et al. 2008; Stuart-Fox and Moussalli 2009; Smith et al. 2016; Cheng et al. 2018; Van Den Berg et al. 2020).

## Author Contributions

The initial concept for assessing the interactions between geometry and lighting environment on camouflage was conceived by JT and added to by IC and GRAH. Colour-calibrated photographs and 3D scans were collected by GRAH. The code for running the online experiment paired with the CamoEvo toolbox was created by GRAH. The original computer game scripts were derived from previous games created by JT. The manuscript was first written by GRAH, with subsequent edits by all authors. GRAH performed all image and data analyses with guidance from JT and IC.

## Supporting information

Supplemental Text B - Stats Tables

## Acknowledgements.

Natural Environment Research Council (NERC) GW4+ NE/S007504/1 funded GRAH in a CASE partnership with the Game and Wildlife Conservancy Trust, UK. JT was funded by a NERC Independent Research Fellowship NE/P018084/1 and ICC by Biotechnology and Biological Research Council grant BB/S00873X/1.

