## Supplemental Text A - Additional Info for "Shining a light on camouflage evolution: using genetic algorithms to determine the effects of geometry and lighting on optimal camouflage"

#### Experiment Setup

##### *CamoEvo Settings*

The work presented in this research predominantly relies on the tools made available from the CamoEvo toolbox. All target generation was carried out using CamoEvo, while the psychophysics game was replaced with a custom online game (Hancock and Troscianko, 2022).

##### *ImageGA*

For CamoEvo's genetic algorithm, the following settings were used:

Sex = **false,**
Population Size =**24**
Starting Gene Distribution = **Uniform**
Deletion Pool = **66.6%**
Breeding Pool = **33.3%**
Point Mutation Prob = **0.005**
Lvl1 Mutation Prob = **0.001**
Lvl2 Mutation Prob = **0.001**
Link Mutation Prob = **0.3333**
Rescue Number = **6**
Gene Displace Prob = **0.01**
Adaptive Mutations = **False**
Crossover Type = **Multi_Point**
Crossover Probability = **1.00**
Recombination = **incomplete**
Mating System = **random**
Breeding Pool Assignment = **ranked**

Sex was disabled, meaning that any individual could mate with any individual in the breeding pool. The population size was set to 24 individuals with a uniform distribution for the starting genes as opposed to a random or normal distribution. 2/3 of the population was deleted each generation with 1/3 surviving and reproducing. The probability of a point mutation occurring for each gene was 0.005, for a lvl 1 group mutation 0.001, and a lvl 2 group mutation 0.002 (see ImageGA handbook for more details). The probability of a group mutation affecting other members of the group was 0.333. The top 6 individuals in a given generation would be protected from a deletion in the next generation to reduce the effect of noise on deletion. Genes with similar values had a 0.01% chance of being displaced during crossover increasing diversity. Crossover could have multiple breakpoints of random length, as opposed to pure random, one-point (single break) or two-point (two breaks) crossover.

Recombination was incomplete meaning that genes were combined with a weighted average simulating recombination for polygenic inheritance. Individuals within the breeding population were randomly paired and were assigned by rank.

##### *Colour Space*

Targets prior to being mapped onto the grey target had a colour space of L* 0 to100, a* -50 to 50 and b* -15 to 80 (Figure SA1).

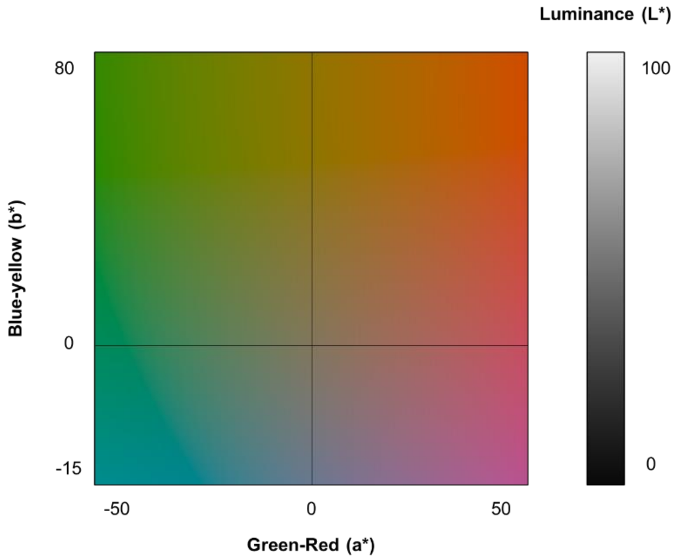


Figure SA1, Colour space range in CIELAB for the un-rendered targets. This map represents the full range of values that the targets could use for the top colour, bottom colour and maculation.

##### *Spherical Deformation*

Prior to saving the output from the CamoEvo pattern generator a spherical deformation function was applied using a custom Java script for ImageJ (Schneider et al., 2012). This function wrapped the CamoEvo pattern to match a 3D sphere, ensuring that the target patterns matched the geometric properties of spherical targets. Without the deformation the targets would appear flatter, lacking the additional cues of shape that occur from changes in the orientation and scale of patterns (Figure SA2).

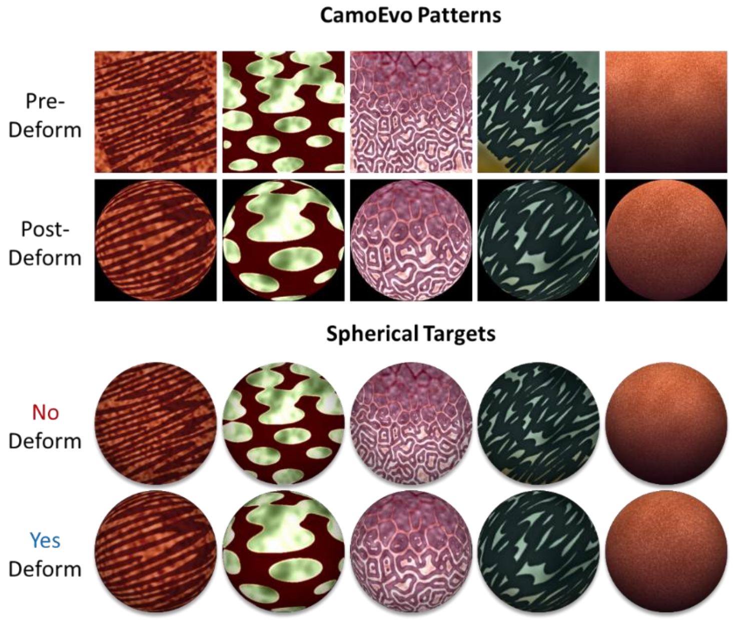


Figure SA2, Showcase of spherical deformation. Above shows 5 different randomly generated CamoEvo patterns before and after the spherical deformation has been applied. Below shows those same patterns applied to a spherical target. Spherical deformation increases the scale of pattern features closer to the centre of the image.

---

##### *Glossiness*

To allow for glossiness to evolve two new decimal genes were added to CamoEvo's animal pattern generator gls_lvl_bkg and gls_lvl_mac (Hancock and Troscianko, 2022). These genes regulated the level of glossiness of the target's background colour and maculation respectively, the maculation being the un-gradiented paattern. Gloss was applied by combining the images for the matt grey target and glossy target using addition (Figure SA3 and Figure SA4). By separating the glossiness of the background and maculation the material properties of the target could vary as would be expected for changes in the material properties of pigments and structural colours, although independent from any real world correlations between chroma and gloss (Cuthill et al., 2017; Shawkey and D'Alba, 2017). Genes controlled glossiness by manipulating the opacity of the glossy target, a value of 0 = 0% opacity and a value of 1 = 100% opacity, with the following formula: opacity = sqr(gene) * 100.

---

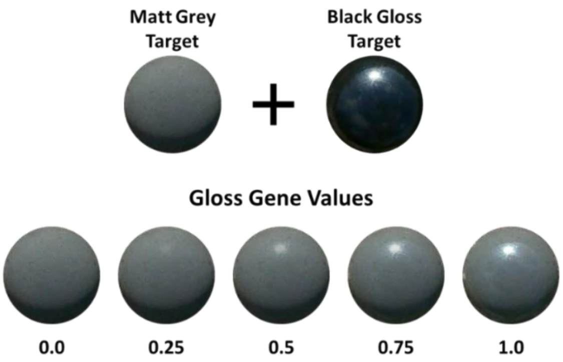


Figure SA3, showcase of how glossiness was added to the targets. Above shows the raw grey target and glossy target. Below shows a sequence of increasing glossiness from left-right with the decimal gene value for glossiness shown below.

---

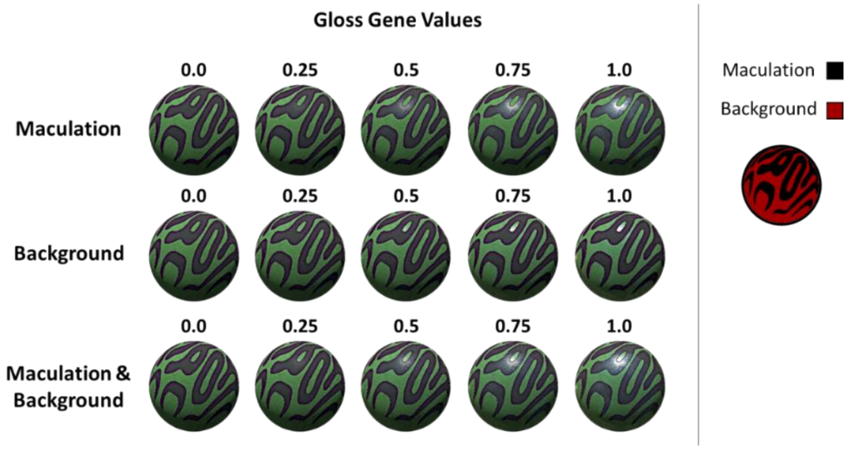


Figure SA4, showcase of how glossiness was divided between the maculation and background using the two different genes. The maculation area is highlighted in black, and the background area is highlighted in red. Targets are shown for a background image with direct lighting.

---

#### *Online Game*

##### *Background Assignment*

Populations were assigned to one of the three lighting treatments, direct, mixed and diffuse lighting and then one of the 28 different habitat types. Each habitat had a total of 48 images, 24 direct and 24 diffuse. For each generation of the game all 24 targets were randomly paired with one of the background images for their select habitats. Direct lighting treatments were only rendered on direct lighting backgrounds, diffuse lighting treatments were only rendered on diffuse, meanwhile mixed lighting were assigned to either direct or diffuse with 50% of each generation being direct and the other 50% being diffuse.

##### *Target Rendering*

The CamoEvo target patterns (target-skin) were applied to the scene dividing the grey target by the reflectance value of the sphere as a 32-bit RGB stack in ImageJ and then multiplying by the target by the RGB values of the CamoEvo animal pattern (Figure SA5). Prior to multiplication the pattern was rotated to match the direction of the targets self-shading gradient providing that the target was under direct lighting and the self-shading gradient was visible. To add glossiness using the methods outlined above, the corresponding glossiness photograph was added to the target after the CamoEvo pattern was rendered.

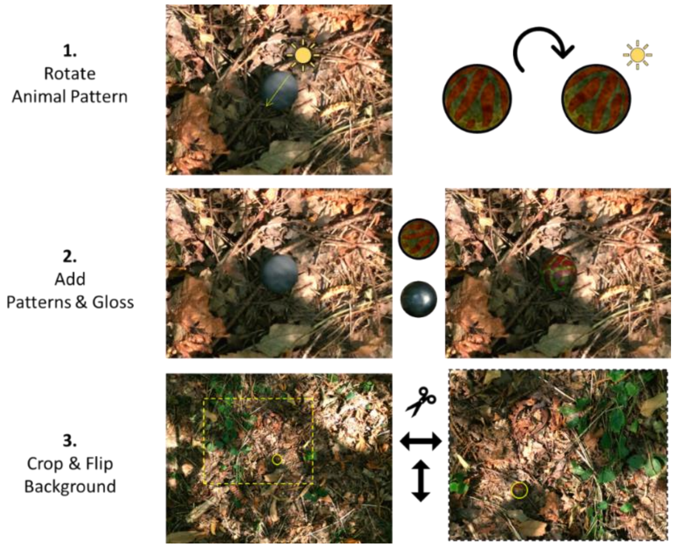


Figure SA5, Steps for creating the rendered scenes. Firstly, the animal pattern is rotated to face the orientation of the light if the background is under direct lighting. Secondly, the target pattern and glossiness levels are rendered to the scene using multiplication and addition respectively. Thirdly, the background is cropped and then randomly flipped with a 50% chance of being flipped horizontally and or vertically.

##### *Background Cropping*

Once the target was rendered the background image, the image was rescaled so that the target diameter was 80 px and then cropped to make the scenes for the game. To make the cropped scenes 1904 × 1488 rectangle was positioned at the centre of the target and then randomly offset such that the target could not be within 2x the target diameter (160px) of the edge or centre of the screen (Figure SA5 & Figure SA6). The scene was then randomly flipped vertically (50%) and/or horizontally (50%) before being saved along with the coordinates X and Y coordinates of the target and the background used. These crops were used to make the 24 scenes corresponding to the 24 individuals in the population. All scenes were assigned a random order in a sequence of 1-24. In addition to the 24 targets within the population, a demo target was randomly assigned to one of the backgrounds. The background used was always one of the 4 closest targets to the middle of the display sequence (11-14), this was to prevent the user from facing the same background back-to-back and to reduce the likelihood of memorising the backgrounds appearance. The demo target was always a randomly selected dead individual from the previous generation, except in the first generation where it was a randomly generated target.

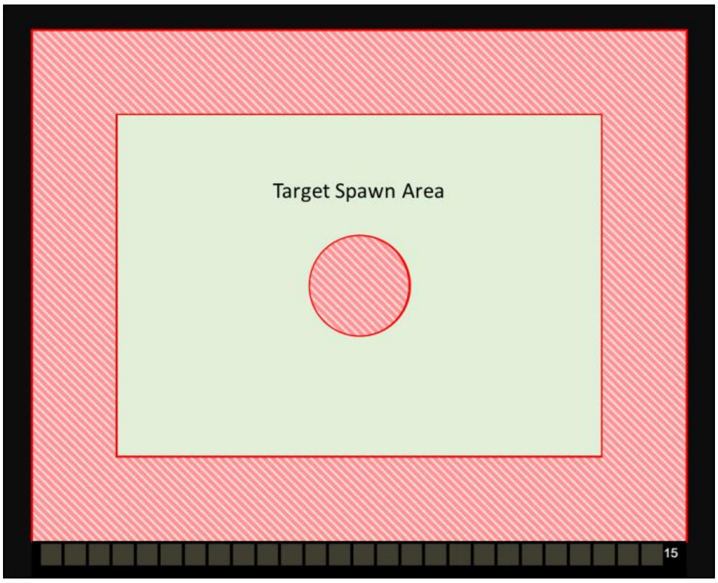


Figure SA6, Showcase of viable target screen locations. Targets could only spawn in the green regions.

##### *Server Hosting*

All background images were saved in a folder labelled “Combined_GenPat_[X]”, where X = the current generation. Images were saved alongside a tab-delimited text file recording the crop coordinates, target location and which background and habitat the files belong. All populations were rendered to scenes using a batch script in ImageJ which would loop through all populations until they had been created for the current generation in CamoEvo. The status of each population was recorded in a separate table listing whether or not the population had been **rendered**, **uploaded**, **played, downloaded** and **evolved**. To manage these phases, 4 instances of ImageJ were run simultaneously. An easy method of doing this is to create multiple copies of the .exe file in Windows within the same folder e.g. Image_J_1.exe, Image_J_2.exe, etc, select them all, and then press run. These instances were managed as follows: *1. Creation* in charge of generating and rendering the evolved targets, *2. Sending* in charge of uploading the targets to the server, *3. Returning* in charge of downloading the data from the online game after the population had been played, and *4. Algorithm* in charge of running the genetic algorithm to create the genes for the new population.

Once a population was rendered, the “Combined_GenPat_[X]” folder was uploaded to an SFTP server hosted by the [Exeter Visual Ecology](https://www.visual-ecology.com/) group (https://www.visual-ecology.com/). This server has been used to host a variety of online virtual camouflage experiments. To allow ImageJ to copy files to the server the program WebDrive [WebDrive, South River Technologies; T1910 Towne Centre Blvd #250, Annapolis, MD 21401, United States] was purchased and used to mount the server to a file directory.

Sending images to the server presented many automation challenges, if the server went offline for any reason ImageJ would still recognise the directory as being present and would error/crash when attempting to upload files to the folder, thus ending the batch loop with a crash. To solve this issue a detection mechanism was developed by exploiting error messages. Even though ImageJ can’t typically upload files to an SFTP server without WebDrive, or a similar system, it is capable of downloading SFTP files providing you give it the right login details. Sending a URL request without the details creates an error stating that the password is wrong, but if the server is offline the error message changes:

```
OnlineCheck1=File.openUrlAsString("ftp://ftp.visual-ecology.com/camoevo/GenPathGate.txt");
Online, error = < Error: sun.net.ftp.FtpLoginException: Invalid username/password >
Offline, error = <Error: java.net.NoRouteToHostException: No route to host: connect>
```

Exploiting these error messages allowed for a gated while loop to be made.

```
OnlineCheck1=File.openUrlAsString("ftp://ftp.visual-ecology.com/camoevo/GenPathGate.txt");
	if(startsWith(OnlineCheck1, "<Error: sun.net.ftp.FtpLoginException: Invalid
	username/password>")==1){
	selectWindow("Gate Status");
	Table.setColumn("Connection",newArray("Online"));
	OnlineCheck2=1;
	}else{
	selectWindow("Gate Status");
	Table.setColumn("Connection",newArray("OFFLINE!"));
	OnlineCheck2=0;
	}
```

While the server was offline (Error != Invalid username/password) ImageJ would loop until it came online again before uploading the images. Once all the images were uploaded, the status for the population would be changed to uploaded and the ImageJ instances and the *2. Sending* instances of ImageJ would move on to the next un-uploaded population or wait until there were new populations to upload.

*Gameplay*

Within the SFTP server, a separate table was used to list all the population directories and their status as played or un-played. When a player launched the game, they would be randomly assigned one of the playable populations. A population was playable if it had not been played already and if it was from the youngest generation. For example, let us say that of the 84 populations, 10 of them are in Generation 5 while 74 of them are in Generation 6. A player could only play 1 of the 10 populations in Generation 5. If all populations had been played but the next generation was yet to be uploaded, players would be randomly assigned a population and the data would be unused. Once a player started a game, they were shown 26 slides. The first and last slide used the demo image, while the middle 24 used the randomly assigned order of target treatments for the population. The demo target on the final slide was also used as the submit button to end the game. Between each slide, the player was tasked with moving their cursor to the centre of the screen and then not moving until they saw the target. Once within the ring it ring would fill with white over a period of 1.5 seconds and then start the next slide (See Figure SA7). The motion of the cursor from the centre was detected by using a smaller ring equal in diameter to ½ the target diameter (40px). As with the CamoEvo psychophysics game (Hancock and Troscianko, 2022), reaction time was recorded as the time taken to leave the ring and capture time was recorded as the time taken to click on the target area (circle 1.5x target diameter) (See Figure SA7) .  Survival time was calculated as reaction time, unless capture time was >600ms then reaction time as leaving the centre in this instance was presumed a mistake. Once the population was played its results for survival time were downloaded and used as the fitness variable for the genetic algorithm to create a new population.

---

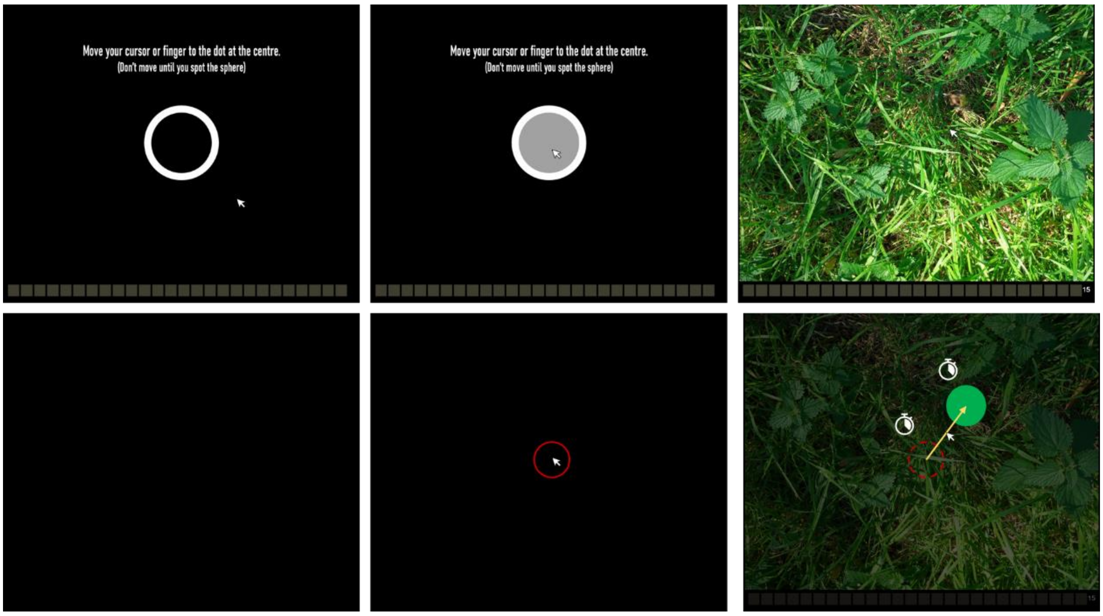

Figure SA7, Target survival time recording. The top row shows what the player sees when they move their mouse to the centre and play each slide. The bottom row shows the reaction time and capture circles, once the mouse leaves the red circle reaction time is marked and capture time and the end of the slide occurs when the target circle is clicked.

#### Measures

##### *Background Measures*

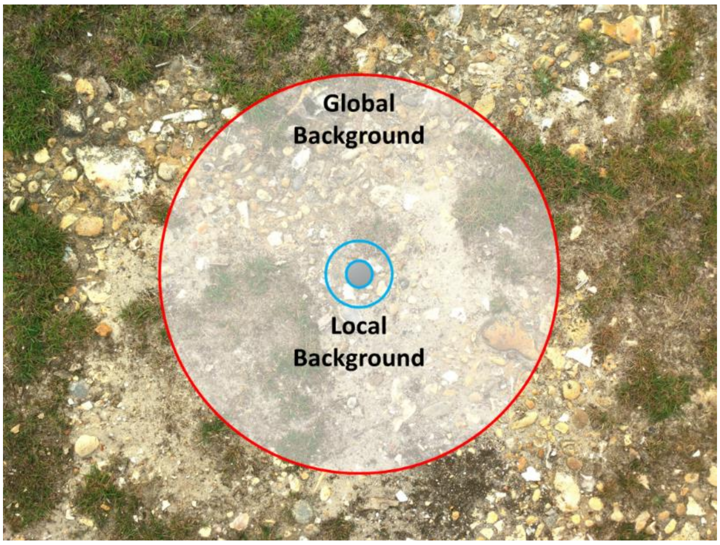


Figure SA8, Illustration of the measurement areas for the local (2.5x target scale, blue) and global background (15x target scale, red).

##### *Countershading*

For each habitat the grey spheres were oriented so that light came from above averaged across all 24 backgrounds for each lighting condition (direct and diffuse). This resulted in 2 averaged grey targets, a direct average and diffuse average. Luminance gradients were measured as the linear gradient of luminance values from the top-bottom of the target (Figure SA9 A positive gradient indicated a dark-light dorsal-ventral gradient while a negative gradient indicated a light-dark dorsal-ventral gradient. Without countershading the rendered targets would have a negative gradient similar to that of the grey targets. Luminance gradients were measured for the averaged grey target, the un-rendered CamoEvo targets and the targets rendered on the direct and diffuse averaged grey target. Countershading was given as the positive change in the luminance gradient. The greater the positive gradient the more effective the countershading (Figure SA10). Reverse countershading would result in a more negative gradient.

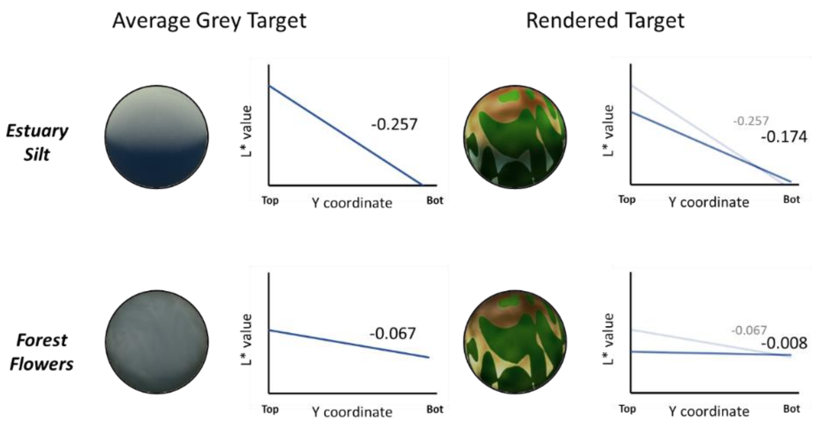


f

Figure SA9, Example difference in luminance gradient between the average grey target and target with a rendered pattern. Each row shows the averaged grey target, the rendered target and their respective luminance slopes and linear gradient values. Above shows the values if the target is presented on an open silt estuary habitat and below shows a closed forest flower habitat. Illustration of the measurement areas for the local (2.5x target scale, blue) and global background (15x target scale, red).

---

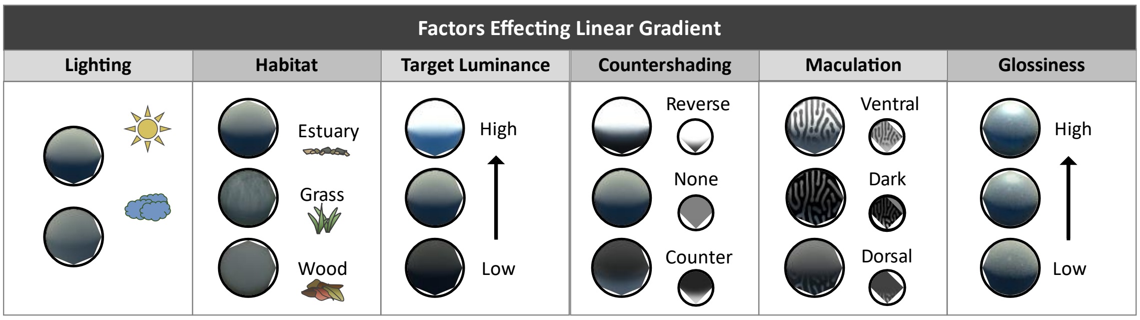


Figure SA10, Different mechanisms that altered the dorsal-ventral luminance gradient of the targets. Steeper negative (light-dark) slopes are shown above. From left-right. Direct lighting increases the intensity of the luminance gradient. Habitats that are more open have a more intense gradient. Lighter targets have a greater luminance contrast while darker targets have a lower contrast. Reverse counter shaded targets have a greater luminance contrast (light-dark) while countershaded (dark-light) have a lower contrast. The distribution of light and dark maculation can shift the luminance gradient. Lastly specular highlights increase the brightness of the surface facing the light and so increase the intensity of the dorsal-ventral gradient.

---

##### *Glossiness*

Just as countershading compared the luminance gradient of the rendered and un-rendered target, to measure glossiness the mean and standard deviation (contrast) of luminance (L*) were measured and compared for the target without gloss (genes = 0), with the level of gloss evolved (genes = x) and the maximum gloss (genes = 1) (Figure SA11) The effect of the glossiness genes on glossiness varied depending on multiple features of both the target and habitat (Figure SA12).

---

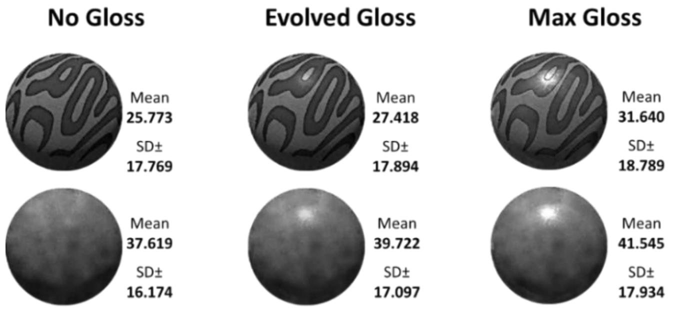


Figure SA11, Example changes in luminance gradient from rendering of the target. Each row shows the averaged grey target, the rendered target and their respective luminance slopes and linear gradient values. Above shows the values if the target is patterned and below shows the values for an un-patterned target.

---

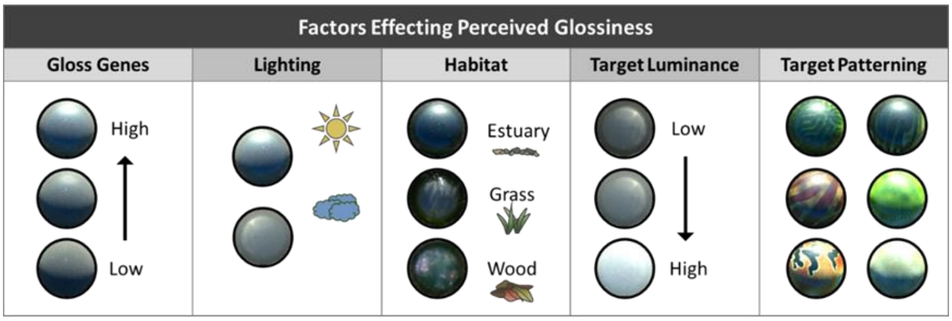


Figure SA12, Different target and background features that altered the glossiness of the targets. From left-right. Increased gloss value for the genes increases glossiness, direct lighting increases the deviation in luminance from gloss, more open habitats have more obvious specular highlights, the lighter the target the lower the luminance increase and contrast difference from gloss and target patterning can mask the appearance of gloss.

---

### Countershading

#### *Luminance and Self-shading*

Countershading was not the only mechanism with which targets could reduce their internal luminance gradient. The darker the target was the lower the luminance gradient (Figure SA13) However, this gradient shifted to be less negative with the number of generations confirming that other aspects of the target’s patterning influenced self-shadow contrast, such as countershading (See Table SA1). This shift was greatest for the direct lighting populations and was negative for the diffuse, meaning diffuse populations became less countershaded.

---

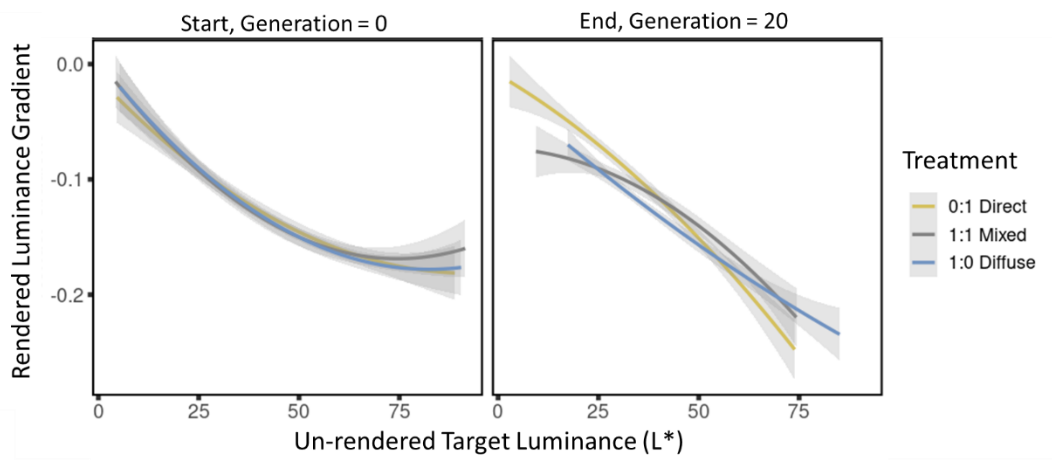


Figure SA13, Comparison of relationship between the luminance (L*) of the un-rendered target-skin and the luminance gradient of the target rendered against the average grey sphere for its respective habitat under direct lighting.

---

Table SA1, Stats output for the effect of the target-skins luminance (L*) on the rendered luminance gradient of the target. Linear mixed model formula: lmer(LuminanceGradient ~ Generation * TargetLuminance * Light + (1|Habitat) + (1|Population) …

| | Predictor | β | SE | DF | t | p value |
|---|---|---|---|---|---|---|
| *Render Luminance Gradient* | Intercept | -0.116 | 0.01 | 31 | -12.56 | <0.001 |
| | Generation | 0.002 | 0.00 | 36240 | 4.22 | <0.001 |
| | Skin Luminance | -0.040 | 0.00 | 34560 | -68.83 | <0.001 |
| | Mixed | -0.002 | 0.00 | 42 | -0.36 | 0.723 |
| | Diffuse | -0.009 | 0.00 | 43 | -2.00 | 0.052 |
| | Gen : Lum | -0.006 | 0.00 | 36090 | -13.52 | <0.001 |
| | Gen : Mixed | -0.003 | 0.00 | 36240 | -4.51 | <0.001 |
| | Gen : Diffuse | -0.004 | 0.00 | 36230 | -6.63 | <0.001 |
| | Lum: Mixed | 0.004 | 0.00 | 34250 | 4.47 | <0.001 |
| | Lum : Diffuse | 0.006 | 0.00 | 32790 | 7.05 | <0.001 |
| | Gen : Lum : Mixed | 0.002 | 0.00 | 36000 | 3.87 | <0.001 |
| | Gen : Lum : Diffuse | 0.004 | 0.00 | 35840 | 6.84 | <0.001 |

---

#### *Internal Gradient and Contrast Match*

One of the leading hypotheses for why countershading evolves is to reduce the contrast difference between the animal and its local background (Donohue et al., 2020; Ruxton et al., 2004). To test whether countershading decreases contrast, the contrast difference of the rendered target against the background was modelled with the target-skin, n-generations and lighting treatment as predictor variables. Unsurprisingly having a more positive dorsal-ventral luminance gradient reduced the contrast difference between the target and the background under direct lighting, while a more negative gradient increases the difference (Figure SA14, Table SA2).

---

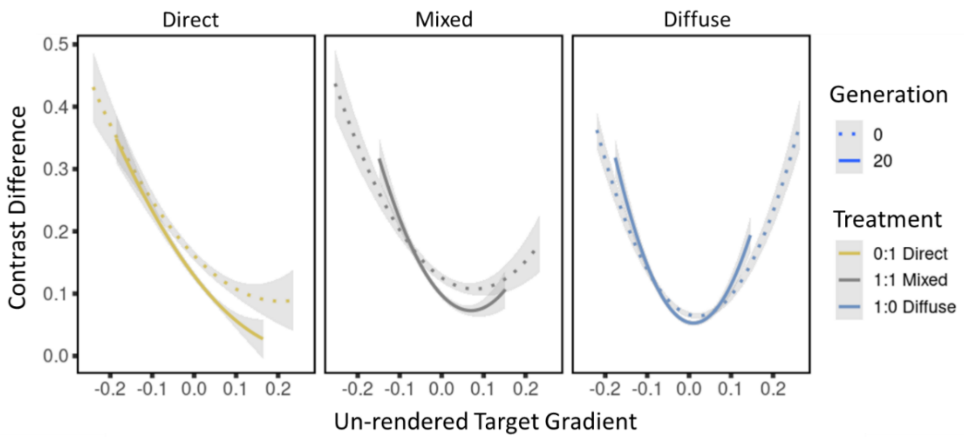


Figure SA14, Comparison of relationship between the luminance (L*) of the un-rendered target-skin and the luminance gradient of the target rendered against the average grey sphere for its respective habitat under its original lighting condition.

Table SA2, Stats output for the effect of the target-skins luminance gradient on the absolute difference between the target contrast and the background. Linear mixed model formula: lmer(ContrastDifference ~ Generation * LuminanceGradient * Light + (1|Habitat) + (1|Population) …

| | Predictor | β | SE | DF | t | p value |
|---|---|---|---|---|---|---|
| *Contrast Difference* | Intercept | 0.447 | 0.06 | 27 | 7.42 | <0.001 |
| | Generation | -0.120 | 0.00 | 36210 | -25.12 | <0.001 |
| | Lum Gradient | -77.270 | 0.91 | 36260 | -84.55 | <0.001 |
| | Mixed | 19.670 | 0.95 | 36250 | 20.72 | <0.001 |
| | Diffuse | -0.869 | 0.01 | 23700 | -73.74 | <0.001 |
| | Gen : LGrad | -4.880 | 0.83 | 36230 | -5.85 | <0.001 |
| | Gen : Mixed | -0.553 | 0.79 | 36230 | -0.70 | 0.486 |
| | Gen : Diffuse | 0.069 | 0.01 | 36210 | 9.95 | <0.001 |
| | LGrad: Mixed | 65.220 | 1.42 | 36240 | 45.87 | <0.001 |
| | LGrad : Diffuse | 66.160 | 1.53 | 36250 | 43.12 | <0.001 |
| | Gen : LGrad : Mixed | 5.395 | 1.30 | 36220 | 4.15 | <0.001 |
| | Gen : LGrad : Diffuse | 9.214 | 1.26 | 36230 | 7.31 | <0.001 |

#### Glossiness and Luminance

The darker the luminance of the target-skin the greater the effect of glossiness on the target's luminance (Figure SA15, Table SA3). To test whether this finding was significant a linear mixed model was used with glossiness (increase in mean L*) as the response variable and the pattern-skin L* mean, n-generations and lighting treatment as fixed effects. As expected, brighter targets had lower glossiness, except in Generation 20 Diffuse evolved targets had higher glossiness, likely due to glossiness being used to bolster brightness.

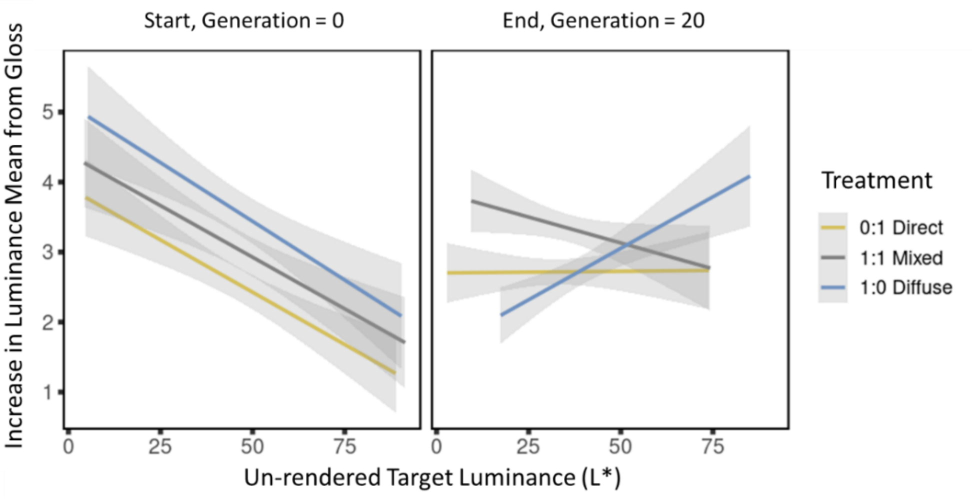


Figure SA15, the effect of target un-rendered luminance on the change in luminance mean from glossiness. Left shows the starting generation (0) and right shows the final generation (20).

---

Table SA3, Stats output for the effect of the target-skins luminance (L*) on the shift in luminance mean from gloss. Linear mixed model formula: lmer(Glossiness ~ Generation * TargetLuminance * Light + (1|Habitat) + (1|Population)

| | Predictor | β | SE | DF | t | p value |
|---|---|---|---|---|---|---|
| *Glossiness* | Intercept | -0.081 | 0.10 | 69 | -0.85 | 0.400 |
| | Generation | 0.026 | 0.01 | 36230 | 3.19 | 0.001 |
| | Skin Luminance | -0.102 | 0.01 | 36130 | -9.65 | <0.001 |
| | Mixed | 0.219 | 0.14 | 69 | 1.59 | 0.116 |
| | Diffuse | 0.051 | 0.14 | 69 | 0.37 | 0.713 |
| | Gen : Lum | 0.021 | 0.01 | 36280 | 2.61 | 0.009 |
| | Gen : Mixed | 0.038 | 0.01 | 36220 | 3.24 | 0.001 |
| | Gen : Diffuse | -0.091 | 0.01 | 36220 | -7.74 | <0.001 |
| | Lum: Mixed | 0.021 | 0.02 | 36200 | 1.41 | 0.159 |
| | Lum : Diffuse | 0.006 | 0.02 | 36060 | 0.36 | 0.716 |
| | Gen : Lum : Mixed | 0.038 | 0.01 | 36270 | 3.32 | 0.001 |
| | Gen : Lum : Diffuse | 0.036 | 0.01 | 36280 | 3.08 | 0.002 |

#### Examples of Masquerade

Numerous instances of apparent masquerade of non-local objects were observed for different habitats, in particular backgrounds which included stones or individual leaves of equal size to the target (Figure SA16).

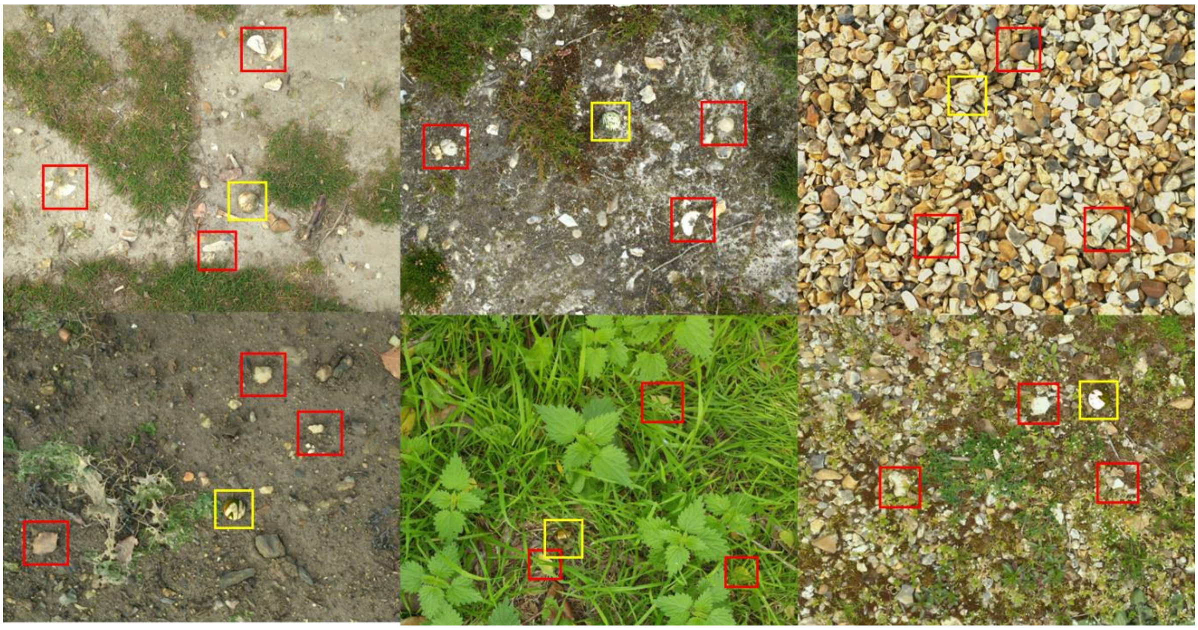

Figure SA16, examples of target masquerade against diffuse backgrounds. The targets are highlighted with a yellow square and the candidate model objects are highlighted in red, three for each background.

#### *References*

Cuthill, I.C., Allen, W.L., Arbuckle, K., Caspers, B., Chaplin, G., Hauber, M.E., Hill, G.E., Jablonski, N.G., Jiggins, C.D., Kelber, A., 2017. The biology of color. Science 357, eaan0221.

Donohue, C.G., Hemmi, J.M., Kelley, J.L., 2020. Countershading enhances camouflage by reducing prey contrast. Proc. R. Soc. B 287, 20200477.

Hancock, G.R.A., Troscianko, J., 2022. CamoEvo: An open access toolbox for artificial camouflage evolution experiments. Evolution 76, 870–882. https://doi.org/10.1111/evo.14476

Ruxton, G.D., Speed, M.P., Kelly, D.J., 2004. What, if anything, is the adaptive function of countershading? Anim. Behav. 68, 445–451.

Shawkey, M.D., D’Alba, L., 2017. Interactions between colour-producing mechanisms and their effects on the integumentary colour palette. Philos. Trans. R. Soc. B Biol. Sci. 372, 20160536.
