## Supplemental Text B - Stats Tables for "Shining a light on camouflage evolution: using genetic algorithms to determine the effects of geometry and lighting on optimal camouflage"

#### Background Variation and Fitness

Table SB1. Stats output for the effect of global background variation in the luminance L*), green-red (a*), blue-yellow (b*) and depth (Z) on fitness and the evolution of increased fitness. The lower the effect size the redder the ***β*** value.

| | Effect | | | | | Effect : Generation | | | | |
|---|---|---|---|---|---|---|---|---|---|---|
| Fixed Effect | β | SE | SD | t | p value | β | SE | SD | t | p value |
| L | 0.027 | 0.00182 | 42330 | 15.000 | <0.001 | 0.001 | 0.000155 | 42330 | 5.767 | <0.001 |
| A | -0.011 | 0.001852 | 42330 | -5.737 | <0.001 | 0.002 | 0.000157 | 42330 | 13.142 | <0.001 |
| B | 0.002 | 0.001851 | 42330 | 1.088 | 0.2764 | 0.002 | 0.000156 | 42330 | 14.241 | <0.001 |
| Z | -0.003 | 0.001842 | 42330 | -1.760 | 0.0784 | 0.002 | 0.000157 | 42330 | 11.400 | <0.001 |
| LA | 0.034 | 0.000978 | 42190 | 35.100 | <0.001 | 0.013 | 0.00094 | 42330 | 13.500 | <0.001 |
| LB | 0.018 | 0.001831 | 42330 | 9.823 | <0.001 | 0.002 | 0.000155 | 42330 | 11.786 | <0.001 |
| LZ | 0.019 | 0.001821 | 42330 | 10.300 | <0.001 | 0.002 | 0.000155 | 42330 | 12.760 | <0.001 |
| AB | -0.005 | 0.001854 | 42330 | -2.494 | 0.0126 | 0.002 | 0.000157 | 42330 | 15.081 | <0.001 |
| AZ | -0.008 | 0.001847 | 42330 | -4.268 | <0.001 | 0.002 | 0.000157 | 42330 | 15.081 | <0.001 |
| BZ | -0.001 | 0.001846 | 42330 | -0.296 | 0.7673 | 0.002 | 0.000156 | 42330 | 15.322 | <0.001 |
| LAB | 0.010 | 0.001841 | 42320 | 5.175 | <0.001 | 0.002 | 0.000155 | 42330 | 14.613 | <0.001 |
| LAZ | 0.009 | 0.001835 | 42330 | 4.646 | <0.001 | 0.002 | 0.000155 | 42330 | 15.740 | <0.001 |
| LBZ | 0.014 | 0.001832 | 42320 | 7.396 | <0.001 | 0.002 | 0.000155 | 42330 | 14.814 | <0.001 |
| ABZ | -0.005 | 0.00185 | 42330 | -2.442 | 0.0146 | 0.002 | 0.000157 | 42330 | 15.809 | <0.001 |
| LABZ | 0.007 | 0.00184 | 42320 | 3.797 | <0.001 | 0.002 | 0.000155 | 42330 | 16.067 | <0.001 |

#### Evolution and Fitness (Survival Time)

##### *Lighting Treatments*

Table SB2. Stats output for the effect of the number of generations, lighting and background 3D variation on fitness, excluding outliers. Linear mixed model formula:

lmer(log(SurvivalTime) ~ Generation * Light * $3D_{Variation}$ + (1|Habitat) + (1|Population) …

| | Predictor | β | SE | DF | t | p value |
|---|---|---|---|---|---|---|
| *log Fitness (Capture Time)* | Intercept | 0.471 | 0.02 | 35 | 29.43 | <0.001 |
| | Generation | 0.050 | 0.00 | 36210 | 31.08 | <0.001 |
| | Mixed | -0.024 | 0.01 | 41 | -2.35 | 0.024 |
| | Diffuse | -0.020 | 0.01 | 42 | -1.84 | 0.072 |
| | Mean 3D Var | 0.033 | 0.02 | 33 | 1.97 | 0.057 |
| | Gen : Mixed | -0.009 | 0.00 | 36210 | -4.17 | <0.001 |
| | Gen : Diffuse | -0.004 | 0.00 | 36210 | -1.90 | 0.058 |
| | Gen : 3D | 0.009 | 0.00 | 36210 | 5.90 | 0.000 |
| | Mixed : 3D | -0.022 | 0.01 | 41 | -2.10 | 0.042 |
| | Diffuse : 3D | -0.025 | 0.01 | 41 | -2.43 | 0.020 |
| | Gen : Mixed : 3D | -0.002 | 0.00 | 36210 | -0.72 | 0.470 |
| | Gen : Diffuse : 3D | 0.005 | 0.00 | 36210 | 2.26 | 0.024 |

##### *Mixed Lighting*

Table SB3. Stats output for the effect of the number of generations, background lighting and background 3D variation on fitness for mixed lighting populations, excluding outliers. Linear mixed model formula:

lmer(log(SurvivalTime) ~ Generation * Light * $3D_{Variation}$ + (1|Habitat) + (1|Population) …

| | Predictor | β | SE | DF | t | p value |
|---|---|---|---|---|---|---|
| *log Fitness (Capture Time)* | Intercept | 0.456 | 0.02 | 22 | 27.22 | <0.001 |
| | Generation | 0.040 | 0.00 | 12070 | 17.69 | <0.001 |
| | bg Diffuse | -0.014 | 0.00 | 12070 | -4.33 | <0.001 |
| | 3D | 0.014 | 0.02 | 22 | 0.85 | 0.404 |
| | Gen : Diffuse | 0.001 | 0.00 | 12070 | 0.25 | 0.806 |
| | Gen : 3D | 0.011 | 0.00 | 12070 | 4.91 | 0.000 |
| | Diffuse: 3D | -0.011 | 0.00 | 12070 | -3.39 | 0.001 |
| | Gen : Diffuse : 3D | -0.007 | 0.00 | 12070 | -2.11 | 0.035 |

#### Camouflage Metrics and Fitness (Survival Time)

##### *Luminance Difference*

Table SB4. Stats output for the effect of luminance match, lighting and background 3D variation on fitness, excluding outliers. Linear mixed model formula:

lmer(log(SurvivalTime) ~ $Luminance_{Difference}$ * Light * $3D_{Variation}$ + (1|Habitat) + (1|Population) …

| | Predictor | β | SE | DF | t | p value |
|---|---|---|---|---|---|---|
| *log Fitness (Capture Time)* | Intercept | 0.470 | 0.02 | 34 | 29.61 | <0.001 |
| | L Difference | -0.036 | 0.00 | 35780 | -20.21 | <0.001 |
| | Mixed | -0.022 | 0.01 | 41 | -2.19 | 0.034 |
| | Diffuse | -0.018 | 0.01 | 42 | -1.70 | 0.096 |
| | Mean 3D Var | 0.028 | 0.02 | 33 | 1.72 | 0.095 |
| | LD : Mixed | 0.003 | 0.00 | 35550 | 1.07 | 0.285 |
| | LD : Diffuse | 0.007 | 0.00 | 35910 | 2.75 | 0.006 |
| | LD : 3D | -0.011 | 0.00 | 36250 | -5.89 | 0.000 |
| | Mixed : 3D | -0.019 | 0.01 | 41 | -1.89 | 0.065 |
| | Diffuse : 3D | -0.020 | 0.01 | 41 | -1.94 | 0.060 |
| | LD : Mixed : 3D | 0.001 | 0.00 | 36250 | 0.41 | 0.683 |
| | LD : Diffuse : 3D | 0.003 | 0.00 | 36230 | 1.04 | 0.301 |

#### *Colour Difference*

Table SB5. Stats output for the effect of colour difference (Euclidean distance of a* and b*), lighting and background 3D variation on fitness, excluding outliers. Linear mixed model formula:

lmer(log(SurvivalTime) ~ $\text{Colour}_{\text{Difference}}$ * Light * $\text{3D}_{\text{Variation}}$ + (1|Habitat) + (1|Population) …

| | Predictor | β | SE | DF | t | p value |
|---|---|---|---|---|---|---|
| *log Fitness (Capture Time)* | Intercept | 0.468 | 0.01 | 36 | 31.49 | <0.001 |
| | AB Difference | -0.043 | 0.00 | 35500 | -23.53 | <0.001 |
| | Mixed | -0.020 | 0.01 | 41 | -1.99 | 0.053 |
| | Diffuse | -0.014 | 0.01 | 42 | -1.26 | 0.215 |
| | Mean 3D Var | 0.031 | 0.02 | 34 | 2.00 | 0.054 |
| | ABD : Mixed | 0.004 | 0.00 | 35470 | 1.47 | 0.142 |
| | ABD : Diffuse | 0.005 | 0.00 | 35780 | 2.19 | 0.028 |
| | ABD : 3D | -0.008 | 0.00 | 35940 | -4.96 | <0.001 |
| | Mixed : 3D | -0.018 | 0.01 | 41 | -1.76 | 0.087 |
| | Diffuse : 3D | -0.021 | 0.01 | 41 | -2.05 | 0.046 |
| | ABD : Mixed : 3D | 0.008 | 0.00 | 35910 | 3.12 | 0.002 |
| | ABD : Diffuse : 3D | 0.004 | 0.00 | 36140 | 1.91 | 0.056 |

#### *Pattern Contrast Difference*

Table SB6. Stats output for the effect of pattern difference (difference in L* StdDev), lighting and background 3D variation on fitness, excluding outliers. Linear mixed model formula:

lmer(log(SurvivalTime) ~ $\text{Pattern}_{\text{Difference}}$ * Light * $\text{3D}_{\text{Variation}}$ + (1|Habitat) + (1|Population) …

| | Predictor | β | SE | DF | t | p value |
|---|---|---|---|---|---|---|
| *log Fitness (Capture Time)* | Intercept | 0.478 | 0.02 | 35 | 29.89 | <0.001 |
| | Pattern Difference | -0.036 | 0.00 | 34420 | -18.66 | <0.001 |
| | Mixed | -0.031 | 0.01 | 42 | -2.95 | 0.005 |
| | Diffuse | -0.043 | 0.01 | 44 | -3.85 | <0.001 |
| | Mean 3D Var | 0.020 | 0.02 | 33 | 1.23 | 0.227 |
| | PD : Mixed | 0.023 | 0.00 | 34240 | 8.58 | <0.001 |
| | PD : Diffuse | 0.007 | 0.00 | 34930 | 2.21 | 0.027 |
| | PD : 3D | -0.020 | 0.00 | 34980 | -9.13 | <0.001 |
| | Mixed : 3D | -0.010 | 0.01 | 41 | -0.92 | 0.361 |
| | Diffuse : 3D | -0.026 | 0.01 | 44 | -2.46 | 0.018 |
| | PD : Mixed : 3D | 0.011 | 0.00 | 35480 | 3.70 | <0.001 |
| | PD : Diffuse : 3D | 0.001 | 0.00 | 35390 | 0.41 | 0.682 |

#### *Edge Disruption (L* GabRat)*

Table SB7. Stats output for the effect edge disruption, lighting and background 3D variation on fitness, excluding outliers. Linear mixed model formula:

lmer(log(SurvivalTime) ~ GabRat * Light * $3D_{Variation}$ + (1|Habitat) + (1|Population) …

| | Predictor | β | SE | DF | t | p value |
|---|---|---|---|---|---|---|
| *log Fitness (Capture Time)* | Intercept | 0.469 | 0.02 | 33 | 28.08 | <0.001 |
| | L GabRat | 0.040 | 0.00 | 33860 | 19.93 | <0.001 |
| | Mixed | -0.021 | 0.01 | 41 | -2.12 | 0.041 |
| | Diffuse | -0.016 | 0.01 | 42 | -1.57 | 0.125 |
| | Mean 3D Var | 0.029 | 0.02 | 32 | 1.67 | 0.104 |
| | Gab : Mixed | -0.009 | 0.00 | 31400 | -3.32 | 0.001 |
| | Gab : Diffuse | -0.014 | 0.00 | 34020 | -5.17 | <0.001 |
| | Gab : 3D | 0.011 | 0.00 | 36190 | 5.09 | <0.001 |
| | Mixed : 3D | -0.021 | 0.01 | 41 | -2.12 | 0.040 |
| | Diffuse : 3D | -0.025 | 0.01 | 41 | -2.46 | 0.018 |
| | Gab : Mixed : 3D | -0.002 | 0.00 | 35750 | -0.83 | 0.406 |
| | Gab : Diffuse : 3D | -0.003 | 0.00 | 36110 | -1.16 | 0.245 |

### A4.2.4 Phenotype Metrics

#### *PC Contributions*

Table SB8. Contributions to principal components (PC), PC1 and PC2 shown in ranked absolute order for variables that contribute greater than equal to 0.10. Positive values are shown in blue, negative values in red. Negative verticalness indicates increasing horizontalness.

| Variable | PC1 Contribution |
|---|---|
| Contrast 1/2 b* | 0.206 |
| Verticalness 1/4 b* | -0.205 |
| Verticalness 1/2 b* | -0.200 |
| Contrast 1/4 b* | 0.197 |
| Verticalness 1/8 b* | -0.195 |
| Contrast 1/8 b* | 0.193 |
| Contrast 1/2 L* | 0.193 |
| Contrast 1/1 b* | 0.193 |
| Verticalness 1/4 L* | -0.191 |
| Verticalness 1/1 b* | -0.191 |
| Verticalness 1/2 L* | -0.191 |
| Contrast 1/4 L* | 0.185 |
| Contrast 1/2 a* | 0.181 |
| Verticalness 1/4 a* | -0.181 |
| Contrast 1/1 L* | 0.180 |
| Verticalness 1/8 a* | -0.179 |
| Verticalness 1/2 a* | -0.179 |
| Verticalness 1/1 L* | -0.177 |
| Contrast 1/8 a* | 0.176 |
| Verticalness 1/8 L* | -0.175 |
| Contrast 1/1 a* | 0.175 |
| Verticalness 1/1 a* | -0.174 |
| Contrast 1/4 a* | 0.172 |
| Contrast 1/8 L* | 0.162 |
| Contrast 1/16 b* | 0.152 |
| Contrast 1/16 a* | 0.143 |
| Contrast 1/16 L* | 0.126 |
| Contrast 1/32 b* | 0.117 |
| Contrast 1/32 a* | 0.113 |
| Contrast 1/32 L* | 0.109 |

| Variable | PC2 Contribution |
|---|---|
| Contrast 1/16 L* | 0.203 |
| Contrast 1/16 b* | 0.199 |
| Verticalness 1/8 a* | 0.194 |
| Verticalness 1/16 b* | 0.192 |
| Contrast 1/8 L* | 0.190 |
| Verticalness 1/8 b* | 0.185 |
| Contrast 1/8 b* | 0.179 |
| Verticalness 1/16 L* | 0.176 |
| Verticalness 1/16 a* | 0.174 |
| Verticalness 1/4 a* | 0.172 |
| Contrast 1/32 L* | 0.172 |
| Verticalness 1/2 a* | 0.169 |
| Contrast 1/32 b* | 0.168 |
| Verticalness 1/8 L* | 0.167 |
| Contrast 1/16 a* | 0.164 |
| Verticalness 1/4 b* | 0.161 |
| Contrast 1/4 L* | 0.160 |
| Verticalness 1/2 b* | 0.159 |
| Verticalness 1/1 a* | 0.157 |
| Verticalness 1/4 L* | 0.157 |
| Verticalness 1/32 b* | 0.155 |
| Verticalness 1/1 b* | 0.153 |
| Verticalness 1/2 L* | 0.153 |
| Verticalness 1/32 L* | 0.146 |
| Contrast 1/4 b* | 0.144 |
| Contrast 1/8 a* | 0.142 |
| Contrast 1/2 L* | 0.142 |
| Directionality 1/32 L* | 0.138 |
| Directionality 1/32 b* | 0.137 |
| Contrast 1/32 a* | 0.136 |
| Verticalness 1/1 L* | 0.136 |
| Verticalness 1/32 a* | 0.129 |
| Contrast 1/2 b* | 0.127 |
| Directionality 1/32 a* | 0.120 |
| Contrast 1/1 L* | 0.117 |
| Contrast 1/4 a* | 0.111 |
| Contrast 1/1 b* | 0.109 |
| Contrast 1/2 a* | 0.100 |

Table SB9. Contributions to principal components (PC), PC3 and PC4 shown in ranked absolute order for variables that contribute greater than equal to 0.10. Positive values are shown in blue, negative values in red. Negative verticalness indicates increasing horizontalness.

| Variable | PC3 Contribution |
|---|---|
| Contrast 1/2 a* | -0.361 |
| Contrast 1/4 a* | -0.356 |
| Contrast 1/8 a* | -0.340 |
| Contrast 1/1 a* | -0.313 |
| Contrast 1/16 a* | -0.297 |
| Contrast 1/32 a* | -0.297 |
| Contrast 1/16 b* | 0.137 |
| Verticalness 1/2 a* | 0.130 |
| Contrast 1/32 b* | 0.130 |
| Contrast 1/1 L* | 0.127 |
| Contrast 1/8 b* | 0.126 |
| Contrast 1/2 L* | 0.126 |
| Verticalness 1/4 a* | 0.125 |
| Verticalness 1/1 a* | 0.125 |
| Contrast 1/2 b* | 0.122 |
| Contrast 1/1 b* | 0.120 |
| Contrast 1/8 L* | 0.116 |
| Contrast 1/16 L* | 0.116 |
| Verticalness 1/1 L* | -0.111 |
| Contrast 1/32 L* | 0.110 |
| Contrast 1/4 b* | 0.108 |
| Verticalness 1/2 L* | -0.108 |
| Contrast 1/4 L* | 0.106 |
| Verticalness 1/8 a* | 0.104 |

| Variable | PC4 Contribution |
|---|---|
| Directionality 1/32 a* | 0.297 |
| Directionality 1/32 b* | 0.285 |
| Directionality 1/32 L* | 0.277 |
| Verticalness 1/32 L* | 0.275 |
| Verticalness 1/32 b* | 0.265 |
| Verticalness 1/32 a* | 0.257 |
| Directionality 1/16 a* | 0.235 |
| Directionality 1/16 b* | 0.222 |
| Directionality 1/16 L* | 0.214 |
| Verticalness 1/16 L* | 0.196 |
| Verticalness 1/16 b* | 0.195 |
| Verticalness 1/2 a* | -0.155 |
| Verticalness 1/16 a* | 0.152 |
| Verticalness 1/4 a* | -0.149 |
| Contrast 1/4 b* | -0.142 |
| Verticalness 1/1 a* | -0.139 |
| Mean L* | -0.135 |
| Contrast 1/2 b* | -0.133 |
| Directionality 1/8 a* | 0.132 |
| Directionality 1/8 b* | 0.112 |
| Contrast 1/8 b* | -0.111 |
| Contrast 1/1 b* | -0.111 |
| Contrast 1/4 L* | -0.101 |

#### *PC1 Model*

Table SB10. Stats output for the effect number of generations, lighting and background 3D variation on the targets PC1 value, excluding outliers. Linear mixed model formula:

lmer(PC1 ~ Generation * Light * $3D_{Variation}$ + (1|Habitat) + (1|Population) …

| | Predictor | β | SE | DF | t | p value |
|---|---|---|---|---|---|---|
| *PC1* | Intercept | 0.108 | 0.06 | 66 | 1.90 | 0.062 |
| | Generation | -0.042 | 0.01 | 36210 | -4.97 | <0.001 |
| | Mixed | -0.059 | 0.08 | 66 | -0.73 | 0.465 |
| | Diffuse | -0.276 | 0.08 | 66 | -3.37 | 0.001 |
| | Mean 3D Var | -0.054 | 0.06 | 66 | -0.94 | 0.351 |
| | Gen : Mixed | -0.096 | 0.01 | 36210 | -8.04 | <0.001 |

| | | | | | | |
|---|---|---|---|---|---|---|
| | Gen : Diffuse | -0.201 | 0.01 | 36210 | -16.63 | <0.001 |
| | Gen : 3D | 0.026 | 0.01 | 36210 | 3.04 | 0.002 |
| | Mixed : 3D | 0.123 | 0.08 | 66 | 1.51 | 0.135 |
| | Diffuse : 3D | -0.009 | 0.08 | 66 | -0.12 | 0.908 |
| | Gen : Mixed : 3D | 0.015 | 0.01 | 36210 | 1.24 | 0.214 |
| | Gen : Diffuse : 3D | -0.072 | 0.01 | 36210 | -5.99 | <0.001 |

#### A4.2.5 Countershading

Table SB11. Stats output for the effect number of generations, lighting and background 3D variation on the targets shift in luminance gradient from the grey target to the rendered target, excluding outliers. Linear mixed model formula: lmer( CS_Gradient ~ Generation * Light * $3D_{Variation}$ + (1|Habitat) + (1|Population) …

| | Predictor | β | SE | DF | t | p value |
|---|---|---|---|---|---|---|
| *Gradient Shift* | Intercept | -0.124 | 0.12 | 31 | -1.00 | 0.323 |
| | Generation | 0.022 | 0.00 | 36210 | 20.99 | <0.001 |
| | Mixed | -0.034 | 0.06 | 46 | -0.57 | 0.573 |
| | Diffuse | -0.053 | 0.06 | 46 | -0.84 | 0.405 |
| | Mean 3D Var | 0.270 | 0.13 | 30 | 2.09 | 0.045 |
| | Gen : Mixed | -0.008 | 0.00 | 36210 | -5.36 | <0.001 |
| | Gen : Diffuse | -0.018 | 0.00 | 36210 | -12.00 | <0.001 |
| | Gen : 3D | 0.011 | 0.00 | 36210 | 10.11 | <0.001 |
| | Mixed : 3D | -0.135 | 0.06 | 46 | -2.25 | 0.029 |
| | Diffuse : 3D | -0.037 | 0.06 | 46 | -0.60 | 0.550 |
| | Gen : Mixed : 3D | 0.000 | 0.00 | 36210 | 0.28 | 0.777 |
| | Gen : Diffuse : 3D | -0.008 | 0.00 | 36210 | -5.18 | <0.001 |

#### A4.2.6 Pattern Direction

##### *Directionality*

Table SB12. Stats output for the effect number of generations, lighting and background 3D variation on the directionality of the targets pattern at 1/32x, excluding outliers. Linear mixed model formula: lmer(CS_Gradient ~ Generation * Light * $3D_{Variation}$ + (1|Habitat) + (1|Population) …

| | Predictor | β | SE | DF | t | p value |
|---|---|---|---|---|---|---|
| *Directionality 1/32* | Intercept | 0.114 | 0.06 | 66 | 1.84 | 0.070 |
| | Generation | 0.023 | 0.01 | 36210 | 2.81 | 0.005 |
| | Mixed | -0.054 | 0.09 | 66 | -0.61 | 0.543 |
| | Diffuse | -0.299 | 0.09 | 66 | -3.34 | 0.001 |
| | Mean 3D Var | 0.166 | 0.06 | 66 | 2.67 | 0.010 |
| | Gen : Mixed | -0.004 | 0.01 | 36210 | -0.30 | 0.762 |
| | Gen : Diffuse | -0.126 | 0.01 | 36210 | -10.46 | <0.001 |
| | Gen : 3D | 0.065 | 0.01 | 36210 | 7.80 | <0.001 |
| | Mixed : 3D | -0.233 | 0.09 | 66 | -2.63 | 0.011 |
| | Diffuse : 3D | -0.175 | 0.09 | 66 | -1.96 | 0.055 |
| | Gen : Mixed : 3D | -0.095 | 0.01 | 36210 | -7.93 | <0.001 |
| | Gen : Diffuse : 3D | -0.096 | 0.01 | 36210 | -7.95 | <0.001 |

#### *Verticality*

Table SB13. Stats output for the effect number of generations, lighting and background 3D variation on the verticality of the targets pattern at 1/32x, excluding outliers. Linear mixed model formula: lmer(CS_Gradient ~ Generation * Light * $3D_{Variation}$ + (1|Habitat) + (1|Population) …

| | Predictor | β | SE | DF | t | p value |
|---|---|---|---|---|---|---|
| *Directionality 1/32* | Intercept | 0.073 | 0.05 | 66 | 1.42 | 0.162 |
| | Generation | 0.086 | 0.01 | 36210 | 10.09 | <0.001 |
| | Mixed | -0.087 | 0.07 | 45 | -1.20 | 0.237 |
| | Diffuse | -0.138 | 0.07 | 49 | -1.89 | 0.065 |
| | Mean 3D Var | 0.158 | 0.05 | 66 | 3.02 | 0.004 |
| | Gen : Mixed | -0.031 | 0.01 | 36210 | -2.58 | 0.010 |
| | Gen : Diffuse | -0.073 | 0.01 | 36210 | -5.94 | <0.001 |
| | Gen : 3D | 0.095 | 0.01 | 36210 | 11.10 | <0.001 |
| | Mixed : 3D | -0.280 | 0.07 | 44 | -3.84 | <0.001 |
| | Diffuse : 3D | -0.182 | 0.07 | 45 | -2.49 | 0.017 |
| | Gen : Mixed : 3D | -0.176 | 0.01 | 36210 | -14.35 | <0.001 |
| | Gen : Diffuse : 3D | -0.126 | 0.01 | 36210 | -10.26 | <0.001 |

#### *Edge Enhancement*

Table SB14. Stats output for the effect number of generations, lighting and background 3D variation on the edge enhancement of the targets (change in luminance contrast at 1/64x scale), excluding outliers. Linear mixed model formula: lmer(CS_Gradient ~ Generation * Light * $3D_{Variation}$ + (1|Habitat) + (1|Population) …

| | Predictor | β | SE | DF | t | p value |
|---|---|---|---|---|---|---|
| *Edge Enhancement* | Intercept | 0.040 | 0.08 | 60 | 0.49 | 0.628 |
| | Generation | 0.043 | 0.01 | 36210 | 5.31 | <0.001 |
| | Mixed | -0.014 | 0.10 | 40 | -0.14 | 0.887 |
| | Diffuse | -0.091 | 0.10 | 43 | -0.88 | 0.383 |
| | Mean 3D Var | 0.109 | 0.08 | 58 | 1.32 | 0.193 |
| | Gen : Mixed | 0.024 | 0.01 | 36210 | 2.09 | 0.037 |
| | Gen : Diffuse | -0.030 | 0.01 | 36210 | -2.54 | 0.011 |
| | Gen : 3D | 0.027 | 0.01 | 36210 | 3.28 | 0.001 |
| | Mixed : 3D | 0.050 | 0.10 | 39 | 0.50 | 0.622 |
| | Diffuse : 3D | -0.120 | 0.10 | 40 | -1.18 | 0.247 |
| | Gen : Mixed : 3D | 0.077 | 0.01 | 36210 | 6.59 | <0.001 |
| | Gen : Diffuse : 3D | -0.014 | 0.01 | 36210 | -1.21 | 0.228 |
