## Supplementary material for "Shining a light on camouflage evolution: using genetic algorithms to determine the effects of geometry and lighting on optimal camouflage": RCode_CamoEvo_Online.R

```
'red' #NB my work computer has a bug where I have to set a manual R directory
# Please delete before using
.libPaths()
new_library_path <- "C:\\Users\\Localadmin_hangeorg\\Documents\\RStudio\\library\\"
.libPaths(new_library_path)


set.seed(123)


#________________________________________________________________________________
#________________________________________________________________________________

#----Lighting and Camouflage Evolution (CamoEvo Online) R Code ---- 
#________________________________________________________________________________
#________________________________________________________________________________

#Here is the finalised analyeses for the CamoEvo Online Project.


#Step 1 Load Package GAMUT


#______________________________________

#---- Packages ---- 
#______________________________________

library(tidyverse)
library(gtools)
library(munsell)
library(colorspace)
library(lme4)
library(lmerTest)
library(plotly)
library(emmeans)
library(ggplot2)
library(ggimage)
library(scales)
library(Matrix)
library(lme4)
library(nlme)
library(lmerTest)
library(pbkrtest)
library(car)
library(lattice)
library(gridExtra)
library(lme4)
library(coxme)
library(gtools)
library(robustlmm)
library(EnvStats)


#______________________________________

#---- Functions ---- 
#______________________________________


mxr <- function(x) {
  x/max(abs(x))
}

mmxn <- function(x) {
  (x - min(x)) / (max(x) - min(x))
}

mmxnG <- function(x) {
  ((x - min(x)) / (max(x) - min(x))) + 1
}

zscore<- function(x) {
  (x - mean(x)) / sd(x)
}


#______________________________________

#---- Themes ---- 
#______________________________________

Hex_Direct2 = "#D5C15F"
Hex_Diffuse2 = "#7294C2"
Hex_Hybrid2 = "#848484"

Hex_Direct = "#E69F00"
Hex_Diffuse = "#56B4E9"
Hex_Hybrid = "#999999"


Hex_Direct_Light = "#ffbe33"
Hex_Diffuse_Light = "#4aaee8"
Hex_Hybrid_Light = "#cccccc"

Hex_Direct_Dark = "#996900"
Hex_Diffuse_Dark = "#125c87"
Hex_Hybrid_Dark = "#4d4d4d"
Hex_Red = "#EE7C7C"


Lighting_Palette <- c(Hex_Direct2, Hex_Hybrid2 , Hex_Diffuse2,  Hex_Red)
Lighting_Levels <- c("0:1 Direct","1:1 Mixed","1:0 Diffuse")
Lighting_Palette2 <- c(Hex_Diffuse2, Hex_Hybrid2 , Hex_Direct2,  Hex_Red)


bgLighting_Palette <- c(Hex_Direct2 , Hex_Diffuse2)

LightingAge_Palette <- c(Hex_Direct_Light , Hex_Hybrid_Light, Hex_Diffuse_Light,Hex_Direct_Dark , Hex_Hybrid_Dark, Hex_Diffuse_Dark)


#______________________________________
#~~~~~~~~~~~~~~~~~~~~~~~~~~~~~~~~~~~~~~
#----.......................----
#----(DATA) Datasets ---- 
#~~~~~~~~~~~~~~~~~~~~~~~~~~~~~~~~~~~~~~
#______________________________________


#Set Directory to the R Code Directory
setwd(dirname(rstudioapi::getSourceEditorContext()$path)) 

# Open the data file
# Reorganize Variables


#.............................

#----....[DATA] Import ----
#.............................

#BASE
df.base <-read.table("Datasets/Base_Data.txt", header=TRUE)

#BG
df.bg.1 <-read.table("Datasets/LightCamo_BG_2.5.txt", header=TRUE)
df.bg.4 <-read.table("Datasets/LightCamo_BG_15.txt", header=TRUE)


#BG-info
df.bg.info <-read.table("Datasets/BG_Info.txt", header=TRUE)
df.bg.site <-read.table("Datasets/BG_SiteInfo.txt", header=TRUE)

#TG
df.tg.render <-read.table("Datasets/LightCamo_Render.txt", header=TRUE)
df.tg.skin <-read.table("Datasets/LightCamo_Skin.txt", header=TRUE)


#Scan
df.scan.all <-read.table("Datasets/BG_Geometry.txt", header=TRUE)

df.tg.eeF <-read.table("Datasets/LightCamo_EdgeEnhance.txt", header=TRUE)

#.............................

#----....[DATA] Merge ----
#.............................

#Note the second entry is the one it gets merged to


df.base$ShowOrder = 0

c=0
for (idx in 1:length(df.base$ShowOrder))
{
  df.base$ShowOrder[[idx]] = c
  c=c+1
  if(c>23){c=0}
}
c


DF = df.base

DF=merge(DF,df.bg.info)
DF=merge(DF,df.bg.site)

##BG
BG = df.bg.1
BG = merge(BG,df.bg.4)       
BG = merge(df.scan.all, BG)

DF=merge(BG,DF)


##TG
TG = df.tg.render
TG = merge(df.tg.skin,TG)

DF=merge(TG,DF)


nrow(DF)/21/28/3 # There should be 24 entries


p1 <- ggplot(subset(DF, Habitat =="07_woodLichen" & Lighting =="1_Natural"), aes(as.factor(Generation), log(Fitness),color=Lighting)) +
  geom_point(alpha=0.5,size=3)+ facet_grid(. ~Lighting)
p1 


#.............................

#----....[DATA] Add new Labels ----
#.............................

DF$Lighting=DF$Lighting.1
unique(DF$Lighting)

DF$S.M.F = factor(DF$S.M.F, levels = c("Start", "Middle", "End"))


DF$Population <- paste(DF$Habitat, DF$Lighting,sep="_") 
DF$Play  <- paste(DF$Population, DF$Generation,sep="_") 
DF$UniquePhoto  <- paste(DF$Habitat, DF$Background.ID,sep="_")  

DF$UniqueTarget  <- paste(DF$Population, DF$ID,sep="_")  


DF$FactorFit = quantcut(DF$Fitness,3)

DF$Thirds = as.factor(DF$Thirds)

DF$DeathWitnessed <- 1

for (idx in 1:length(DF$DeathWitnessed))
{
  if(DF$Fitness[[idx]]==15000) DF$DeathWitnessed[[idx]] = 0
}


# Categorical Generation

DF$Cat_Gen<-as.factor(DF$Generation)

DF$UniquePop = paste(DF$Population, DF$Generation,sep="_")  


# Background Lighting

DF$Background_Lighting<-as.vector(DF$Lighting)

for (i in 1:length(DF$Background_Lighting)){
  tmp=as.vector(DF$Background.ID.1[i])
  v=startsWith(tmp, 'N')
  if(startsWith(tmp, 'N')==TRUE){ DF$Background_Lighting[[i]] = "Anisotropic (direct)" }
  if(startsWith(tmp, 'N')==FALSE){ DF$Background_Lighting[[i]] = "Isotropic (diffuse)" }
}

DF$Background_Lighting<-as.factor(DF$Background_Lighting)  


# Lighting Evolution Population

DF$LightingTreatment <- as.vector(DF$Lighting)

DF$LightingTreatment[DF$LightingTreatment=="1_Natural"] <- Lighting_Levels[1]
DF$LightingTreatment[DF$LightingTreatment=="2_Diffuse"] <- Lighting_Levels[3]
DF$LightingTreatment[DF$LightingTreatment=="3_Hybrid"] <- Lighting_Levels[2]


DF$LightingTreatment<- factor(DF$LightingTreatment,levels = Lighting_Levels)


DF$Background.ID

DF$Background.ID=as.character(DF$Background.ID)
DF$Split_BG_Ldv = ifelse(DF$BG_E15_L_stddv<mean(DF$BG_E15_L_stddv), "low", "high")

DF$Split_BG_Zdv = ifelse(DF$BG_E15_Z_Dev<mean(DF$BG_E15_Z_Dev), "low", "high")


DF$Habitat

DF$HabitatType = as.character(DF$Habitat)
DF$HabitatType=gsub(".*_", "", DF$HabitatType)


unique(DF$HabitatType)

DF$HabitatType = ifelse(startsWith(DF$HabitatType,"grass")==TRUE, "grass" , DF$HabitatType)
DF$HabitatType = ifelse(startsWith(DF$HabitatType,"wood")==TRUE, "wood" , DF$HabitatType)
DF$HabitatType = ifelse(startsWith(DF$HabitatType,"heath")==TRUE, "heath" , DF$HabitatType)
DF$HabitatType = ifelse(startsWith(DF$HabitatType,"gravel")==TRUE, "gravel" , DF$HabitatType)
DF$HabitatType = ifelse(startsWith(DF$HabitatType,"estuary")==TRUE, "estuary" , DF$HabitatType)

DF$Closed = ifelse(startsWith(DF$HabitatType,"wood")==TRUE, "1" , "0")
DF$Shore = ifelse(startsWith(DF$HabitatType,"estuary")==TRUE, "1" , "0")


DF$LightingType = ifelse(startsWith(DF$Lighting,"0:1 Direct")==TRUE, "Direct" , "Diffuse")


DF_Dappled = subset(DF,Dappled==1)
unique(DF_Dappled$Habitat)


DF$Roofed=as.factor(0)
DF$Roofed = ifelse(DF$HabitatType=="03_woodLeaflitter","1", DF$Roofed)
DF$Roofed = ifelse(DF$HabitatType=="04_woodTrail","1", DF$Roofed)
DF$Roofed = ifelse(DF$HabitatType=="07_woodLichen","1", DF$Roofed)
DF$Roofed = ifelse(DF$HabitatType=="10_woodEdgeGrass","1", DF$Roofed)
DF$Roofed = ifelse(DF$HabitatType=="13_heathPond","1", DF$Roofed)
DF$Roofed = ifelse(DF$HabitatType=="15_heathFerns","1", DF$Roofed)
DF$Roofed = ifelse(DF$HabitatType=="19_woodNettle","1", DF$Roofed)


#.............................

#----....[DATA] Measures ----
#.............................


DF$SmotheredShadow = as.factor(DF$SmotheredShadow)
DF$RecievedShadows = as.factor(DF$RecievedShadows)
DF$Dappled = as.factor(DF$Dappled)
DF$Anthropogenic = as.factor(DF$Anthropogenic)


#----............,Measures Rank & Survived ----

DF =  DF%>%
  group_by(UniquePop)  %>% 
  mutate(PopRelativeFitness = mmxn(Fitness))   %>% 
  mutate(PopRankedFitness = rank(Fitness)) 

DF$Survived = ifelse(as.numeric(DF$PopRankedFitness)>12,1,0)


DF =  DF%>%
  group_by(Population)  %>% 
  mutate(TreatRelativeFitness = mmxn(Fitness))   %>% 
  mutate(TreatRankedFitness = rank(Fitness)) 


DF =  DF%>%
  group_by(Habitat)  %>% 
  mutate(HabitatRelativeFitness = mmxn(Fitness))   %>% 
  mutate(HabitatRankedFitness = rank(Fitness)) 


#----............,Measures Averages ----


#----............,Measures Colour Difference ----

#Difference Measures (E2.5, Local)
#............................................
DF$L_Mean_Difference_BG_E2.5 = abs(as.numeric(DF$TG_Render_L_mean) - as.numeric(DF$BG_E2.5_L_mean))
DF$A_Mean_Difference_BG_E2.5 = abs(as.numeric(DF$TG_Render_A_mean) - as.numeric(DF$BG_E2.5_A_mean))                             
DF$B_Mean_Difference_BG_E2.5 = abs(as.numeric(DF$TG_Render_B_mean) - as.numeric(DF$BG_E2.5_B_mean))                             

DF$AB_Mean_Difference_BG_E2.5  = ((DF$A_Mean_Difference_BG_E2.5)^2 + (DF$B_Mean_Difference_BG_E2.5)^2)^0.5
DF$LAB_Mean_Difference_BG_E2.5  = ((DF$AB_Mean_Difference_BG_E2.5 )^2 + (DF$L_Mean_Difference_BG_E2.5 )^2)^0.5


DF$L_Pat_Difference_BG_E2.5_S1 = abs(DF$TG_Render_E_L_Gabor_S1_contrast-DF$BG_E2.5_L_Gabor_S1_contrast)
DF$L_Pat_Difference_BG_E2.5_S2 = abs(DF$TG_Render_E_L_Gabor_S2_contrast-DF$BG_E2.5_L_Gabor_S2_contrast)
DF$L_Pat_Difference_BG_E2.5_S3 = abs(DF$TG_Render_E_L_Gabor_S3_contrast-DF$BG_E2.5_L_Gabor_S3_contrast)
DF$L_Pat_Difference_BG_E2.5_S4 = abs(DF$TG_Render_E_L_Gabor_S4_contrast-DF$BG_E2.5_L_Gabor_S4_contrast)
DF$L_Pat_Difference_BG_E2.5_S5 = abs(DF$TG_Render_E_L_Gabor_S5_contrast-DF$BG_E2.5_L_Gabor_S5_contrast)
DF$L_Pat_Difference_BG_E2.5_S6 = abs(DF$TG_Render_E_L_Gabor_S6_contrast-DF$BG_E2.5_L_Gabor_S6_contrast)


DF$L_Pat_Difference_BG_E2.5   = DF$L_Pat_Difference_BG_E2.5_S1+DF$L_Pat_Difference_BG_E2.5_S2+DF$L_Pat_Difference_BG_E2.5_S3+DF$L_Pat_Difference_BG_E2.5_S4+DF$L_Pat_Difference_BG_E2.5_S5+DF$L_Pat_Difference_BG_E2.5_S6

DF$L_Pat_Difference_BG_E2.5_Alt   = DF$L_Pat_Difference_BG_E2.5_S3+DF$L_Pat_Difference_BG_E2.5_S4+DF$L_Pat_Difference_BG_E2.5_S5+DF$L_Pat_Difference_BG_E2.5_S6


DF$A_Pat_Difference_BG_E2.5_S1 = abs(DF$TG_Render_E_L_Gabor_S1_contrast-DF$BG_E2.5_L_Gabor_S1_contrast)
DF$A_Pat_Difference_BG_E2.5_S2 = abs(DF$TG_Render_E_L_Gabor_S2_contrast-DF$BG_E2.5_L_Gabor_S2_contrast)
DF$A_Pat_Difference_BG_E2.5_S3 = abs(DF$TG_Render_E_L_Gabor_S3_contrast-DF$BG_E2.5_L_Gabor_S3_contrast)
DF$A_Pat_Difference_BG_E2.5_S4 = abs(DF$TG_Render_E_L_Gabor_S4_contrast-DF$BG_E2.5_L_Gabor_S4_contrast)
DF$A_Pat_Difference_BG_E2.5_S5 = abs(DF$TG_Render_E_L_Gabor_S5_contrast-DF$BG_E2.5_L_Gabor_S5_contrast)
DF$A_Pat_Difference_BG_E2.5_S6 = abs(DF$TG_Render_E_L_Gabor_S6_contrast-DF$BG_E2.5_L_Gabor_S6_contrast)


DF$A_Pat_Difference_BG_E2.5   = DF$A_Pat_Difference_BG_E2.5_S1+DF$A_Pat_Difference_BG_E2.5_S2+DF$A_Pat_Difference_BG_E2.5_S3+DF$A_Pat_Difference_BG_E2.5_S4+DF$A_Pat_Difference_BG_E2.5_S5+DF$A_Pat_Difference_BG_E2.5_S6


DF$B_Pat_Difference_BG_E2.5_S1 = abs(DF$TG_Render_E_L_Gabor_S1_contrast-DF$BG_E2.5_L_Gabor_S1_contrast)
DF$B_Pat_Difference_BG_E2.5_S2 = abs(DF$TG_Render_E_L_Gabor_S2_contrast-DF$BG_E2.5_L_Gabor_S2_contrast)
DF$B_Pat_Difference_BG_E2.5_S3 = abs(DF$TG_Render_E_L_Gabor_S3_contrast-DF$BG_E2.5_L_Gabor_S3_contrast)
DF$B_Pat_Difference_BG_E2.5_S4 = abs(DF$TG_Render_E_L_Gabor_S4_contrast-DF$BG_E2.5_L_Gabor_S4_contrast)
DF$B_Pat_Difference_BG_E2.5_S5 = abs(DF$TG_Render_E_L_Gabor_S5_contrast-DF$BG_E2.5_L_Gabor_S5_contrast)
DF$B_Pat_Difference_BG_E2.5_S6 = abs(DF$TG_Render_E_L_Gabor_S6_contrast-DF$BG_E2.5_L_Gabor_S6_contrast)


DF$B_Pat_Difference_BG_E2.5   = DF$B_Pat_Difference_BG_E2.5_S1+DF$B_Pat_Difference_BG_E2.5_S2+DF$B_Pat_Difference_BG_E2.5_S3+DF$B_Pat_Difference_BG_E2.5_S4+DF$B_Pat_Difference_BG_E2.5_S5+DF$B_Pat_Difference_BG_E2.5_S6


#Difference Measures (E15, Global)
#...............................................
DF$L_Mean_Difference_BG_E15 = abs(as.numeric(DF$TG_Render_L_mean) - as.numeric(DF$BG_E15_L_mean))
DF$A_Mean_Difference_BG_E15 = abs(as.numeric(DF$TG_Render_A_mean) - as.numeric(DF$BG_E15_L_mean))                             
DF$B_Mean_Difference_BG_E15 = abs(as.numeric(DF$TG_Render_B_mean) - as.numeric(DF$BG_E15_L_mean))                             

DF$AB_Mean_Difference_BG_E15  = ((DF$A_Mean_Difference_BG_E15)^2 + (DF$B_Mean_Difference_BG_E15)^2)^0.5
DF$LAB_Mean_Difference_BG_E15  = ((DF$AB_Mean_Difference_BG_E15 )^2 + (DF$L_Mean_Difference_BG_E15 )^2)^0.5


DF$L_Pat_Difference_BG_E15_S1 = abs(DF$TG_Render_E_L_Gabor_S1_contrast-DF$BG_E15_L_Gabor_S1_contrast)
DF$L_Pat_Difference_BG_E15_S2 = abs(DF$TG_Render_E_L_Gabor_S2_contrast-DF$BG_E15_L_Gabor_S2_contrast)
DF$L_Pat_Difference_BG_E15_S3 = abs(DF$TG_Render_E_L_Gabor_S3_contrast-DF$BG_E15_L_Gabor_S3_contrast)
DF$L_Pat_Difference_BG_E15_S4 = abs(DF$TG_Render_E_L_Gabor_S4_contrast-DF$BG_E15_L_Gabor_S4_contrast)
DF$L_Pat_Difference_BG_E15_S5 = abs(DF$TG_Render_E_L_Gabor_S5_contrast-DF$BG_E15_L_Gabor_S5_contrast)
DF$L_Pat_Difference_BG_E15_S6 = abs(DF$TG_Render_E_L_Gabor_S6_contrast-DF$BG_E15_L_Gabor_S6_contrast)


DF$L_Pat_Difference_BG_E15   = DF$L_Pat_Difference_BG_E15_S1+DF$L_Pat_Difference_BG_E15_S2+DF$L_Pat_Difference_BG_E15_S3+DF$L_Pat_Difference_BG_E15_S4+DF$L_Pat_Difference_BG_E15_S5+DF$L_Pat_Difference_BG_E15_S6


DF$FactorGeneration = as.factor(DF$Generation)

DF$Fail  <- ifelse(DF$Fitness >=15000 ,1,0)

DF$Fail = as.factor(DF$Fail)


DF$Fitness[DF$Fitness <  300] <-  300 # Introduce minimum fitness, from cheaters


#______________________________________
#~~~~~~~~~~~~~~~~~~~~~~~~~~~~~~~~~~~~~~
#----.......................----
#----(IMP) Subset Sig Improved Pops ---- 
#~~~~~~~~~~~~~~~~~~~~~~~~~~~~~~~~~~~~~~
#______________________________________

#.............................

#----....[Imp] linear model ----
#.............................

#Determine which populations significantly improved by using regression data
#If a populations fitness significantly declined it is excluded.


#Setup Dataframes
TreatmentList <- unique(DF$Population)
Significance <- 1:length(TreatmentList)
T_Value <- 1:length(TreatmentList)
R_Value <- 1:length(TreatmentList)

#Loop linear regression accounting for unique targets and unique background image

for (idx in  1:length(TreatmentList)){
  
  tStr = TreatmentList[[idx]]
  
  SubData <- subset(DF  , Population==tStr)
  
  m1 <- lm(log(Fitness) ~ Generation, data = SubData)
  modelm1 <- summary(m1)
  tablem1 <- coef(modelm1)
  tablem1
  
  
  
  p <- ggplot(SubData, aes(Generation,log(Fitness))) +
    geom_point(shape =21,  alpha = 0.1, size=1)  +
    geom_smooth(method=glm,formula=y~poly(x,1),
                se=TRUE)
  p
  
  p <- p +labs(title=tStr)
  
  #print(p)
  
  t<-
    tablem1%>%
    as_tibble()%>%
    dplyr::select(`t value`)%>%
    pull()
  
  
  
  p<-
    tablem1%>%
    as_tibble()%>%
    dplyr::select(`Pr(>|t|)`)%>%
    pull()
  
  E<-
    tablem1%>%
    as_tibble()%>%
    dplyr::select(`Estimate`)%>%
    pull()
  
  T_Value[[idx ]] <- t[2]
  Significance[[idx ]]<- p[2]
  R_Value[[idx ]]<- E[2]
  
}

Batch_Significance <- data.frame(TreatmentList,Significance,T_Value,R_Value)

sigThreshold = 0.05

Batch_Significance$Positive <- ifelse(Batch_Significance$T_Value >0 ,1,0)
Batch_Significance$Significant <- ifelse(Batch_Significance$Significance <= 0.05,1,0)

Batch_Significance$Improved = Batch_Significance$Positive * Batch_Significance$Significant


#Prepare and Merge

ImproveList <- with(Batch_Significance,data.frame(TreatmentList,Improved,T_Value))
ImproveList<- ImproveList %>% 
  dplyr::rename(
    Population = TreatmentList,
    RevisedImproved = Improved,
    FitnessSlope =T_Value
  )

DF$Population

DF = merge(DF,ImproveList)


#Improved Only

DF_Improved <- subset(DF,RevisedImproved==1)
DF_ImprovedArchive <- subset(DF,Improved=="Yes")


unique(DF_Improved$Population)
unique(DF_ImprovedArchive$Population)


#Normalised Variables
NM_DF = DF

cols <- sapply(NM_DF, is.numeric)
NM_DF[cols] <- lapply(NM_DF[cols], mmxn)

cols <- sapply(NM_DF, is.double)
NM_DF[cols] <- lapply(NM_DF[cols], mmxn)

NM_DF


NM_DF_Improved <- subset(NM_DF,Improved=="Yes")


d=subset(DF, Habitat =="07_woodLichen" & Generation==0)

d$ID


DF =  DF%>%
  group_by(Habitat)  %>% 
  mutate(NoFail = mean(RevisedImproved))  


p1 <- ggplot(subset(DF, Habitat =="07_woodLichen" & Lighting=="1_Natural"), aes(as.factor(Generation), log(Fitness),color=Lighting)) +
  geom_point(alpha=0.5)
p1 


#______________________________________
#~~~~~~~~~~~~~~~~~~~~~~~~~~~~~~~~~~~~~~
#----.......................----
#----(PAPER) Final Plots ---- 
#~~~~~~~~~~~~~~~~~~~~~~~~~~~~~~~~~~~~~~
#______________________________________

DF$L_Mean_Difference_BG_E5

#----..............Data Format----
df1=subset(DF,Generation>=0)


df1$LV =df1$BG_E15_L_stddv
df1$AV = df1$BG_E15_A_stddv
df1$BV = df1$BG_E15_B_stddv
df1$ZV = df1$BG_E15_Z_Dev
  


df1$bgLGT = df1$Background_Lighting
df1$trLGT = df1$LightingTreatment
df1$Fit = mmxn(mmxn(log(df1$Fitness))^0.5)


df1 =  df1%>%
  group_by(Population)  %>% 
  mutate(MeanZV =mean(as.numeric(ZV))) %>% 
  mutate(MeanLV = mean(as.numeric(BG_E15_L_stddv))) %>% 
  mutate(MeanAV = mean(as.numeric(BG_E15_A_stddv))) %>% 
  mutate(MeanBV = mean(as.numeric(BG_E15_B_stddv))) %>% 
  mutate(DevZV = sd(as.numeric(ZV))) %>% 
  mutate(DevLV = sd(as.numeric(BG_E15_L_stddv))) %>% 
  mutate(DevAV = sd(as.numeric(BG_E15_A_stddv))) %>% 
  mutate(DevBV = sd(as.numeric(BG_E15_B_stddv)))
  

df1$MeanCplx = with(df1,zscore(MeanZV)+zscore(MeanLV)+zscore(MeanAV)+zscore(MeanBV))
df1$MeanVar = with(df1,zscore(DevZV)+zscore(DevLV)+zscore(DevAV)+zscore(DevBV))
df1$MeanSCplx = with(df1,zscore(MeanLV)+zscore(MeanAV)+zscore(MeanBV))

df1$Cplx  = with(df1,zscore(ZV)+zscore(LV)+zscore(AV)+zscore(BV))
df1$SCplx  = with(df1,zscore(ZV)+zscore(LV)+zscore(AV)+zscore(BV))


df1$Z_Split= quantcut(df1$MeanZV,2)
levels(df1$Z_Split) <- c("Z_Low", "Z_High")

df1$L_Split= quantcut(df1$MeanLV,2)
levels(df1$L_Split) <- c("L_Low", "L_High")

df1$A_Split= quantcut(df1$MeanAV,2)
levels(df1$A_Split) <- c("A_Low", "A_High")

df1$B_Split= quantcut(df1$MeanBV,2)
levels(df1$B_Split) <- c("B_Low", "B_High")

df1$Cplx_Split= quantcut(df1$MeanCplx,2)
levels(df1$Cplx_Split) <- c("Cplx_Low", "Cplx_High")

df1$SCplx_Split= quantcut(df1$MeanSCplx,2)
levels(df1$SCplx_Split) <- c("SCplx_Low", "SCplx_High")


    
    
    df1$Z_Split2= quantcut(df1$MeanZV,3)
    
    df1$L_Split2= quantcut(df1$MeanLV,3)

    df1$A_Split2= quantcut(df1$MeanAV,3)
 
    df1$B_Split2= quantcut(df1$MeanBV,3)

    df1$Cplx_Split2= quantcut(df1$MeanCplx,3)
    
    df1$Cplx_Split4= quantcut(df1$MeanCplx,4)

    df1$SCplx_Split2= quantcut(df1$MeanSCplx,3)


df1 =  df1%>%
  group_by(trLGT,Generation,Z_Split)  %>% 
  mutate(MeanFitTR = mean(as.numeric(Fitness))) 

df1 =  df1%>%
  group_by(trLGT,Generation,Z_Split)  %>% 
  mutate(MeanFitTR = mean(as.numeric(Fitness))) 

df1 =  df1%>%
  group_by(UniquePop)  %>% 
  mutate(MeanFitPop = mean(as.numeric(Fitness))) 


df1 =  df1%>%
  group_by(Population)  %>% 
  mutate(Mean_BG_LE_S1 = mean(BG_E2.5_L_Gabor_S1_contrast))  %>%  
  mutate(Mean_BG_LE_S2 = mean(BG_E2.5_L_Gabor_S2_contrast))  %>%    
  mutate(Mean_BG_LE_S3 = mean(BG_E2.5_L_Gabor_S3_contrast))  %>%    
  mutate(Mean_BG_LE_S4 = mean(BG_E2.5_L_Gabor_S4_contrast))  %>%    
  mutate(Mean_BG_LE_S5 = mean(BG_E2.5_L_Gabor_S5_contrast))  %>%  
  mutate(Mean_BG_LE_S6 = mean(BG_E2.5_L_Gabor_S6_contrast))  

  


#----___________________________________________________________________----
#----..............1. Camouflage: Fitness, Match and Edge Disruption----
#----___________________________________________________________________----

#----..............Paragraph 1 Text  ...----

#----....................P1.i Data Size ...----

#How many individual players?
      nrow(DF)/24
      
      'N players = 1764'
  
#How many presentations?
      nrow(DF)

      '#N presentations = 42336'

      
      
#----....................P1.ii Fitness ...----


      
#[ STATS Model Fitness - Generation
      mod1 <- lmer(Fit~ zscore(Generation^0.5)+(1|Population)+(1|Habitat), data=df1)
      summary(mod1)
      anova(mod1)
      qqPlot(residuals(mod1))
      hist(residuals(mod1))  
      
      # 
      #   Fixed effects:
      #   Estimate Std. Error        df t value Pr(>|t|)    
      #   (Intercept)            4.485e-01  1.460e-02 2.700e+01   30.73   <2e-16 ***
      #   zscore(Generation^0.5) 3.543e-02  8.778e-04 4.225e+04   40.37   <2e-16 ***
      
      
      
#Starting Fitness
      t = subset(DF, Generation ==0)
      mean(t$Fitness)
      sd(t$Fitness)
      'Start Fit Mean =  923.0779'
      'Start Fit Dev = 1196.655'
      
      median(t$Fitness)
      IQR(t$Fitness)
      'Start Fit Median =649'
      'Start Fit IQR = 667.5'
      
#Ending Fitness
      t = subset(DF, Generation ==20)
      mean(t$Fitness)
      sd(t$Fitness)
      'End Fit = mean 1750.488'
      'End Fit Dev = 2438.083'
      
      median(t$Fitness)
      IQR(t$Fitness)
      'Start Fit Median =1026'
      'Start Fit IQR = 1130.25'
      
#How Many Improved?
      t = subset(DF,RevisedImproved==1 & Generation ==0)
      nrow(t)/24
      'A. 65 out of 84 (76%)'
      
#How Many Improved for each Population?      
      t = subset(DF,RevisedImproved==1 & Generation ==0 & Lighting =="1_Natural") #Direct Lighting
      nrow(t)/24 
      'A. 24 out of 28 (85%)'
     
      t = subset(DF,RevisedImproved==1 & Generation ==0 & Lighting =="2_Diffuse") #Diffuse Lighting
      nrow(t)/24 
      'A. 22 out of 28 (78%)'
      
      t = subset(DF,RevisedImproved==1 & Generation ==0 & Lighting =="3_Hybrid") #Mixed Lighting
      nrow(t)/24 
      'A. 19 out of 28 (67%)'
      

#Fittest Population?
      df1 =  df1%>%
        group_by(Population)  %>% 
        mutate(TreatMeanFitness = mean(Fitness)) %>% 
        mutate(TreatDevFitness = sd(Fitness)) %>% 
        mutate(TreatMedianFitness = median(Fitness)) %>% 
        mutate(TreatIQRFitness = IQR(Fitness)) 

      t = subset(df1, Generation==0 & ShowOrder==1)
      t =  t [order( t$TreatMedianFitness), ]
      t$Population
      t$TreatMeanFitness
      t$TreatDevFitness
      'A. Mean = 3328, Dev = 3542'
      
      t$TreatMedianFitness
      t$TreatIQRFitness
      'Fit Median = 1321.5'
      'Fit IQR = 3119.50'
      
      # Create Categorisation as outliers for failure.
      mn = mean(df1$FitnessSlope)
      sdv = sd(df1$FitnessSlope)
      
      
      df1$OutlierFail = ifelse(df1$FitnessSlope<=(mn-sdv*2.5), 1,0)
      df1$NegativeSlope = ifelse(df1$FitnessSlope<0, 1,0)
      
      nrow(subset(df1,Generation==0 & ShowOrder==1 & NegativeSlope==1))
      '12 with negative slope'
    
#----....................P1.iii Background Variables and Fitness ...----
      
# What is the distribution of fitness slopes for those which failed to improve with the lm      
      p<-ggplot(subset(df1,Generation==0 & ShowOrder==1), aes(x=FitnessSlope, fill=as.factor(RevisedImproved), color=as.factor(RevisedImproved))) +
        geom_histogram(position="identity", alpha=0.5)
      p
      
# Create Categorisation as outliers for failure.

      p<-ggplot(subset(df1,Generation==0 & ShowOrder==1), aes(x=FitnessSlope, fill=as.factor(OutlierFail), color=as.factor(OutlierFail))) +
        geom_histogram(position="identity", alpha=0.5)
      p
      
      p<-ggplot(subset(df1,Generation==0 & ShowOrder==1), aes(x=FitnessSlope, fill=as.factor(NegativeSlope), color=as.factor(NegativeSlope))) +
        geom_histogram(position="identity", alpha=0.5)
      p
      
      
      nrow(subset(df1,Generation==0 & ShowOrder==1 & OutlierFail==1))
      '2 with outliers'
      

      

# Which background feature(s) most explains fitness slope?
      
      
      #L = 2.729e-0
      #L*Gen = 8.943e-04
      mod1 <- lmer(Fit~ zscore(MeanLV)*Generation +(1|NegativeSlope), data=df1) 
      summary(mod1)
      
      qqPlot(residuals(mod1))
      hist(residuals(mod1))  

      
      #A = -1.062e-02 
      #A*Gen= 2.067e-03
      mod1 <- lmer(Fit~ zscore(MeanAV)*Generation +(1|NegativeSlope), data=df1) 
      summary(mod1)
      
      qqPlot(residuals(mod1))
      hist(residuals(mod1))  
      

      #B =  2.071e-02
      #B*Gen= 2.669e-03
      mod1 <- lmer(Fit~ zscore(MeanBV)*Generation +(1|NegativeSlope), data=df1) 
      summary(mod1)
      
      qqPlot(residuals(mod1))
      hist(residuals(mod1)) 
      
      
      #3D = 3.243e-03
      #3D*Gen =  1.791e-03
      mod1 <- lmer(Fit~ zscore(MeanZV)*Generation +(1|NegativeSlope), data=df1) 
      summary(mod1)
      
      qqPlot(residuals(mod1))
      hist(residuals(mod1))       
      
      
      #L + A = 3.432e-02
      #L+A*Gen = 1.269e-02
      mod1 <- lmer(Fit~ zscore(zscore(MeanLV)+zscore(MeanAV))*zscore(Generation)+(1|NegativeSlope), data=df1) 
      summary(mod1)
      
      qqPlot(residuals(mod1))
      hist(residuals(mod1))       
      
      
      #L + B =   1.799e-02
      #L+B*Gen =  1.827e-03 
      mod1 <- lmer(Fit~ zscore(zscore(MeanLV)+zscore(MeanBV))*Generation +(1|NegativeSlope), data=df1) 
      summary(mod1)
      
      qqPlot(residuals(mod1))
      hist(residuals(mod1))       
      
      
      #L + 3D =   1.876e-02
      #L+3D*Gen = 1.972e-03 
      mod1 <- lmer(Fit~ zscore(zscore(MeanLV)+zscore(MeanZV))*Generation+(1|NegativeSlope), data=df1) 
      summary(mod1)
      
      qqPlot(residuals(mod1))
      hist(residuals(mod1))       
      
      
      #A + B =  -4.624e-03 
      #A+B*Gen = 2.363e-03
      mod1 <- lmer(Fit~ zscore(zscore(MeanAV)+zscore(MeanBV))*Generation+(1|NegativeSlope), data=df1) 
      summary(mod1)
      qqPlot(residuals(mod1))
      hist(residuals(mod1))       
      
      
      #A + Z =  -7.885e-03
      #A+Z*Gen = 2.253e-03
      mod1 <- lmer(Fit~ zscore(zscore(MeanAV)+zscore(MeanZV))*Generation +(1|NegativeSlope), data=df1) 
      summary(mod1)
      
      qqPlot(residuals(mod1))
      hist(residuals(mod1))       
      
      
      #B + Z =  -5.461e-04
      #B+Z*Gen =  2.395e-03
      mod1 <- lmer(Fit~ zscore(zscore(MeanBV)+zscore(MeanZV))*Generation +(1|NegativeSlope), data=df1) 
      summary(mod1)
      
      qqPlot(residuals(mod1))
      hist(residuals(mod1))       
      
      
      #L + A + B =   9.529e-03 
      #L + A + B*Gen = 2.271e-03
      mod1 <- lmer(Fit~ zscore(zscore(MeanLV)+zscore(MeanAV)+zscore(MeanBV))*Generation +(1|NegativeSlope), data=df1) 
      summary(mod1)
      
      qqPlot(residuals(mod1))
      hist(residuals(mod1))       
      
      
      #L + A + Z =  8.526e-03
      #L + A + Z*Gen = 2.444e-03 
      mod1 <- lmer(Fit~ zscore(zscore(MeanLV)+zscore(MeanAV)+zscore(MeanZV))*Generation +(1|NegativeSlope), data=df1) 
      summary(mod1)
      
      qqPlot(residuals(mod1))
      hist(residuals(mod1))    
      
      
      #L + B + Z =  1.355e-02
      #L + B + Z*Gen = 2.294e-03
      mod1 <- lmer(Fit~ zscore(zscore(MeanLV)+zscore(MeanBV)+zscore(MeanZV))*Generation +(1|NegativeSlope), data=df1) 
      summary(mod1)
      
      qqPlot(residuals(mod1))
      hist(residuals(mod1))    
      
      
      
      #A + B + Z =  -4.518e-03
      #A + B + Z*Gen =  2.475e-03
      mod1 <- lmer(Fit~ zscore(zscore(MeanAV)+zscore(MeanBV)+zscore(MeanZV))*Generation +(1|NegativeSlope), data=df1) 
      summary(mod1)
      
      qqPlot(residuals(mod1))
      hist(residuals(mod1))    
  
      
      #L + A + B + Z =  6.989e-03 
      #L + A + B + Z*Gen =  2.497e-03
      mod1 <- lmer(Fit~ zscore(zscore(MeanLV)+zscore(MeanAV)+zscore(MeanBV)+zscore(MeanZV))*Generation +(1|NegativeSlope), data=df1) 
      summary(mod1)
      
      qqPlot(residuals(mod1))
      hist(residuals(mod1))       
      
      

      
  
     
#----..............Paragraph 2 Text  ...----
      
#----....................P2.i Fitness with treatments ...----

# Regular Population, Supplementary Material Table SB2      
      
    mod1 <- lmer(Fit~ zscore(Generation^0.5)*trLGT*zscore(MeanZV) +(1|Habitat)+(1|Population), data=subset(df1,NegativeSlope==0)) 
    summary(mod1)
    qqPlot(residuals(mod1))
    hist(residuals(mod1))       
    
    # Fixed effects:
    #   Estimate Std. Error         df t value Pr(>|t|)    
    #  (Intercept)                                             4.713e-01  1.601e-02  3.455e+01  29.433  < 2e-16 ***
    #   zscore(Generation^0.5)                                  4.954e-02  1.594e-03  3.621e+04  31.075  < 2e-16 ***
    #   trLGT1:1 Mixed                                         -2.435e-02  1.037e-02  4.101e+01  -2.347   0.0238 *  
    #   trLGT1:0 Diffuse                                       -2.001e-02  1.086e-02  4.203e+01  -1.843   0.0724 .  
    #  zscore(MeanZV)                                          3.264e-02  1.657e-02  3.319e+01   1.970   0.0573 .  
    #  zscore(Generation^0.5):trLGT1:1 Mixed                  -9.499e-03  2.278e-03  3.621e+04  -4.170 3.06e-05 ***
    #   zscore(Generation^0.5):trLGT1:0 Diffuse                -4.368e-03  2.303e-03  3.621e+04  -1.896   0.0579 .  
    #  zscore(Generation^0.5):zscore(MeanZV)                   9.486e-03  1.608e-03  3.621e+04   5.898 3.72e-09 ***
    #   trLGT1:1 Mixed:zscore(MeanZV)                          -2.164e-02  1.033e-02  4.083e+01  -2.095   0.0424 *  
    #   trLGT1:0 Diffuse:zscore(MeanZV)                        -2.548e-02  1.051e-02  4.116e+01  -2.425   0.0198 *  
    #   zscore(Generation^0.5):trLGT1:1 Mixed:zscore(MeanZV)   -1.653e-03  2.289e-03  3.621e+04  -0.722   0.4703    
    #  zscore(Generation^0.5):trLGT1:0 Diffuse:zscore(MeanZV)  5.194e-03  2.301e-03  3.621e+04   2.257   0.0240 * 

# Mixed Lighting, Supplementary Material Table SB3        

    mod1 <- lmer(Fit~ zscore(Generation^0.5)*bgLGT*zscore(MeanZV) +(1|Habitat), data=subset(df1, trLGT=="1:1 Mixed" & NegativeSlope==0)) 
    summary(mod1)
    qqPlot(residuals(mod1))
    hist(residuals(mod1))       
    

    # Fixed effects:
    #   Estimate Std. Error         df t value Pr(>|t|)    
    # (Intercept)                                                     4.557e-01  1.674e-02  2.237e+01  27.215  < 2e-16 ***
    #   zscore(Generation^0.5)                                          3.979e-02  2.249e-03  1.207e+04  17.690  < 2e-16 ***
    #   bgLGTIsotropic (diffuse)                                       -1.440e-02  3.322e-03  1.207e+04  -4.334 1.48e-05 ***
    #   zscore(MeanZV)                                                  1.425e-02  1.675e-02  2.237e+01   0.851 0.403749    
    # zscore(Generation^0.5):bgLGTIsotropic (diffuse)                 8.167e-04  3.322e-03  1.207e+04   0.246 0.805826    
    # zscore(Generation^0.5):zscore(MeanZV)                           1.104e-02  2.249e-03  1.207e+04   4.909 9.28e-07 ***
    #   bgLGTIsotropic (diffuse):zscore(MeanZV)                        -1.126e-02  3.322e-03  1.207e+04  -3.389 0.000705 ***
    #   zscore(Generation^0.5):bgLGTIsotropic (diffuse):zscore(MeanZV) -7.012e-03  3.322e-03  1.207e+04  -2.110 0.034847 * 
    # 
    

#----....................P2.ii Camouflage Metrics and Treatments ...----


#Luminance Match, Supplementary Material Table SB4    
#...................................................

  mod1 <- lmer(Fit~zscore((L_Mean_Difference_BG_E2.5))*trLGT*zscore(MeanZV)+(1|Habitat)+(1|Population), data=subset(df1,NegativeSlope==0))
  summary(mod1)
  qqPlot(residuals(mod1))
  hist(residuals(mod1))  

  
#Colour Match, Supplementary Material Table SB5  
#...................................................  
  mod1 <- lmer(Fit~zscore((AB_Mean_Difference_BG_E2.5))*trLGT*zscore(MeanZV)+(1|Habitat)+(1|Population), data=subset(df1,NegativeSlope==0))
  summary(mod1)
  qqPlot(residuals(mod1))
  hist(residuals(mod1))  
  
  
#Luminance Contrast Match, Supplementary Material Table SB6 
#...................................................  
  mod1 <- lmer(Fit~zscore((L_Pat_Difference_BG_E2.5))*trLGT*zscore(MeanZV)+(1|Habitat)+(1|Population), data=subset(df1,NegativeSlope==0))
  summary(mod1)
  qqPlot(residuals(mod1))
  hist(residuals(mod1))  
  

  
#GabRat, Supplementary Material Table SB7 
#................................................... 
  mod1 <- lmer(Fit~zscore((Render_L_GabRat))*trLGT*zscore(MeanZV)+(1|Habitat)+(1|Population), data=subset(df1,NegativeSlope==0))
  summary(mod1)
  qqPlot(residuals(mod1))
  hist(residuals(mod1))  
  

  
  #----..............Figure 2 Panels ...----
  
  #----....................Fig B.i) Fitness ...----
  

  p1 <- ggplot(subset(df1,NegativeSlope==0), aes(x=Generation, y=Fitness,col=trLGT,fill=trLGT, linetype=bgLGT)) +
    geom_smooth (aes(),method=glm,formula=y~poly(x,1),alpha=0.25,size=0.75)+
    geom_line (aes(y=MeanFitTR),alpha=0.35)+
    facet_grid(. ~ as.factor(Z_Split))
  p1 + theme_classic() +theme(legend.position = "none") +
    scale_fill_manual(values=Lighting_Palette)+
    scale_color_manual(values=Lighting_Palette)+
    scale_linetype_manual(values=c( "solid","dotted"))+
    coord_cartesian( ylim = c(700, 3000))+
    labs(title="",x ="Generation", y = "Fitness (milliseconds)")+
    theme(text = element_text(size = 14), line = element_line(size=0.4))  + 
    theme(
      axis.text = element_text(family = "sans", size = 12),
      axis.line = element_line(colour = "black", size=0),
      panel.border = element_rect(colour = "black", fill=NA, size=0.4))
  
  
  #----...................Fig .B.ii) Camouflage Metrics ...----
  
  #Top Left, Luminance Match
  
  p1 <- ggplot(subset(df1, NegativeSlope==0), aes(x=(L_Mean_Difference_BG_E2.5), y=TreatRelativeFitness,col=trLGT,fill=trLGT)) +
    geom_smooth (aes(),method=glm,formula=y~poly(x,1),alpha=0.25,size=1)+
    facet_grid(. ~ as.factor(Z_Split))
  
  p1 + theme_classic() +theme(legend.position = "none") +
    scale_fill_manual(values=Lighting_Palette)+
    scale_color_manual(values=Lighting_Palette)+
    scale_linetype_manual(values=c("dotted", "solid"))+
    labs(title="",x ="Luminance Difference", y = "Population Relative Fitness")+
    theme(text = element_text(size = 14), line = element_line(size=0.4))  + 
    theme(
      axis.text = element_text(family = "sans", size = 12),
      axis.line = element_line(colour = "black", size=0),
      panel.border = element_rect(colour = "black", fill=NA, size=0.4))
  

  
  #Top Right, Colour Match 
  
  p1 <- ggplot(subset(df1, NegativeSlope==0), aes(x=(AB_Mean_Difference_BG_E2.5), y=TreatRelativeFitness,col=trLGT,fill=trLGT)) +
    geom_smooth (aes(),method=glm,formula=y~poly(x,1),alpha=0.25,size=1)+
    facet_grid(. ~ as.factor(Z_Split))
  
  p1 + theme_classic() +theme(legend.position = "none") +
    scale_fill_manual(values=Lighting_Palette)+
    scale_color_manual(values=Lighting_Palette)+
    scale_linetype_manual(values=c("dotted", "solid"))+
    labs(title="",x ="Colour Difference", y = "Population Relative Fitness")+
    theme(text = element_text(size = 14), line = element_line(size=0.4))  + 
    theme(
      axis.text = element_text(family = "sans", size = 12),
      axis.line = element_line(colour = "black", size=0),
      panel.border = element_rect(colour = "black", fill=NA, size=0.4))
  
  
  #Bottom Left, Pattern Difference
  
  p1 <- ggplot(subset(df1, NegativeSlope==0), aes(x=(L_Pat_Difference_BG_E2.5), y=TreatRelativeFitness,col=trLGT,fill=trLGT)) +
    geom_smooth (aes(),method=glm,formula=y~poly(x,1),alpha=0.25,size=1)+facet_grid(. ~ as.factor(Z_Split))
  
  p1 + theme_classic() +theme(legend.position = "none") +
    scale_fill_manual(values=Lighting_Palette)+
    scale_color_manual(values=Lighting_Palette)+
    scale_linetype_manual(values=c("dotted", "solid"))+
    labs(title="",x ="Pattern Difference", y = "Population Relative Fitness")+
    theme(text = element_text(size = 14), line = element_line(size=0.4))  + 
    theme(
      axis.text = element_text(family = "sans", size = 12),
      axis.line = element_line(colour = "black", size=0),
      panel.border = element_rect(colour = "black", fill=NA, size=0.4))
  
  

  
  
  #Bottom Right, GabRat
  
  p1 <- ggplot(subset(df1,NegativeSlope==0), aes(x=Render_L_GabRat, y=TreatRelativeFitness,col=trLGT,fill=trLGT)) +
    geom_smooth (aes(),method=glm,formula=y~poly(x,1),alpha=0.25,size=1)+
    facet_grid(. ~ as.factor(Z_Split))
  
  p1 + theme_classic() +theme(legend.position = "none") +
    scale_fill_manual(values=Lighting_Palette)+
    scale_color_manual(values=Lighting_Palette)+
    scale_linetype_manual(values=c("dotted", "solid"))+
    labs(title="",x ="Render L GabRat", y = "Population Relative Fitness")+
    theme(text = element_text(size = 14), line = element_line(size=0.4))  + 
    theme(
      axis.text = element_text(family = "sans", size = 12),
      axis.line = element_line(colour = "black", size=0),
      panel.border = element_rect(colour = "black", fill=NA, size=0.4))
  
  
  

#----___________________________________________________________________----
#----..............2. Colour: Mean and Contrast----
#----___________________________________________________________________----      

#----..............Setup Principle Components of Phenotype  ...----
  
  df1 = df1%>%
    ungroup()
  
  df_pc=df1
  
  df_pc$TG_PC_Mean_L=df_pc$TG_Skin_L_mean
  df_pc$TG_PC_Mean_A=df_pc$TG_Skin_A_mean
  df_pc$TG_PC_Mean_B=df_pc$TG_Skin_B_mean
  
  
  #Luminance
  df_pc$TG_PC_Energy_S1_L=df_pc$TG_Skin_E_L_Gabor_S1_contrast
  df_pc$TG_PC_Aniso_S1_L=df_pc$TG_Skin_E_L_Gabor_S1_peak/df_pc$TG_Skin_E_L_Gabor_S1_contrast
  df_pc$TG_PC_Vert_S1_L=df_pc$TG_Skin_E_L_Gabor_S1_D0-df_pc$TG_Skin_E_L_Gabor_S1_D90
  
  df_pc$TG_PC_Energy_S2_L=df_pc$TG_Skin_E_L_Gabor_S2_contrast
  df_pc$TG_PC_Aniso_S2_L=df_pc$TG_Skin_E_L_Gabor_S2_peak/df_pc$TG_Skin_E_L_Gabor_S2_contrast
  df_pc$TG_PC_Vert_S2_L=df_pc$TG_Skin_E_L_Gabor_S2_D0-df_pc$TG_Skin_E_L_Gabor_S2_D90
  
  df_pc$TG_PC_Energy_S3_L=df_pc$TG_Skin_E_L_Gabor_S3_contrast
  df_pc$TG_PC_Aniso_S3_L=df_pc$TG_Skin_E_L_Gabor_S3_peak/df_pc$TG_Skin_E_L_Gabor_S3_contrast
  df_pc$TG_PC_Vert_S3_L=df_pc$TG_Skin_E_L_Gabor_S3_D0-df_pc$TG_Skin_E_L_Gabor_S3_D90
  
  df_pc$TG_PC_Energy_S4_L=df_pc$TG_Skin_E_L_Gabor_S4_contrast
  df_pc$TG_PC_Aniso_S4_L=df_pc$TG_Skin_E_L_Gabor_S4_peak/df_pc$TG_Skin_E_L_Gabor_S4_contrast
  df_pc$TG_PC_Vert_S4_L=df_pc$TG_Skin_E_L_Gabor_S4_D0-df_pc$TG_Skin_E_L_Gabor_S4_D90
  
  df_pc$TG_PC_Energy_S5_L=df_pc$TG_Skin_E_L_Gabor_S5_contrast
  df_pc$TG_PC_Aniso_S5_L=df_pc$TG_Skin_E_L_Gabor_S5_peak/df_pc$TG_Skin_E_L_Gabor_S5_contrast
  df_pc$TG_PC_Vert_S5_L=df_pc$TG_Skin_E_L_Gabor_S5_D0-df_pc$TG_Skin_E_L_Gabor_S5_D90
  
  df_pc$TG_PC_Energy_S6_L=df_pc$TG_Skin_E_L_Gabor_S6_contrast
  df_pc$TG_PC_Aniso_S6_L=df_pc$TG_Skin_E_L_Gabor_S6_peak/df_pc$TG_Skin_E_L_Gabor_S6_contrast
  df_pc$TG_PC_Vert_S6_L=df_pc$TG_Skin_E_L_Gabor_S6_D0-df_pc$TG_Skin_E_L_Gabor_S6_D90
  
  
  #Blue Yellow
  
  df_pc$TG_PC_Energy_S1_B=df_pc$TG_Skin_E_B_Gabor_S1_contrast
  df_pc$TG_PC_Aniso_S1_B=df_pc$TG_Skin_E_B_Gabor_S1_peak/df_pc$TG_Skin_E_B_Gabor_S1_contrast
  df_pc$TG_PC_Vert_S1_B=df_pc$TG_Skin_E_B_Gabor_S1_D0-df_pc$TG_Skin_E_B_Gabor_S1_D90
  
  df_pc$TG_PC_Energy_S2_B=df_pc$TG_Skin_E_B_Gabor_S2_contrast
  df_pc$TG_PC_Aniso_S2_B=df_pc$TG_Skin_E_B_Gabor_S2_peak/df_pc$TG_Skin_E_B_Gabor_S2_contrast
  df_pc$TG_PC_Vert_S2_B=df_pc$TG_Skin_E_B_Gabor_S2_D0-df_pc$TG_Skin_E_B_Gabor_S2_D90
  
  df_pc$TG_PC_Energy_S3_B=df_pc$TG_Skin_E_B_Gabor_S3_contrast
  df_pc$TG_PC_Aniso_S3_B=df_pc$TG_Skin_E_B_Gabor_S3_peak/df_pc$TG_Skin_E_B_Gabor_S3_contrast
  df_pc$TG_PC_Vert_S3_B=df_pc$TG_Skin_E_B_Gabor_S3_D0-df_pc$TG_Skin_E_B_Gabor_S3_D90
  
  df_pc$TG_PC_Energy_S4_B=df_pc$TG_Skin_E_B_Gabor_S4_contrast
  df_pc$TG_PC_Aniso_S4_B=df_pc$TG_Skin_E_B_Gabor_S4_peak/df_pc$TG_Skin_E_B_Gabor_S4_contrast
  df_pc$TG_PC_Vert_S4_B=df_pc$TG_Skin_E_B_Gabor_S4_D0-df_pc$TG_Skin_E_B_Gabor_S4_D90
  
  df_pc$TG_PC_Energy_S5_B=df_pc$TG_Skin_E_B_Gabor_S5_contrast
  df_pc$TG_PC_Aniso_S5_B=df_pc$TG_Skin_E_B_Gabor_S5_peak/df_pc$TG_Skin_E_B_Gabor_S5_contrast
  df_pc$TG_PC_Vert_S5_B=df_pc$TG_Skin_E_B_Gabor_S5_D0-df_pc$TG_Skin_E_B_Gabor_S5_D90
  
  df_pc$TG_PC_Energy_S6_B=df_pc$TG_Skin_E_B_Gabor_S6_contrast
  df_pc$TG_PC_Aniso_S6_B=df_pc$TG_Skin_E_B_Gabor_S6_peak/df_pc$TG_Skin_E_B_Gabor_S6_contrast
  df_pc$TG_PC_Vert_S6_B=df_pc$TG_Skin_E_B_Gabor_S6_D0-df_pc$TG_Skin_E_B_Gabor_S6_D90
  
  
  
  #Green Red
  
  df_pc$TG_PC_Energy_S1_A=df_pc$TG_Skin_E_A_Gabor_S1_contrast
  df_pc$TG_PC_Aniso_S1_A=df_pc$TG_Skin_E_A_Gabor_S1_peak/df_pc$TG_Skin_E_A_Gabor_S1_contrast
  df_pc$TG_PC_Vert_S1_A=df_pc$TG_Skin_E_A_Gabor_S1_D0-df_pc$TG_Skin_E_A_Gabor_S1_D90
  
  df_pc$TG_PC_Energy_S2_A=df_pc$TG_Skin_E_A_Gabor_S2_contrast
  df_pc$TG_PC_Aniso_S2_A=df_pc$TG_Skin_E_A_Gabor_S2_peak/df_pc$TG_Skin_E_A_Gabor_S2_contrast
  df_pc$TG_PC_Vert_S2_A=df_pc$TG_Skin_E_A_Gabor_S2_D0-df_pc$TG_Skin_E_A_Gabor_S2_D90
  
  df_pc$TG_PC_Energy_S3_A=df_pc$TG_Skin_E_A_Gabor_S3_contrast
  df_pc$TG_PC_Aniso_S3_A=df_pc$TG_Skin_E_A_Gabor_S3_peak/df_pc$TG_Skin_E_A_Gabor_S3_contrast
  df_pc$TG_PC_Vert_S3_A=df_pc$TG_Skin_E_A_Gabor_S3_D0-df_pc$TG_Skin_E_A_Gabor_S3_D90
  
  df_pc$TG_PC_Energy_S4_A=df_pc$TG_Skin_E_A_Gabor_S4_contrast
  df_pc$TG_PC_Aniso_S4_A=df_pc$TG_Skin_E_A_Gabor_S4_peak/df_pc$TG_Skin_E_A_Gabor_S4_contrast
  df_pc$TG_PC_Vert_S4_A=df_pc$TG_Skin_E_A_Gabor_S4_D0-df_pc$TG_Skin_E_A_Gabor_S4_D90
  
  df_pc$TG_PC_Energy_S5_A=df_pc$TG_Skin_E_A_Gabor_S5_contrast
  df_pc$TG_PC_Aniso_S5_A=df_pc$TG_Skin_E_A_Gabor_S5_peak/df_pc$TG_Skin_E_A_Gabor_S5_contrast
  df_pc$TG_PC_Vert_S5_A=df_pc$TG_Skin_E_A_Gabor_S5_D0-df_pc$TG_Skin_E_A_Gabor_S5_D90
  
  df_pc$TG_PC_Energy_S6_A=df_pc$TG_Skin_E_A_Gabor_S6_contrast
  df_pc$TG_PC_Aniso_S6_A=df_pc$TG_Skin_E_A_Gabor_S6_peak/df_pc$TG_Skin_E_A_Gabor_S6_contrast
  df_pc$TG_PC_Vert_S6_A=df_pc$TG_Skin_E_A_Gabor_S6_D0-df_pc$TG_Skin_E_A_Gabor_S6_D90
  

  
  
  my_df = subset(df_pc)
  pcData=my_df
  pcData <- dplyr::select(pcData, starts_with("TG_PC_") )
  myPr <- prcomp(pcData, scale = TRUE, center = TRUE) #prcomp
  
  summary(myPr)
  

  # PC1     PC2    PC3     PC4     PC5     PC6     PC7    PC8     PC9   PC10    PC11
  # Standard deviation     2.9875 2.35846 2.0201 1.87597 1.81019 1.78717 1.42570 1.3901 1.33889 1.2678 1.20720
  # Proportion of Variance 0.1566 0.09759 0.0716 0.06174 0.05749 0.05603 0.03566 0.0339 0.03145 0.0282 0.02557
  # Cumulative Proportion  0.1566 0.25416 0.3258 0.38750 0.44499 0.50102 0.53668 0.5706 0.60203 0.6302 0.65580
  # 
   #PCs which explain > 50% of variance, PC1-PC9
  
  loadings <- myPr$rotation
  loadings
  
  
  PC1 = loadings[,1]
  PC2 = loadings[,2]
  PC3 = loadings[,3]
  PC4 = loadings[,4]
  PC5 = loadings[,5]
  PC6 = loadings[,6]
  
  PCs = loadings[,1:6]
  PCsRank = sqrt(PCs^2)
  PCsRank 
  
  PCsRank<- as.data.frame(PCsRank)
  PCsRank$PC1R <- as.numeric(PCsRank$PC1)
  PCsRank$PC2R <- as.numeric(PCsRank$PC2)
  PCsRank$PC3R <- as.numeric(PCsRank$PC3)
  PCsRank$PC4R <- as.numeric(PCsRank$PC4)
  PCsRank$PC5R <- as.numeric(PCsRank$PC5)
  PCsRank$PC6R <- as.numeric(PCsRank$PC6)
  
  PCSRANKED <-cbind(PCs[,1:6],PCsRank[,7:12])
  
  #Loadings
  #.....................................................................................
  # Get Most Important for PC1
  PCsRanked1 <- PCSRANKED[order(PCSRANKED$PC1R, decreasing = TRUE),]  
  PCsRanked1 <- cbind(PCsRanked1[1])
  PCsRanked1
  
  
  # Get Most Important for PC2
  PCsRanked2 <- PCSRANKED[order(PCSRANKED$PC2R, decreasing = TRUE),]  
  PCsRanked2 <- cbind(PCsRanked2[2])
  PCsRanked2
  
  
  # Get Most Important for PC3
  PCsRanked3 <- PCSRANKED[order(PCSRANKED$PC3R, decreasing = TRUE),]  
  PCsRanked3 <- cbind(PCsRanked3[3])
  PCsRanked3
  
  # Get Most Important for PC4
  PCsRanked4 <- PCSRANKED[order(PCSRANKED$PC4R, decreasing = TRUE),]  
  PCsRanked4 <- cbind(PCsRanked4[4])
  PCsRanked4
  
  # Get Most Important for PC5
  PCsRanked5 <- PCSRANKED[order(PCSRANKED$PC5R, decreasing = TRUE),]  
  PCsRanked5 <- cbind(PCsRanked5[5])
  PCsRanked5
  
  
  
  # Get Most Important for PC6
  PCsRanked6 <- PCSRANKED[order(PCSRANKED$PC6R, decreasing = TRUE),]  
  PCsRanked6 <- cbind(PCsRanked6[6])
  PCsRanked6
  
  CompTemp = cbind(my_df, myPr$x[,1:6])  # This will create the new dataset which
  
  CompTemp$trLGT2 = fct_rev(CompTemp$trLGT)
  
  plotData = subset(CompTemp,Generation==0 | Generation ==20)
  
  p1 <- 
    ggplot(subset(  plotData ,NegativeSlope==0 & Generation==20), aes(x =PC1, y =PC2,col=trLGT)) +
    geom_point() +
    facet_grid(. ~Z_Split) 
  
  p1 + theme_classic() +theme(legend.position = "none") +
    scale_fill_manual(values=Lighting_Palette2)+
    scale_color_manual(values=Lighting_Palette2)+
    scale_linetype_manual(values=c("dotted", "solid"))+
    theme(text = element_text(size = 14), line = element_line(size=0.5))  + 
    theme(
      axis.text = element_text(family = "sans", size = 12),
      axis.line = element_line(colour = "black", size=0),
      panel.border = element_rect(colour = "black", fill=NA, size=0.5))
  
  
  
  
  
  
#----..............Figure 3 Panels ...----
  
  
#----....................Fig A.ii) Colour Space ...----  
  
#----.................................Column 1 ...----
  
  
  #Column 1, Row 1, Direct Lighting Generation 0

  ColData=subset(DF,Lighting=="1_Natural" & Generation==0 & NoFail==1)

  LAB_Col <- with(ColData,LAB(TG_Skin_L_mean,TG_Skin_A_mean,TG_Skin_B_mean))
  p1<-ggplot(subset(ColData), aes(x=TG_Skin_A_mean, y=TG_Skin_B_mean,col=Lighting,fill=Lighting)) +
    geom_vline(xintercept = 0, linetype = "dashed",alpha=0.5) +
    geom_point(size=5, shape=16,  aes(colour=hex(LAB_Col,fix = TRUE))) +
    scale_color_identity()
  
  p1=p1 + theme_classic() +theme(legend.position = "none")+ 
    labs(title="",x ="", y = "")+
    scale_x_continuous(breaks = seq(-50, 50, by = 20))+
    scale_y_continuous(breaks = seq(-10, 80, by = 20))+
    coord_cartesian(xlim = c(-50, 50), ylim = c(-20, 80))+
    theme(text = element_text(size = 0))  +
    theme(text = element_text(size = 14), line = element_line(size=0.4))  + 
    theme(
      axis.text = element_text(family = "sans", size = 0),
      axis.line = element_line(colour = "black", size=0),
      panel.border = element_rect(colour = "black", fill=NA, size=0.4))
  p1+ theme(panel.grid.major = element_line(color = "gray90", linetype = 1))
  
  
  #Column 1, Row 2, Mixed Lighting Generation 0
  
  ColData=subset(DF,Lighting=="3_Hybrid" & Generation==0 & NoFail==1)
  
  LAB_Col <- with(ColData,LAB(TG_Skin_L_mean,TG_Skin_A_mean,TG_Skin_B_mean))
  p1<-ggplot(subset(ColData), aes(x=TG_Skin_A_mean, y=TG_Skin_B_mean,col=Lighting,fill=Lighting))+
    geom_vline(xintercept = 0, linetype = "dashed",alpha=0.5) +
    geom_point(size=5, shape=16,  aes(colour=hex(LAB_Col,fix = TRUE))) +
    scale_color_identity()
  
  p1=p1 + theme_classic() +theme(legend.position = "none")+ 
    labs(title="",x ="", y = "")+
    scale_x_continuous(breaks = seq(-50, 50, by = 20))+
    scale_y_continuous(breaks = seq(-10, 80, by = 20))+
    coord_cartesian(xlim = c(-50, 50), ylim = c(-20, 80))+
    theme(text = element_text(size = 0))  +
    theme(text = element_text(size = 14), line = element_line(size=0.4))  + 
    theme(
      axis.text = element_text(family = "sans", size = 0),
      axis.line = element_line(colour = "black", size=0),
      panel.border = element_rect(colour = "black", fill=NA, size=0.4))
  p1+ theme(panel.grid.major = element_line(color = "gray90", linetype = 1)) 
  
  #Column 1, Row 3, Diffuse Lighting Generation 0
  
  ColData=subset(DF,Lighting=="2_Diffuse" & Generation==0 & NoFail==1)
  
  LAB_Col <- with(ColData,LAB(TG_Skin_L_mean,TG_Skin_A_mean,TG_Skin_B_mean))
  p1<-ggplot(subset(ColData), aes(x=TG_Skin_A_mean, y=TG_Skin_B_mean,col=Lighting,fill=Lighting))+
    geom_vline(xintercept = 0, linetype = "dashed",alpha=0.5) +
    geom_point(size=5, shape=16,  aes(colour=hex(LAB_Col,fix = TRUE))) +
    scale_color_identity()
  
  p1=p1 + theme_classic() +theme(legend.position = "none")+ 
    labs(title="",x ="", y = "")+
    scale_x_continuous(breaks = seq(-50, 50, by = 20))+
    scale_y_continuous(breaks = seq(-10, 80, by = 20))+
    coord_cartesian(xlim = c(-50, 50), ylim = c(-20, 80))+
    theme(text = element_text(size = 0))  +
    theme(text = element_text(size = 14), line = element_line(size=0.4))  + 
    theme(
      axis.text = element_text(family = "sans", size = 0),
      axis.line = element_line(colour = "black", size=0),
      panel.border = element_rect(colour = "black", fill=NA, size=0.4))
  p1+ theme(panel.grid.major = element_line(color = "gray90", linetype = 1))
  
  
 
  
  #----.................................Column 2 ...----
  
  
  #Column 2, Row 1, Direct Lighting Generation 20
  
  ColData=subset(DF,Lighting=="1_Natural" & Generation==20 & NoFail==1)
  
  LAB_Col <- with(ColData,LAB(TG_Skin_L_mean,TG_Skin_A_mean,TG_Skin_B_mean))
  p1<-ggplot(subset(ColData), aes(x=TG_Skin_A_mean, y=TG_Skin_B_mean,col=Lighting,fill=Lighting))+
    geom_vline(xintercept = 0, linetype = "dashed",alpha=0.5) +
    geom_point(size=5, shape=16,  aes(colour=hex(LAB_Col,fix = TRUE))) +
    scale_color_identity()
  
  p1=p1 + theme_classic() +theme(legend.position = "none")+ 
    labs(title="",x ="", y = "")+
    scale_x_continuous(breaks = seq(-50, 50, by = 20))+
    scale_y_continuous(breaks = seq(-10, 80, by = 20))+
    coord_cartesian(xlim = c(-50, 50), ylim = c(-20, 80))+
    theme(text = element_text(size = 0))  +
    theme(text = element_text(size = 14), line = element_line(size=0.4))  + 
    theme(
      axis.text = element_text(family = "sans", size = 0),
      axis.line = element_line(colour = "black", size=0),
      panel.border = element_rect(colour = "black", fill=NA, size=0.4))
  p1+ theme(panel.grid.major = element_line(color = "gray90", linetype = 1))
  
  
  #Column 2, Row 2, Mixed Lighting Generation 20
  
  ColData=subset(DF,Lighting=="3_Hybrid" & Generation==20 & NoFail==1)
  
  LAB_Col <- with(ColData,LAB(TG_Skin_L_mean,TG_Skin_A_mean,TG_Skin_B_mean))
  p1<-ggplot(subset(ColData), aes(x=TG_Skin_A_mean, y=TG_Skin_B_mean,col=Lighting,fill=Lighting))+
    geom_vline(xintercept = 0, linetype = "dashed",alpha=0.5) +
    geom_point(size=5, shape=16,  aes(colour=hex(LAB_Col,fix = TRUE))) +
    scale_color_identity()
  
  p1=p1 + theme_classic() +theme(legend.position = "none")+ 
    labs(title="",x ="", y = "")+
    scale_x_continuous(breaks = seq(-50, 50, by = 20))+
    scale_y_continuous(breaks = seq(-10, 80, by = 20))+
    coord_cartesian(xlim = c(-50, 50), ylim = c(-20, 80))+
    theme(text = element_text(size = 0))  +
    theme(text = element_text(size = 14), line = element_line(size=0.4))  + 
    theme(
      axis.text = element_text(family = "sans", size = 0),
      axis.line = element_line(colour = "black", size=0),
      panel.border = element_rect(colour = "black", fill=NA, size=0.4))
  p1+ theme(panel.grid.major = element_line(color = "gray90", linetype = 1))
  
  
  #Column 2, Row 3, Diffuse Lighting Generation 20
  
  ColData=subset(DF,Lighting=="2_Diffuse" & Generation==20 & NoFail==1)
  
  LAB_Col <- with(ColData,LAB(TG_Skin_L_mean,TG_Skin_A_mean,TG_Skin_B_mean))
  p1<-ggplot(subset(ColData), aes(x=TG_Skin_A_mean, y=TG_Skin_B_mean,col=Lighting,fill=Lighting))+
    geom_vline(xintercept = 0, linetype = "dashed",alpha=0.5) +
    geom_point(size=5, shape=16,  aes(colour=hex(LAB_Col,fix = TRUE))) +
    scale_color_identity()
  
  p1=p1 + theme_classic() +theme(legend.position = "none")+ 
    labs(title="",x ="", y = "")+
    scale_x_continuous(breaks = seq(-50, 50, by = 20))+
    scale_y_continuous(breaks = seq(-10, 80, by = 20))+
    coord_cartesian(xlim = c(-50, 50), ylim = c(-20, 80))+
    theme(text = element_text(size = 0))  +
    theme(text = element_text(size = 14), line = element_line(size=0.4))  + 
    theme(
      axis.text = element_text(family = "sans", size = 0),
      axis.line = element_line(colour = "black", size=0),
      panel.border = element_rect(colour = "black", fill=NA, size=0.4))
  p1+ theme(panel.grid.major = element_line(color = "gray90", linetype = 1))
  
  
  
  
  #----.................................Column 3 ...----
  
  
  #Column 3, Row 1, Direct Lighting Background
  
  ColData=subset(DF,Lighting=="1_Natural" & Generation==20 & NoFail==1)
  
  LAB_Col <- with(ColData,LAB(BG_E2.5_L_mean,BG_E2.5_A_mean,BG_E2.5_B_mean))
  p1<-ggplot(subset(ColData), aes(x=BG_E2.5_A_mean, y=BG_E2.5_B_mean,col=Lighting,fill=Lighting))+
    geom_vline(xintercept = 0, linetype = "dashed",alpha=0.5) +
    geom_point(size=5, shape=16,  aes(colour=hex(LAB_Col,fix = TRUE))) +
    scale_color_identity()
  
  p1=p1 + theme_classic() +theme(legend.position = "none")+ 
    labs(title="",x ="", y = "")+
    scale_x_continuous(breaks = seq(-50, 50, by = 20))+
    scale_y_continuous(breaks = seq(-10, 80, by = 20))+
    coord_cartesian(xlim = c(-50, 50), ylim = c(-20, 80))+
    theme(text = element_text(size = 0))  +
    theme(text = element_text(size = 14), line = element_line(size=0.4))  + 
    theme(
      axis.text = element_text(family = "sans", size = 0),
      axis.line = element_line(colour = "black", size=0),
      panel.border = element_rect(colour = "black", fill=NA, size=0.4))
  p1+ theme(panel.grid.major = element_line(color = "gray90", linetype = 1))
  
  
  #Column 2, Row 2, Mixed Lighting Background
  
  ColData=subset(DF,Lighting=="3_Hybrid" & Generation==20 & NoFail==1)
  
  LAB_Col <- with(ColData,LAB(BG_E2.5_L_mean,BG_E2.5_A_mean,BG_E2.5_B_mean))
  p1<-ggplot(subset(ColData), aes(x=BG_E2.5_A_mean, y=BG_E2.5_B_mean,col=Lighting,fill=Lighting))+
    geom_vline(xintercept = 0, linetype = "dashed",alpha=0.5) +
    geom_point(size=5, shape=16,  aes(colour=hex(LAB_Col,fix = TRUE))) +
    scale_color_identity()
  
  
  p1=p1 + theme_classic() +theme(legend.position = "none")+ 
    labs(title="",x ="", y = "")+
    scale_x_continuous(breaks = seq(-50, 50, by = 20))+
    scale_y_continuous(breaks = seq(-10, 80, by = 20))+
    coord_cartesian(xlim = c(-50, 50), ylim = c(-20, 80))+
    theme(text = element_text(size = 0))  +
    theme(text = element_text(size = 14), line = element_line(size=0.4))  + 
    theme(
      axis.text = element_text(family = "sans", size = 0),
      axis.line = element_line(colour = "black", size=0),
      panel.border = element_rect(colour = "black", fill=NA, size=0.4))
  p1+ theme(panel.grid.major = element_line(color = "gray90", linetype = 1))
  
  
  
  #Column 2, Row 3, Diffuse Lighting Background
  
  ColData=subset(DF,Lighting=="2_Diffuse" & Generation==20 & NoFail==1)
  
  LAB_Col <- with(ColData,LAB(BG_E2.5_L_mean,BG_E2.5_A_mean,BG_E2.5_B_mean))
  p1<-ggplot(subset(ColData), aes(x=BG_E2.5_A_mean, y=BG_E2.5_B_mean,col=Lighting,fill=Lighting))+
    geom_vline(xintercept = 0, linetype = "dashed",alpha=0.5) +
    geom_point(size=5, shape=16,  aes(colour=hex(LAB_Col,fix = TRUE))) +
    scale_color_identity()
  
  p1=p1 + theme_classic() +theme(legend.position = "none")+ 
    labs(title="",x ="", y = "")+
    scale_x_continuous(breaks = seq(-50, 50, by = 20))+
    scale_y_continuous(breaks = seq(-10, 80, by = 20))+
    coord_cartesian(xlim = c(-50, 50), ylim = c(-20, 80))+
    theme(text = element_text(size = 0))  +
    theme(text = element_text(size = 14), line = element_line(size=0.4))  + 
    theme(
      axis.text = element_text(family = "sans", size = 0),
      axis.line = element_line(colour = "black", size=0),
      panel.border = element_rect(colour = "black", fill=NA, size=0.4))
  p1+ theme(panel.grid.major = element_line(color = "gray90", linetype = 1))
  
  
  #----....................Fig B.ii) PC1 ...----
  
  CompTemp$trLGT2 = fct_rev(CompTemp$trLGT)
  
  plotData = subset(CompTemp,Generation==0 | Generation ==20)
  
  p1 <- 
    ggplot(subset(  plotData ,NegativeSlope==0 & Generation==20), aes(x = -PC1, y =trLGT2)) +
    geom_violin(aes(fill = trLGT2),alpha=0.5) +
    geom_boxplot(aes(fill = trLGT2),alpha=0.5,width=0.35) +
    #geom_line(data= subset(CompTemp,RevisedImproved==1 & Generation==20 & ShowOrder==1),aes(x = avgPC1, group=HabitatType),alpha=0.5) +
    facet_grid(. ~Z_Split) 
  
  p1 + theme_classic() +theme(legend.position = "none") +
    scale_fill_manual(values=Lighting_Palette2)+
    scale_color_manual(values=Lighting_Palette2)+
    scale_linetype_manual(values=c("dotted", "solid"))+
    theme(text = element_text(size = 14), line = element_line(size=0.5))  + 
    theme(
      axis.text = element_text(family = "sans", size = 12),
      axis.line = element_line(colour = "black", size=0),
      panel.border = element_rect(colour = "black", fill=NA, size=0.5))
  
  
  
  plotData = subset(CompTemp,Generation==0 | Generation ==20)
  
  p1 <- 
    ggplot(subset(  plotData ,NegativeSlope==0 & Generation==0), aes(x = -PC1, y =trLGT2)) +
    geom_violin(aes(fill = trLGT2),alpha=0.5) +
    geom_boxplot(aes(fill = trLGT2),alpha=0.5,width=0.35) +
    #geom_line(data= subset(CompTemp,RevisedImproved==1 & Generation==20 & ShowOrder==1),aes(x = avgPC1, group=HabitatType),alpha=0.5) +
    facet_grid(. ~Z_Split) 
  
  p1 + theme_classic() +theme(legend.position = "none") +
    scale_fill_manual(values=Lighting_Palette2)+
    scale_color_manual(values=Lighting_Palette2)+
    scale_linetype_manual(values=c("dotted", "solid"))+
    theme(text = element_text(size = 14), line = element_line(size=0.5))  + 
    theme(
      axis.text = element_text(family = "sans", size = 12),
      axis.line = element_line(colour = "black", size=0),
      panel.border = element_rect(colour = "black", fill=NA, size=0.5))
  

#----..............Paragraph 1 Text  ...----
  
#----....................P1.i How much variance do the PCs explain ...----      
      
summary(myPr)      
  
# Sum of PCs 1-6 = 47% 

# PC1 explains 15.66% of variance
# PC2 explains 9.76% of variance
# PC3 explains 7.16% of variance     
# PC4 explains 6.17% of variance
  
       
#----....................P1.ii Which variables contribute the most to PCs 1-4 ...----      

  # Get Most Important for PC1, Supplementary Material Table SB8    
  
      PCsRanked1 # Gives ranked order of PC1
      
      #PC1 is mostly low blue-yellow and low luminance contrast at larger spatial scales, and higher verticality
  
          #As majority of values are negative we will used the - of the PC.    
  

  
  # Get Most Important for PC2, Supplementary Material Table SB8    

      PCsRanked2 # Gives ranked order of PC2
      
      #PC2 is mostly verticality in colour at medium scales
    
  
  # Get Most Important for PC3, Supplementary Material Table SB9    

        PCsRanked3 # Gives ranked order of PC3
        
        #PC3 is mostly contrast in the red-green channels.
        
    
  
  # Get Most Important for PC4, Supplementary Material Table SB9    
  
      PCsRanked4 # Gives ranked order of PC4
      
      #PC4 is mostly lower verticality at the smallest scales
      
          #As majority of values are negative we will used the - of the PC.    
          
          #PC4 is mostly higher verticality at the smallest scales
          
  
#----....................P1.iii How does Population effect PCs ...----    
  
  #PCs are inverted if they are explained predominantley by negative variables
  
      
  # PC1, Supplementary Material Table SB10    
  mod1 <- lmer(zscore(-PC1) ~ zscore(Generation^0.5)*trLGT*zscore(MeanZV) +(1|Population)+(1|Habitat), data=subset(CompTemp,NegativeSlope==0))
  summary(mod1)
  qqPlot(residuals(mod1)) 
  hist(resid(mod1))  

  # PC2
  mod1 <- lmer(zscore(PC2) ~ zscore(Generation^0.5)*trLGT*zscore(MeanZV) +(1|Population)+(1|Habitat), data=subset(CompTemp,NegativeSlope==0))
  summary(mod1)
  qqPlot(residuals(mod1)) 
  hist(resid(mod1))  
  
  # PC3
  mod1 <- lmer(zscore(PC3) ~ zscore(Generation^0.5)*trLGT*zscore(MeanZV) +(1|Population)+(1|Habitat), data=subset(CompTemp,NegativeSlope==0))
  summary(mod1)
  qqPlot(residuals(mod1)) 
  hist(resid(mod1))  
  
  # PC4
  mod1 <- lmer(zscore(PC4) ~ zscore(Generation^0.5)*trLGT*zscore(MeanZV) +(1|Population)+(1|Habitat), data=subset(CompTemp,NegativeSlope==0))
  summary(mod1)
  qqPlot(residuals(mod1)) 
  hist(resid(mod1))  
  
  
  p1 <- ggplot(subset(CompTemp,NegativeSlope==0), aes(x=Generation, y=-PC1,col=trLGT,fill=trLGT)) +
    geom_smooth (aes(),method=glm,formula=y~poly(x,2),alpha=0.25,size=0.75)+
    facet_grid(. ~ as.factor(Z_Split))
  p1 + theme_classic() +theme(legend.position = "none") +
    scale_fill_manual(values=Lighting_Palette)+
    scale_color_manual(values=Lighting_Palette)

  p1 <- ggplot(subset(CompTemp,NegativeSlope==0), aes(x=Generation, y=PC2,col=trLGT,fill=trLGT)) +
    geom_smooth (aes(),method=glm,formula=y~poly(x,2),alpha=0.25,size=0.75)+
  facet_grid(. ~ as.factor(Z_Split))
  p1 + theme_classic() +theme(legend.position = "none") +
    scale_fill_manual(values=Lighting_Palette)+
    scale_color_manual(values=Lighting_Palette)
  
  
  p1 <- ggplot(subset(CompTemp,NegativeSlope==0), aes(x=Generation, y=PC4,col=trLGT,fill=trLGT)) +
    geom_smooth (aes(),method=glm,formula=y~poly(x,1),alpha=0.25,size=0.75)+
    facet_grid(. ~ as.factor(Z_Split))
  p1 + theme_classic() +theme(legend.position = "none") +
    scale_fill_manual(values=Lighting_Palette)+
    scale_color_manual(values=Lighting_Palette)
  
  
  
  
  
  
  
  #______________________________________
  #~~~~~~~~~~~~~~~~~~~~~~~~~~~~~~~~~~~~~~
  #----.......................----
  #----(SPLIT) Split Data Pattern Comp ---- 
  #~~~~~~~~~~~~~~~~~~~~~~~~~~~~~~~~~~~~~~
  #______________________________________
  
  DF_S1 = df1
  DF_S2 = df1
  DF_S3 = df1
  DF_S4 = df1
  DF_S5 = df1
  DF_S6 = df1
  
  
  #S1
  DF_S1$Scale= 1
  DF_S1$ScaleRelative= 1/32
  DF_S1$ScaleMM= 1/32*30
  
  DF_S1$BG_L_Energy = DF$BG_E2.5_L_Gabor_S1_contrast
  DF_S1$BG_A_Energy = DF$BG_E2.5_A_Gabor_S1_contrast
  DF_S1$BG_B_Energy = DF$BG_E2.5_B_Gabor_S1_contrast
  
  DF_S1$TG_L_Energy = DF$TG_Render_E_L_Gabor_S1_contrast
  DF_S1$TG_A_Energy = DF$TG_Render_E_A_Gabor_S1_contrast
  DF_S1$TG_B_Energy = DF$TG_Render_E_B_Gabor_S1_contrast
  
  DF_S1$TG_Sk_L_Energy = DF$TG_Skin_E_L_Gabor_S1_contrast
  
  #S2
  DF_S2$Scale= 2
  DF_S2$ScaleRelative= 1/16
  DF_S2$ScaleMM= 1/16*30
  
  DF_S2$BG_L_Energy = DF$BG_E2.5_L_Gabor_S2_contrast
  DF_S2$BG_A_Energy = DF$BG_E2.5_A_Gabor_S2_contrast
  DF_S2$BG_B_Energy = DF$BG_E2.5_B_Gabor_S2_contrast
  
  DF_S2$TG_L_Energy = DF$TG_Render_E_L_Gabor_S2_contrast
  DF_S2$TG_A_Energy = DF$TG_Render_E_A_Gabor_S2_contrast
  DF_S2$TG_B_Energy = DF$TG_Render_E_B_Gabor_S2_contrast
  
  DF_S2$TG_Sk_L_Energy = DF$TG_Skin_E_L_Gabor_S2_contrast
  
  #S3
  DF_S3$Scale= 3
  DF_S3$ScaleRelative= 1/8
  DF_S3$ScaleMM= 1/8*30
  
  DF_S3$BG_L_Energy = DF$BG_E2.5_L_Gabor_S3_contrast
  DF_S3$BG_A_Energy = DF$BG_E2.5_A_Gabor_S3_contrast
  DF_S3$BG_B_Energy = DF$BG_E2.5_B_Gabor_S3_contrast
  
  DF_S3$TG_L_Energy = DF$TG_Render_E_L_Gabor_S3_contrast
  DF_S3$TG_A_Energy = DF$TG_Render_E_A_Gabor_S3_contrast
  DF_S3$TG_B_Energy = DF$TG_Render_E_B_Gabor_S3_contrast
  
  DF_S3$TG_Sk_L_Energy = DF$TG_Skin_E_L_Gabor_S3_contrast
  
  #S4
  DF_S4$Scale= 4
  DF_S4$ScaleRelative= 1/4
  DF_S4$ScaleMM= 1/4*30
  
  DF_S4$BG_L_Energy = DF$BG_E2.5_L_Gabor_S4_contrast
  DF_S4$BG_A_Energy = DF$BG_E2.5_A_Gabor_S4_contrast
  DF_S4$BG_B_Energy = DF$BG_E2.5_B_Gabor_S4_contrast
  
  DF_S4$TG_L_Energy = DF$TG_Render_E_L_Gabor_S4_contrast
  DF_S4$TG_A_Energy = DF$TG_Render_E_A_Gabor_S4_contrast
  DF_S4$TG_B_Energy = DF$TG_Render_E_B_Gabor_S4_contrast
  
  DF_S4$TG_Sk_L_Energy = DF$TG_Skin_E_L_Gabor_S4_contrast
  
  #S5
  DF_S5$Scale= 5
  DF_S5$ScaleRelative= 1/2
  DF_S5$ScaleMM= 1/2*30
  
  DF_S5$BG_L_Energy = DF$BG_E2.5_L_Gabor_S5_contrast
  DF_S5$BG_A_Energy = DF$BG_E2.5_A_Gabor_S5_contrast
  DF_S5$BG_B_Energy = DF$BG_E2.5_B_Gabor_S5_contrast
  
  DF_S5$TG_L_Energy = DF$TG_Render_E_L_Gabor_S5_contrast
  DF_S5$TG_A_Energy = DF$TG_Render_E_A_Gabor_S5_contrast
  DF_S5$TG_B_Energy = DF$TG_Render_E_B_Gabor_S5_contrast
  
  DF_S5$TG_Sk_L_Energy = DF$TG_Skin_E_L_Gabor_S5_contrast
  
  #S6
  DF_S6$Scale= 6
  DF_S6$ScaleRelative= 1/1
  DF_S6$ScaleMM= 1/1*30
  
  DF_S6$BG_L_Energy = DF$BG_E2.5_L_Gabor_S6_contrast
  DF_S6$BG_A_Energy = DF$BG_E2.5_A_Gabor_S6_contrast
  DF_S6$BG_B_Energy = DF$BG_E2.5_B_Gabor_S6_contrast
  
  DF_S6$TG_L_Energy = DF$TG_Render_E_L_Gabor_S6_contrast
  DF_S6$TG_A_Energy = DF$TG_Render_E_A_Gabor_S6_contrast
  DF_S6$TG_B_Energy = DF$TG_Render_E_B_Gabor_S6_contrast
  
  DF_S6$TG_Sk_L_Energy = DF$TG_Skin_E_L_Gabor_S6_contrast
  
  
  #Combine
  
  DF_Com = rbind(DF_S1,DF_S2,DF_S3,DF_S4,DF_S5,DF_S6)
  
  
  
  
  

  
  
#----___________________________________________________________________----
#----..............3. Pattern: Gradient and Shape----
#----___________________________________________________________________----      
  

  
#----..............Figure 4 Panels ...----
  
  
#----....................Fig A.i) Countershading and Population ...----  
  
  
  p1 <- ggplot(df1, aes(x=Generation, y=Render_L_Gradient_Natural-Natural_Sphere_L_Gradient,col=trLGT)) +
    geom_smooth ( data=subset(df1),aes(),method=glm,formula=y~poly(x,1),alpha=0.25)+
    facet_grid(. ~ as.factor(Z_Split))
  
  p1 + theme_classic() +theme(legend.position = "none") +
    scale_fill_manual(values=Lighting_Palette)+
    scale_color_manual(values=Lighting_Palette)+
    scale_linetype_manual(values=c("dotted", "solid"))+
    labs(title="",x ="Generation", y = "Render - Grey Target Gradient")+
    theme(text = element_text(size = 14), line = element_line(size=1.4))  + 
    theme(
      axis.text = element_text(family = "sans", size = 12),
      axis.line = element_line(colour = "black", size=0),
      panel.border = element_rect(colour = "black", fill=NA, size=1.2))
  
  
  
  
#----....................Fig A.ii) Countershading and Substrate ...----  
      
  p1 <- ggplot(subset(df1, bgLGT != "Isotropic (diffuse)"  & NegativeSlope==0), aes(x=Render_L_Gradient_Natural-Natural_Sphere_L_Gradient, y=TreatRelativeFitness,col=trLGT)) +
    geom_smooth (method=glm,formula=y~poly(x,1),alpha=0.25, size =1)+
    facet_grid(. ~ as.factor(DominantSubstrate))
  
  p1 + theme_classic() +theme(legend.position = "none") +
    scale_fill_manual(values=Lighting_Palette)+
    scale_color_manual(values=Lighting_Palette)+
    scale_linetype_manual(values=c("solid", "dashed"))+
    scale_x_continuous(breaks = seq(-0.2, 0.2, by = 0.2))+
    labs(title="",x ="Gradient Difference", y = "Treat Relative Fitness")+
    geom_vline(xintercept = 0, linetype = "dotted",alpha=0.5) +
    theme(text = element_text(size = 14), line = element_line(size=0.4))  + 
    theme(
      strip.background = element_rect(size = 0.4),
      axis.text = element_text(family = "sans", size = 12),
      axis.line = element_line(colour = "black", size=0),
      panel.border = element_rect(colour = "black", fill=NA, size=0.4))
  
  

  df1$Mean_BG_LE_S1
  
  
#----....................Fig B.i)  Pattern Contrast Scales ...----  
  Hex_Direct2 = "#D5C15F"
  Hex_Diffuse2 = "#7294C2"
 
  DF_T=subset(DF_Com,Generation==20 )
  p1 <- ggplot(subset( DF_T, NegativeSlope==0), aes(x=ScaleRelative*30, y=BG_L_Energy,col=LightingTreatment)) +
    geom_smooth (data=subset(DF_Com,Generation==0 &NegativeSlope==0),aes(y=(TG_Sk_L_Energy)),method=glm,formula=y~poly(x,1),col=Hex_Red,fill=Hex_Red,se=FALSE,linetype=3)+
    geom_smooth (data=subset(DF_Com,Generation==0 &NegativeSlope==0 & bgLGT=="Anisotropic (direct)"),aes(y=(BG_L_Energy)),method=glm,formula=y~poly(x,1),col= Hex_Direct2,fill= Hex_Direct2,se=FALSE,linetype=2,alpha=0.1, size=0.5)+
    geom_smooth (data=subset(DF_Com,Generation==0 &NegativeSlope==0 & bgLGT =="Isotropic (diffuse)"),aes(y=(BG_L_Energy)),method=glm,formula=y~poly(x,1),col= Hex_Diffuse2,fill= Hex_Diffuse2,se=FALSE,linetype=2,alpha=0.1, size=0.5)+
    geom_smooth(method=glm,formula=y~poly(x,1), 
                aes( y=(TG_Sk_L_Energy))
                , alpha=0.15)+

    scale_x_log10()+
    scale_y_log10()+
    facet_grid(. ~ Z_Split)
  
  p1 + theme_classic() +theme(legend.position = "none") +
    scale_fill_manual(values=Lighting_Palette)+
    scale_color_manual(values=Lighting_Palette)+
    scale_linetype_manual(values=c("dotted", "solid"))+
    labs(title="",x ="Log Scale mm", y = "Log Pattern Contrast")+
    theme(text = element_text(size = 14), line = element_line(size=0.4))  + 
    theme(
      axis.text = element_text(family = "sans", size = 12),
      axis.line = element_line(colour = "black", size=0),
      panel.border = element_rect(colour = "black", fill=NA, size=0.4))
  
  
  
  
  
  
  
  
  
  
  

#----....................Fig B.ii) Pattern Direction and Directionality ...----  
  
  df1$A=df1$TG_Skin_E_L_Gabor_S1_peak/df1$TG_Skin_E_L_Gabor_S1_contrast
  df1$E=df1$TG_Skin_E_L_Gabor_S1_contrast
  df1$V=df1$TG_Skin_E_L_Gabor_S1_D0-df1$TG_Skin_E_L_Gabor_S1_D90
  
  
  
  plotdata <- subset(df1, Generation==0 & NegativeSlope==0 | Generation ==20 & NegativeSlope==0)
  
  plotdata <- plotdata  %>%
    group_by(Population,Generation)  %>%
    mutate(VectorX1 = mean(E))   %>%
    mutate(VectorY1 = mean(A))   %>%
    mutate(vert = mean(V))  
  
  plotdata$VectorX2 = with(plotdata, ifelse(vert<=-0.0,VectorX1+0.001, VectorX1))
  plotdata$VectorY2 = with(plotdata, ifelse(vert>=0.0,VectorY1+0.05, VectorY1))
  
  plotdata$VectorX3 = with(plotdata, ifelse(vert<=-0.0,VectorX1-0.001, VectorX1))
  plotdata$VectorY3 = with(plotdata, ifelse(vert>=0.0,VectorY1-0.05, VectorY1))
  
  plotdata$Angled = 0
  plotdata$Angled = with(plotdata, ifelse(vert<=-0.0,1, plotdata$Angled ))
  plotdata$Angled = with(plotdata, ifelse(vert>=0.0,1, plotdata$Angled ))
  
  

  
  p1 <- ggplot(plotdata, aes(x=(E), y=(A),col=trLGT)) +
    geom_smooth ( data=subset(plotdata,Generation==20 & RevisedImproved==1),aes(),method=glm,formula=y~poly(x,1),alpha=0.15)+
    geom_smooth (data=subset(plotdata,Generation==0 & RevisedImproved==1 ),aes(),method=glm,formula=y~poly(x,1),col=Hex_Red,fill=Hex_Red,se=FALSE,linetype=3)+
    
    geom_segment( data=subset(plotdata,Angled==1 & Generation==20 & RevisedImproved==1 & ShowOrder==1),
                  aes(x= VectorX3, y=VectorY3, xend = VectorX2, yend = VectorY2, col=trLGT),
                  arrow=arrow(type = 'closed', length = unit(0.1, "inches")), size=0.2, alpha=0.5) +
    geom_point( data=subset(plotdata,Generation==20 & RevisedImproved==1 & ShowOrder==1),
                aes(x= VectorX1, y=VectorY1, col=trLGT), alpha=0.5) +
    
    facet_grid(. ~ as.factor(Z_Split))
  
  p1 + theme_classic() +theme(legend.position = "none") +
    scale_fill_manual(values=Lighting_Palette)+
    scale_color_manual(values=Lighting_Palette)+
    scale_linetype_manual(values=c("dotted", "solid"))+
    labs(title="",x ="Pattern Contrast 1/32", y = "Pattern Anisotropy 1/32")+
    theme(text = element_text(size = 14), line = element_line(size=0.4))  + 
    theme(
      axis.text = element_text(family = "sans", size = 12),
      axis.line = element_line(colour = "black", size=0),
      panel.border = element_rect(colour = "black", fill=NA, size=0.4))
  
  
#----..............Paragraph 1 Text  ...----  
  
#----....................P1.i Countershading with Population ...----       

 # Supplementary Material Table SB11    
# Gradient Shift
df1$CounterShade = zscore( df1$Render_L_Gradient_Natural- df1$Natural_Sphere_L_Gradient)
mod1 <- lmer(zscore(Render_L_Gradient_Natural)~Generation*trLGT*zscore(MeanZV)+(1|Habitat)+(1|Population), data=subset(df1,NegativeSlope==0))
summary(mod1)
qqPlot(residuals(mod1))
hist(residuals(mod1))  


#----....................P1.ii Substrates ...----       

mod1 <- lmer(Fit~PatternSkin_L_Gradient:DominantSubstrate+(1|Population)+(1|Habitat), data=subset(df1, bgLGT != "Isotropic (diffuse)" & NegativeSlope==0))
summary(mod1)
qqPlot(residuals(mod1))
hist(residuals(mod1))  

emmeans_results <- emmeans(mod1, pairwise ~ PatternSkin_L_Gradient:DominantSubstrate,pbkrtest.limit = 21483)
summary(emmeans_results )

  
  
#----..............Paragraph 2 Text  ...----  

#----....................P2.i Which scale most explains maculation? ...----         

#Create measure of genetic contrast
df1$Gene_Pattern_Contrast = with(df1, abs(col_mac_lmv-(col_top_lmv+col_bot_lmv)/2))
df1$Gene_Gradient_Contrast = with(df1, abs(col_top_lmv-col_bot_lmv))


mod1 <- lm(zscore(Gene_Pattern_Contrast^0.5)~zscore(TG_Skin_E_L_Gabor_S1_deviation), data=subset(df1))
mod2 <- lm(zscore(Gene_Pattern_Contrast^0.5)~zscore(TG_Skin_E_L_Gabor_S2_deviation), data=subset(df1))
mod3 <- lm(zscore(Gene_Pattern_Contrast^0.5)~zscore(TG_Skin_E_L_Gabor_S3_deviation), data=subset(df1))
mod4 <- lm(zscore(Gene_Pattern_Contrast^0.5)~zscore(TG_Skin_E_L_Gabor_S4_deviation), data=subset(df1))
mod5 <- lm(zscore(Gene_Pattern_Contrast^0.5)~zscore(TG_Skin_E_L_Gabor_S5_deviation), data=subset(df1))
mod6 <- lm(zscore(Gene_Pattern_Contrast^0.5)~zscore(TG_Skin_E_L_Gabor_S6_deviation), data=subset(df1))

aic1 <- AIC(mod1)
aic2 <- AIC(mod2)
aic3 <- AIC(mod3)
aic4 <- AIC(mod4)
aic5 <- AIC(mod5)
aic6 <- AIC(mod6)


aic_values <- c(mod1=aic1,mod2=aic2,mod3=aic3,mod4=aic4,mod5=aic5,mod6=aic6)
aic_values

#Lowest AIC belongs to models 1 

# As scale increases gradient contrast is more explained.


#----....................P2.ii Table for correlation ...----      

DF_Com$ED=abs(DF_Com$TG_Sk_L_Energy-DF_Com$BG_L_Energy)
mod1 <- lmer(zscore((TG_Sk_L_Energy^0.5))~
               Scale*LightingTreatment*zscore(MeanZV)+(1|Population)+(1|Habitat)+(1|UniquePhoto), 
             data=subset(DF_Com, Generation==20 & NegativeSlope==0))
summary(mod1)
qqPlot(residuals(mod1))
hist(residuals(mod1))  


mod1 <- lmer(zscore((TG_Sk_L_Energy^0.5))~
               Scale*LightingTreatment*zscore(MeanZV)+(1|Population)+(1|Habitat)+(1|UniquePhoto), 
             data=subset(DF_Com, Generation==20 & NegativeSlope==0))
summary(mod1)
qqPlot(residuals(mod1))
hist(residuals(mod1))  


#----....................P2.iii Pattern Directionality ...----   


# Correlations between maculation contrast, directionality and verticality

    
    mod1 <- lmer(zscore(log(A))~zscore(log(E))+(1|Population)+(1|Habitat), data=subset(df1, NegativeSlope==0 & Generation==0))
    summary(mod1)
    qqPlot(residuals(mod1))
    hist(residuals(mod1))  
    
    mod1 <- lmer(zscore((V))~zscore(log(E))+(1|Population)+(1|Habitat), data=subset(df1, NegativeSlope==0 & Generation==0))
    summary(mod1)
    qqPlot(residuals(mod1))
    hist(residuals(mod1))  
    
    mod1 <- lmer(zscore((V))~zscore(log(A))+(1|Population)+(1|Habitat), data=subset(df1, NegativeSlope==0 & Generation==0))
    summary(mod1)
    qqPlot(residuals(mod1))
    hist(residuals(mod1))  

# How does evolution effect directionality, Supplementary Material Table SB12
    mod1 <- lmer(zscore(log(A))~zscore(Generation)*trLGT*zscore(MeanZV)+(1|Population)+(1|Habitat), data=subset(df1, NegativeSlope==0 ))
    summary(mod1)
    qqPlot(residuals(mod1))
    hist(residuals(mod1))  

# How does evolution effect verticality, Supplementary Material Table SB13 

    mod1 <- lmer(zscore(V)~zscore(Generation)*trLGT*zscore(MeanZV)+(1|Population)+(1|Habitat), data=subset(df1, NegativeSlope==0 ))
    summary(mod1)
    qqPlot(residuals(mod1))
    hist(residuals(mod1))  


    
    
  
  
  
#----___________________________________________________________________----
#----..............4. Specular Effects: Edge enhancement and glossiness----
#----___________________________________________________________________----      


#----..............Edge Enhancement and Glossiness  ...----
    

df.tg.eeF
eeD = df.tg.eeF

eeD$TG_Skin_EE_stddv

eeD=merge(df1,eeD)

eeD$LightingTreatment <- as.vector(eeD$Lighting)

eeD$LightingTreatment[eeD$LightingTreatment=="1_Natural"] <- Lighting_Levels[1]
eeD$LightingTreatment[eeD$LightingTreatment=="2_Diffuse"] <- Lighting_Levels[3]
eeD$LightingTreatment[eeD$LightingTreatment=="3_Hybrid"] <- Lighting_Levels[2]


eeD$LightingTreatment<- factor(eeD$LightingTreatment,levels = Lighting_Levels)

eeD$EdgeDiff = (eeD$TG_Skin_EE_stddv-eeD$TG_Skin_NE_stddv)
eeD$EdgeDiff_S1 = (eeD$TG_Skin_EE_stddv_S1-eeD$TG_Skin_NE_stddv_S1)
eeD$EdgeDiff_S2 = (eeD$TG_Skin_EE_stddv_S2-eeD$TG_Skin_NE_stddv_S2)
eeD$EdgeDiff_S3 = (eeD$TG_Skin_EE_stddv_S3-eeD$TG_Skin_NE_stddv_S3)
eeD$EdgeDiff_S4 = (eeD$TG_Skin_EE_stddv_S4-eeD$TG_Skin_NE_stddv_S4)
eeD$EdgeDiff_Sum = eeD$EdgeDiff_S1+eeD$EdgeDiff_S2+eeD$EdgeDiff_S3+eeD$EdgeDiff_S4

eeD$EdgeShift = (eeD$TG_Skin_EE_mean-eeD$TG_Skin_NE_mean)


eeD$ZV = eeD$BG_E15_Z_GAB_all

eeD =  eeD%>%
  group_by(Population)  %>% 
  mutate(MeanLV = mean(as.numeric(BG_E15_L_stddv))) %>% 
  mutate(MeanAV = mean(as.numeric(BG_E15_A_stddv))) %>% 
  mutate(MeanBV = mean(as.numeric(BG_E15_B_stddv)))
  
  
eeD1 = eeD
eeD2 = eeD
eeD3 = eeD

eeD1$Scale = 1/64
eeD2$Scale = 1/32
eeD3$Scale = 1/16


eeD1$Energy = eeD$EdgeDiff_S1
eeD2$Energy = eeD$EdgeDiff_S2
eeD3$Energy = eeD$EdgeDiff_S3

eeD_Combine = rbind(eeD1,eeD2,eeD3)


df1$Glossiness= df1$Gloss_Whole_Mean
df1$RelativeGlossiness= df1$Gloss_Whole_Mean/df1$GlossMx_Whole_Mean
df1$PotentialGlossiness= df1$GlossMx_Whole_Mean

df1$GlossContrast= abs(df1$Gloss_Back_Mean-df1$Gloss_Mac_Mean)

df1$Glossy = ifelse(df1$Gloss_Whole_Mean>10,1,0)

t=subset(df1,Generation==20 & ShowOrder==1)
ordered_indices <- order(t$Glossiness)
ordered_data <- t[ordered_indices, ]
ordered_data$Population

#----..............Figure 5 Panels, NB figure 5 was ...----
#----....................Fig A.i) Edge Enhancement and Population ...----  

p1 <- ggplot(subset(eeD,NegativeSlope==0), aes(x=Generation^0.5, y=EdgeDiff,col=trLGT)) +
  geom_smooth (method=glm,formula=y~poly(x,1))+
  facet_grid(. ~ as.factor(Z_Split))

p1 + theme_classic() +theme(legend.position = "none") +
  scale_fill_manual(values=Lighting_Palette)+
  scale_color_manual(values=Lighting_Palette)+
  scale_linetype_manual(values=c("dotted", "solid"))+
  labs(title="",x ="Generation", y = "Edge Difference")+
  theme(text = element_text(size = 14), line = element_line(size=0.4))  + 
  theme(
    axis.text = element_text(family = "sans", size = 12),
    axis.line = element_line(colour = "black", size=0),
    panel.border = element_rect(colour = "black", fill=NA, size=0.4))


#----....................Fig B.i) Glossiness and Population ...----  

p1 <- ggplot(subset(df1, NegativeSlope==0), aes(x=Generation, y=Glossiness,col=trLGT,fill=trLGT)) +
  geom_smooth(method=glm,formula=y~poly(x,1))
p1 + theme_classic() +theme(legend.position = "none") +
  scale_fill_manual(values=Lighting_Palette)+
  scale_color_manual(values=Lighting_Palette)+
  labs(title="",x ="Generation", y = "Relative Glossiness")+
  theme(text = element_text(size = 14), line = element_line(size=0.4))  + 
  theme(
    axis.text = element_text(family = "sans", size = 12),
    axis.line = element_line(colour = "black", size=0),
    panel.border = element_rect(colour = "black", fill=NA, size=0.4))


p1 <- ggplot(subset(df1, NegativeSlope==0), aes(x=Generation, y=RelativeGlossiness,col=trLGT,fill=trLGT)) +
  geom_smooth(method=glm,formula=y~poly(x,1))+
  facet_grid(. ~ as.factor(Z_Split))
p1 + theme_classic() +theme(legend.position = "none") +
  scale_fill_manual(values=Lighting_Palette)+
  scale_color_manual(values=Lighting_Palette)+
  labs(title="",x ="Generation", y = "Relative Glossiness")+
  theme(text = element_text(size = 14), line = element_line(size=0.4))  + 
  theme(
    axis.text = element_text(family = "sans", size = 12),
    axis.line = element_line(colour = "black", size=0),
    panel.border = element_rect(colour = "black", fill=NA, size=0.4))


#----..............Paragraph 1 Text  ...----  

#----....................P1.i Edge Enhancement ...----        
    
      eeD$Maculated = ifelse(eeD$EdgeDiff==0 & eeD$eem_int_lvl !=0, 0, 1)

# Create Categorisation as outliers for failure.
mn = mean(eeD$FitnessSlope)
sdv = sd(eeD$FitnessSlope)
eeD$EdgeOutlier = ifelse(abs(eeD$EdgeDiff)>=mn+sdv*2,1,0)


#Which Scales did EE change? a. all of them
    mod1 <- lmer(zscore(EdgeDiff_S1)~zscore(Generation^0.5)*trLGT*zscore(MeanZV)+(1|Population)+(1|Habitat), data=subset(eeD, NegativeSlope==0 & eeD$EdgeOutlier==0))
    summary(mod1)
    qqPlot(residuals(mod1))
    hist(residuals(mod1))  #YES
    
    mod1 <- lmer(zscore(EdgeDiff_S2)~zscore(Generation^0.5)*trLGT*zscore(MeanZV)+(1|Population)+(1|Habitat), data=subset(eeD, NegativeSlope==0 & eeD$EdgeOutlier==0))
    summary(mod1)
    qqPlot(residuals(mod1))
    hist(residuals(mod1)) #YES  
    
    mod1 <- lmer(zscore(EdgeDiff_S3)~zscore(Generation^0.5)*trLGT*zscore(MeanZV)+(1|Population)+(1|Habitat), data=subset(eeD, NegativeSlope==0 & eeD$EdgeOutlier==0))
    summary(mod1)
    qqPlot(residuals(mod1))
    hist(residuals(mod1))   #YES 
    
    mod1 <- lmer(zscore(EdgeDiff_S4)~zscore(Generation^0.5)*trLGT*zscore(MeanZV)+(1|Population)+(1|Habitat), data=subset(eeD, NegativeSlope==0 & eeD$EdgeOutlier==0))
    summary(mod1)
    qqPlot(residuals(mod1))
    hist(residuals(mod1))   #YES 
    
     #All Scales, Data used, Supplementary Material Table SB2   
    #,,,,,,,,,,,,,,,,,,,,,,,,,,,,,,,,,,,,,,,,,,,,,,,,,,,,,,,,,,,,,
    mod1 <- lmer(zscore(EdgeDiff)~zscore(Generation^0.5)*trLGT*zscore(MeanZV)+(1|Population)+(1|Habitat), data=subset(eeD, NegativeSlope==0  & eeD$EdgeOutlier==0 ))
    summary(mod1)
    qqPlot(residuals(mod1))
    hist(residuals(mod1))   #YES 
    

    
    
    
#----..............Paragraph 2 Text  ...----      
    
#----....................P2.i Glossiness ...----        
    
#Relative Gloss Evolution
    
    #Relative Gloss, Used in Paper
    mod1 <- lmer(zscore(RelativeGlossiness^0.5)~zscore(Generation^0.5)*trLGT*zscore(MeanZV)+(1|Population)+(1|Habitat), data=subset(df1, NegativeSlope==0))
    summary(mod1)
    qqPlot(residuals(mod1))
    hist(residuals(mod1))  
    
    mod1 <- lmer(zscore(Glossiness^0.5)~zscore(Generation^0.5)*trLGT*zscore(MeanZV)+(1|Population)+(1|Habitat), data=subset(df1, NegativeSlope==0))
    summary(mod1)
    qqPlot(residuals(mod1))
    hist(residuals(mod1))  
    
    
    
#Factors effecting potential Glossiness
    mod1 <- lmer(zscore(PotentialGlossiness^0.5)~zscore(TG_Skin_L_mean)*bgLGT*zscore(Generation)+(1|Population)+(1|Habitat), data=subset(df1, NegativeSlope==0))
    summary(mod1)
    qqPlot(residuals(mod1))
    hist(residuals(mod1))  
    
    
    mod1 <- lmer(zscore(GlossMx_Whole_Dev^0.5)~zscore(TG_Skin_L_mean)*bgLGT*zscore(Generation)+(1|Population)+(1|Habitat), data=subset(df1, NegativeSlope==0))
    summary(mod1)
    qqPlot(residuals(mod1))
    hist(residuals(mod1)) 
    
    
#Does countershading negatively impact glossiness?
    mod1 <- lmer(zscore(Glossiness^0.5)~zscore(Generation^0.5)*zscore(PatternSkin_L_Gradient)*zscore(MeanZV)+(1|Population)+(1|Habitat), data=subset(df1, NegativeSlope==0))
    summary(mod1)
    qqPlot(residuals(mod1))
    hist(residuals(mod1))  
    
    mod1 <- lmer(zscore(Glossiness^0.5)~zscore(Generation^0.5)*zscore(PatternSkin_L_Gradient)*trLGT+(1|Population)+(1|Habitat), data=subset(df1, NegativeSlope==0))
    summary(mod1)
    qqPlot(residuals(mod1))
    hist(residuals(mod1))  
    
        
    
#______________________________________
#~~~~~~~~~~~~~~~~~~~~~~~~~~~~~~~~~~~~~~
#----.......................----
#----(Supplement) Plots ---- 
#~~~~~~~~~~~~~~~~~~~~~~~~~~~~~~~~~~~~~~
#______________________________________
    
    #----___________________________________________________________________----
    #----..............1. CounterShading----
    #----___________________________________________________________________
    
    #----..............Being Darker effects self-shadow Gradient  ...----  

    p1 <- ggplot(subset(df1,Generation==0 | Generation==20), aes(x=TG_Skin_L_mean, y=Render_L_Gradient_Natural,col=trLGT)) +
      geom_smooth ( data=subset(df1,Generation==0 | Generation==20),aes(),method=glm,formula=y~poly(x,2),alpha=0.25)+
      facet_grid(. ~ as.factor(Generation))
    p1 + theme_classic() +
      scale_fill_manual(values=Lighting_Palette)+
      scale_color_manual(values=Lighting_Palette)+
      scale_linetype_manual(values=c("dotted", "solid"))+
      labs(title="",x ="Target-Skin", y = "Render Gradient")+
      theme(text = element_text(size = 14), line = element_line(size=1.4))  + 
      theme(
        axis.text = element_text(family = "sans", size = 12),
        axis.line = element_line(colour = "black", size=0),
        panel.border = element_rect(colour = "black", fill=NA, size=1.2))
    
    
    mod1 <- lmer(Render_L_Gradient_Natural~zscore(Generation^0.5)*zscore(TG_Skin_L_mean)*trLGT+(1|Population)+(1|Habitat), data=subset(df1, NegativeSlope==0))
    summary(mod1)
    qqPlot(residuals(mod1))
    hist(residuals(mod1))  
    
    
    #----..............Gradient effects contrast match  ...----  
    
    p1 <- ggplot(subset(df1,Generation==0 | Generation==20), aes(x=PatternSkin_L_Gradient, y=L_Pat_Difference_BG_E2.5 ,col=trLGT,linetype=as.factor(Generation))) +
      geom_smooth ( data=subset(df1,Generation==0 | Generation==20),aes(),method=glm,formula=y~poly(x,2),alpha=0.25)+
      facet_grid(. ~ as.factor(trLGT))
    p1 + theme_classic() +
      scale_fill_manual(values=Lighting_Palette)+
      scale_color_manual(values=Lighting_Palette)+
      scale_linetype_manual(values=c("dotted", "solid"))+
      labs(title="",x ="Target-Skin Gradient", y = "Contrast Difference")+
      theme(text = element_text(size = 14), line = element_line(size=1.4))  + 
      theme(
        axis.text = element_text(family = "sans", size = 12),
        axis.line = element_line(colour = "black", size=0),
        panel.border = element_rect(colour = "black", fill=NA, size=1.2))
    
    
    
    mod1 <- lmer(zscore((L_Pat_Difference_BG_E2.5)^0.5)~zscore(Generation^0.5)*poly(zscore(PatternSkin_L_Gradient),2)*bgLGT+(1|Population)+(1|Habitat), data=subset(df1, NegativeSlope==0))
    summary(mod1)
    qqPlot(residuals(mod1))
    hist(residuals(mod1))  
    
    
    
    #----___________________________________________________________________----
    #----..............2. Glossiness----
    #----___________________________________________________________________
    
    #----..............Being Darker effects glossiness  ...----  
    
    p1 <- ggplot(subset(df1,Generation==0 | Generation==20), aes(x=TG_Skin_L_mean, y=(Gloss_Back_Mean/gls_lvl_bkg)+(Gloss_Mac_Mean/gls_lvl_mac),col=trLGT)) +
      geom_smooth (aes(),method=glm,formula=y~poly(x,1),alpha=0.25)+
      facet_grid(. ~ as.factor(Generation))
    p1 + theme_classic() +
      scale_fill_manual(values=Lighting_Palette)+
      scale_color_manual(values=Lighting_Palette)+
      scale_linetype_manual(values=c("dotted", "solid"))+
      labs(title="",x ="Un-rendered Pattern Luminance", y = "Glossiness")+
      theme(text = element_text(size = 14), line = element_line(size=1.4))  + 
      theme(
        axis.text = element_text(family = "sans", size = 12),
        axis.line = element_line(colour = "black", size=0),
        panel.border = element_rect(colour = "black", fill=NA, size=1.2))
    
    
    mod1 <- lmer(zscore(Gloss_Whole_Mean^0.5)~zscore(Generation^0.5)*zscore(TG_Skin_L_mean)*trLGT+(1|Population)+(1|Habitat), data=subset(df1, NegativeSlope==0))
    summary(mod1)
    qqPlot(residuals(mod1))
    hist(residuals(mod1))  
    
    #----___________________________________________________________________----
    #----..............3. Offset Fitness----
    #----___________________________________________________________________
    
    
    p1 <- ggplot(subset(df1,NegativeSlope==0 ), aes(x=TG_Render_L_mean - BG_E2.5_L_mean, y=Fitness/1000,col=trLGT,fill=trLGT)) +
      geom_smooth (aes(),method=glm,formula=y~poly(x,2),alpha=0.25,size=0.75)+
      facet_grid(. ~ as.factor(Z_Split))
    p1 + theme_classic() +theme(legend.position = "none") +
      scale_fill_manual(values=Lighting_Palette)+
      scale_color_manual(values=Lighting_Palette)+
      scale_linetype_manual(values=c( "solid","dotted"))+
      coord_cartesian( ylim = c(0.5, 2.5))+
      labs(title="",x ="Target L - Background L", y = "Capture Time (seconds)")+
      theme(text = element_text(size = 14), line = element_line(size=0.4))  + 
      theme(
        axis.text = element_text(family = "sans", size = 12),
        axis.line = element_line(colour = "black", size=0),
        panel.border = element_rect(colour = "black", fill=NA, size=0.4))
    

    df1$Lvar = quantcut(df1$BG_E15_L_Gabor_S6_contrast,3)
    
    p1 <- ggplot(subset(df1,NegativeSlope==0 ), aes(x=TG_Render_L_mean - BG_E2.5_L_mean, y=Fitness/1000,col=Z_Split,fill=Z_Split)) +
      geom_smooth (aes(),method=glm,formula=y~poly(x,2),alpha=0.25,size=0.75)
    p1 + theme_classic()  +
      scale_linetype_manual(values=c( "solid","dotted"))+
      coord_cartesian( ylim = c(0.5, 2.5))+
      labs(title="",x ="Target L - Background L", y = "Capture Time (seconds)")+
      theme(text = element_text(size = 14), line = element_line(size=0.4))  + 
      theme(
        axis.text = element_text(family = "sans", size = 12),
        axis.line = element_line(colour = "black", size=0),
        panel.border = element_rect(colour = "black", fill=NA, size=0.4))
    
    
mod1 <- lmer(Fit~trLGT*poly(zscore(TG_Render_L_mean - BG_E2.5_L_mean),2)+(1|Population)+(1|Habitat), data=subset(df1, NegativeSlope==0))
    summary(mod1)
    qqPlot(residuals(mod1))
    hist(residuals(mod1))  
    
    
  
    df1$NegPos = ifelse(df1$TG_Render_L_mean - df1$BG_E2.5_L_mean<0,"Negative","Positive")
    mod1 <- lmer(Fit~NegPos*trLGT*BG_E15_L_Gabor_S6_contrast+(1|Population)+(1|Habitat), data=subset(df1, NegativeSlope==0))
    summary(mod1)
    qqPlot(residuals(mod1))
    hist(residuals(mod1))  
    
    
    
    df1$NegPos = ifelse(df1$TG_Render_L_mean - df1$BG_E2.5_L_mean<0,"Negative","Positive")
    mod1 <- lmer(Fit~NegPos*trLGT+(1|Population)+(1|Habitat)+(1|LAB_Mean_Difference_BG_E2.5), data=subset(df1, NegativeSlope==0))
    summary(mod1)
    qqPlot(residuals(mod1))
    hist(residuals(mod1))  


#---___________________________________________________________________----
#----..............4. Match, Fitness and Complexity----
#----___________________________________________________________________

# UNUSED, UNUSED, UNUSED,UNUSED,UNUSED
#......................................
    
      p1 <- ggplot(subset(df1,OutlierFail==0), aes(x=LAB_Mean_Difference_BG_E2.5, y=Fitness,col=Cplx_Split2,fill=Cplx_Split2)) +
      geom_smooth (alpha=0.25,size=0.75)
    p1 + theme_classic()  +
      scale_linetype_manual(values=c( "solid","dotted"))+
      labs(title="",x ="LAB Difference", y = "Fitness", col="Complexity", fill="Complexity")+
      theme(text = element_text(size = 14), line = element_line(size=0.4))  + 
      theme(
        axis.text = element_text(family = "sans", size = 12),
        axis.line = element_line(colour = "black", size=0),
        panel.border = element_rect(colour = "black", fill=NA, size=0.4))
    
# The more complex the background the more linear the slope
```
