## Supplementary material for "Shining a light on camouflage evolution: using genetic algorithms to determine the effects of geometry and lighting on optimal camouflage": README_file.txt

This README.txt file was generated on 2025.02.24 [yyy,mm,dd] by George RA Hancock

1) GENERAL INFORMATION

---------------------------------------------------------------------------------------------------------------------------------------------------------

1. Title of Dataset: Shining a light on camouflage evolution: using genetic algorithms to determine the effects of geometry and lighting on optimal camouflage

2. Author Information
	A. Principal Investigator Contact Information
		Name: George Hancock
		Institution: University of Exeter
	

3. Date of data collection. June 2020 - June 2024 

4. Data collection location: University of Exeter, Penryn Campus. 

5. Funding Body: Natural Environment Research Council (NERC) GW4+ PhD and the Game and Wildlife Conservation Trust (GWCT).

6. Software recommendations. 

	(All software listed is freely available)

	- R version 4.3.2+ [https://www.r-project.org/]
	
	- ImageJ 1.5.2 [https://imagej.net/ij/]

	- MICA image analysis toolbox v2.1.1 [http://www.empiricalimaging.com/download/micatoolbox/]

	- Supplementary ImageJ Plugins.zip [Included in repository]


2) DATA & FILE OVERVIEW

---------------------------------------------------------------------------------------------------------------------------------------------------------


1. File List: 

   o. /Datasets/
	(In Alphabetical order)	

	# Base_Data.txt
	
	- Contains most of the base information for each target including:
		The background it was shown against.
		The length of the cast shadow.
		All of its gene values.
		The GabRat of the target with and without the background.
		The luminance gradient of the target when it is against the averaged natural (direct), diffuse and mixed lighting rey sphere.

	# BG_Geometry.txt
	
	- Contains the mean and standard deviation of Z (depth) for an area x15 the diameter of the target (30mm).
	- Additionally the dataset includes information necessary to merge with other datasets and the number for the original scan.
	- In instances where the scan failed the values are replaced by the average for the habitat it belonged to.


	# BG_Info.txt
	
	- Contains useful information for each backgorund photograph, most of which was unused within the final manuscript.
		 Information about the shadows (are cast shadows visible, is the target entirely in shadow).
		 Information about the backgorund, what is the dominant substrate, are certain features present e.g. stones, is the habitat
			anthropogenic?

	# BG_SiteInfo.txt
	
	- Contains useful information for the photography session of each site.
		 Year,month,day the photographs were taken.
		 What was the location UK county, lat, long.
		 What was the sun elevation and sun azimuth for that time of year.	
	- Some of this data is included in Site Info Table in the supplementary material.


	# LightCamo_BG_2.5.txt
	
	- Contains the gabor output metrics for the background or an area x2.5 (local) the diameter of the target (30mm).

	# LightCamo_BG_15.txt
	
	- Contains the gabor output metrics for the background or an area x15 (global) the diameter of the target (30mm).


	# LightCamo_EdgeEnhance.txt
	
	- Contains the gabor output metrics for the target skin with the max edge enhancement and without the edge enhancement.


	# LightCamo_Render.txt
	
	- Contains the gabor output metrics for the target pattern when it is rendered on its background.

	# LightCamo_Skin.txt
	
	- Contains the gabor output metrics for the target pattern when it isn't rendered on its background.


   o. /Sample_Evolved_Patterns/

	- Each habitat folder contains a subfolder for each evolution condition "1_Natural" = direct lighting only, "2_Diffuse" = diffuse only, 
	  and "3_Mixed" = 50:50 of direct and diffuse.

	- Within each folder is the fittest target from the final generation, G=20.


   o. /Supplementary_ImageJ_Plugins/
	
	# /3D_Light_CamoEvo_Scripts/

	- Contains the batch gabor measurement scripts. These can be run on the sample calibrated photos as a demo.
		To do so run the script and select the /Calibrated_Photos/ folder when prompted to.
	
	- Step 1, run the Batch_ scripts by running them in ImageJ and selecting the folder /Sample_Scene_Data/

	- Step 2, run the _Combine_ scripts by running them in ImageJ and selecting the folder /Sample_Scene_Data/

	NB1. requires sample scene data from full download link.

	NB2. for a more accessible version of these scripts that operate for the micaToolbox please see:

			https://github.com/GeorgeHancock471/3D_RNL_Tools


	# /3D_Measures/

	- Import_Normalised_PLY.txt, allows the user to select a ply that has been standardised in meshlab into ImageJ.

	- PLY Labeller, allows the user to select a ply and add the rois and adjust the height values to be relative to the selected target.

	NB1. requires sample scene data from full download link.


	# /ButtonGame/

	- Includes scripts used to run the online evolution game including the setup script which sets directories and the 
	  gated scripts for obtraining fitness data, generating the targets and sending them to the server, downloading the
	  server data and running the genetic algorithm.

	# ButtonGame CamoGen

	- Includes scripts used to create the CamoEvo patterns with spherical deformation.


	# /Spatial_Analyses/

	- Includes Jolyon Troscianko's Difference of Gaussians (DoG) and Gabor filter spatial analysis scripts.


   o. RCode_CamoEvo_Online.R
     
        Contains the necessary R code for recreating the plots and statistical analyses used in the paper.


2. Relationship between files, if important: 


   o. /Sample_Scene_Data/Adjusted Scans/
       	To run the R code, it must be in a folder that contains /Datasets/ with all of the stored .txt files
	This is because the R code automatically detects the directory of the R code and sets it as the working directory.


   o. /Supplementary_ImageJ_Plugins/
       	To install copy and paste the content of the plugins folder into the plugins folder of ImageJ v1.52 with MICA installed.


   o. RCode_CamoEvo_Online and Datasets

       	To run the R code, it must be in a folder that contains /Datasets/ with all of the stored .txt files
	This is because the R code automatically detects the directory of the R code and sets it as the working directory.


3. Additional related data collected that was not included in the current data package: N/A

4. Are there multiple versions of the dataset? NO
	A. If yes, name of file(s) that was updated: N/A
		i. Why was the file updated? N/A
		ii. When was the file updated? N/A


3) METHODOLOGICAL INFORMATION

---------------------------------------------------------------------------------------------------------------------------------------------------------

1. Description of methods used for collection/generation of data: 

These methods are a duplicate of: https://doi.org/10.1101/2025.02.19.639053

   o. Photography

	Photographs of 28 different habitats were taken from a ~1.2m using an ASUS A002 smartphone calibrated with imageJ using a 8% reflectance standard. 
	Direct lighting photos were taken on sunny days with <10% cloud cover and the sun unobstructed by passing cloud. Diffuse lighting photos were taken
	from the same location and on the same day using a Neewer 1.5m^3 diffuser tent placed over the top. 

        Uncalibrated copies of the photos are available on request.


   o. 3D Scans

	Scans were taken using the Matterport Scenes app for Android IOS with Tango enabled google depth sensors (a feature of the ASUS A002 smartphone). Each
	scan was taken from the same location as the scene photograph. As the scans worked better under diffuse lighting, but the app could not be triggered via
	bluetooth the photography tent was folded and used as a single diffuser screen when taking the scans. 3D scans were standardised in MeshLab v.2022.02 and 
	exported as .ply files containing only the X,Y, and Z coordinates. .ply files were loaded in ImageJ with a custom script to render an X,Y, Z image where 
	1px equated to 1mm. Blank z values were replaced locally using nearest-neighbour filling, and the grey-targets were labelled using ROIs. 


   o. Analysis Measures.
     
      	For each scene the the photographs and height maps were rescaled to the minimum number of pixels per mm. The photographs were then converted into the
	human CIE L*a*b* colour space. For the photographs and the scans the mean and standard deviation were measured. Then for the pattern scale and orientation
	analyses gabor filters were used to extract the pattern contrast at 8 different spatial scales [0.878mm, 1.758mm, 3.516mm , 7.031mm, 14.062mm, 28.125mm, 
	56.25mm, 112.5mm] and 4 different angles [0°,45°,90°,135°]. For each scale the max energy at a given angle and the energy at the orthogonal angle were
	also reported. 


2. Methods for processing the data: 

      Reconstructing the data used requires the calibrated background photographs and the supplementary ImageJ plugins


3. Instrument- or software-specific information needed to interpret the data: 

	- R version 4.3.2+ [https://www.r-project.org/]


4. Standards and calibration information, if appropriate: 

	- 8% reflectance standard

5. Environmental/experimental conditions: 

	- artificially diffuse lighting conditions, dry with <10% cloud cover


4) DATA-SPECIFIC INFORMATION FOR: 
	(In Alphabetical order)	


4.1) Base_Data.txt
---------------------------------------------------------------------------------------------------------------------------------------------------------
Background Geometry Data

See handbook abbreviation, CamoEvo = CE      ImageGA = IG
CamoEvo Handbook: 
ImageGA Handbook:

1. Number of variables: 100

2. Number of cases/rows: 42336

3. Variable List: 

	[1] Habitat. the unique habitat ID e.g. 01_grassWildFlowers or 22_estuaryHabitat.
	[2] Background.ID. the photograph code (N=natural, D=diffuse), number equals scene number (0-23) 
					e.g. N_background_0.tif = the natural (direct lighting photo) for the first scene.
	[3] Unique-Background. the unique code for each photograph = Habitat + PhotoName.
	[4] Unique-Location. the unique code for each scene = Habitat + Scene Number (starts from zero, 0-23).
	[5] Lighting. what lighting condition was the target evolved under [1_Natural = direct only, 2_Diffuse = diffuse only, and 3_Mixed = 50:50 direct and diffuse)
	[6] Unique-ID.  the unique ID for the habitat, lighting and target.
				e.g. 01_grassWildFlowers.1_Natural.Gen0_Mut0_ID10.N_background_14
					Habitat = 01_grassWildFlowers
					Lighting = 1_Natural
					TargetID = Gen0_Mut0_ID10

	[7] Unique-Render-ID. the unique ID for the habitat, lighting, target and background.
				e.g. 01_grassWildFlowers.1_Natural.Gen0_Mut0_ID10.N_background_14
					Habitat = 01_grassWildFlowers
					Lighting = 1_Natural
					TargetID = Gen0_Mut0_ID10
					Background = N_background_14 (direct lighting, background 14).

	[8] Shadow_Distance. the length of the cast shadow in pixels.
	[9] Background_ID. Duplicate of Background-ID, carry over from old data
	[10] Index. row number within the dataset
	[11] HabitatType. the broad habitat type of the habitat.
	[12] Habitat. the unique habitat ID e.g. 01_grassWildFlowers or 22_estuaryHabitat.
	[13] Lighting. what lighting condition was the target evolved under [1_Natural = direct only, 2_Diffuse = diffuse only, and 3_Mixed = 50:50 direct and diffuse)
	[14] Generation. which generation 0-20 the target is from.
	[15] Thirds. whehther the generation is in the starting, middle or final third (1,2,3);
	[16] S-M-F. whehther the generation is in the starting, middle or final third (s,m,f)
	[17 ID. the ID code for the target.
	[18] Fitness. the capture time in milliseconds for the target
	[19] Parent1. The parent ID of the individual, generation 0 have no parents.
	[20] Parent2. The parent ID of the individual, generation 0 have no parents.
	[21] Sex. Indicates the sex of the individual, though targets for this were sexless (-).
      	[22-55] Genes. The genes that regulate the target phenotype. See CE & IG. 
      	[56] gls_lvl_bkg. (Unique Gene) The gloss level of the target background (unique gene for online game, see supplementary material A)
      	[57] gls_lvl_mac. (Unique Gene) The gloss level of the target maculation (unique gene for online game, see supplementary material A)
	[58] Generation. which generation 0-20 the target is from.
	[59] Target-ID.  repeat of ID variable, some instances used Target-ID as a legacy variable.
	[60] Background-ID.  repeat of Background_ID variable, some instances used Background-ID as a legacy variable.
	[61-66] GabRat Variables. Indicate the GabRat metric for edge disruption, variables are encoded as follows
				  e.g. Render_L_GabRat
				  Render = against background, Independent = against average background
				  L = CIE L* channel, A = CIE a* channel, B = CIE b* channel
				  GabRat indicates GabRat
	[67-78] Gloss Variables. Measures of glossiness
				e.g. Gloss_Whole_Mean
				Gloss = change in value when gloss is applied, GlossMx = change in value when max gloss is applied
				Whole = whole target, Back = target background, Mac = target maculation
				Mean = mean luminance L*, Dev = standard deviation of luminance
	[79-90] Sphere Gradients. Measures of channel gradients used for countershading analyses
				e.g. Natural_Sphere_L_Gradient
				Natural_Sphere = average grey sphere for direct lighting, Diffuse_Sphere = average grey sphere for direct lighting, 
				... Mixed_Sphere = average grey sphere for direct lighting, PatternSkin = the un-rendered target
  				L = CIE L* channel, A = CIE a* channel, B = CIE b* channel
	[90-100] Sphere Gradients. Measures of channel gradients used for countershading analyses for the rendered targets
				e.g. Render_L_Gradient_Natural
  				L = CIE L* channel, A = CIE a* channel, B = CIE b* channel
				_Natural = target rendered on average direct, _Diffuse = target rendered on average diffuse, _Diffuse = target rendered on average mixed


4.2) BG_Geometry.txt
---------------------------------------------------------------------------------------------------------------------------------------------------------
Background Geometry Data

1. Number of variables: 6

2. Number of cases/rows: 672

3. Variable List: 

	[1] Habitat. the unique habitat ID e.g. 01_grassWildFlowers or 22_estuaryHabitat
	[2] Unique-Location. the unique code for each scene = Habitat + Scene Number (starts from zero, 0-23)
	[3] Scan-Original. The code number of the original scan (starts from 1, 1-24)
	[4] Unique-Scan. The unique code of the original scan  = Habitat + Scan-Original
	[5] BG_E15_Z_Mean. The Mean Z(height) for an area 15x the diameter of the target.
	[6] BG_E15_Z_Dev. The StdDev of Z(height) for an area 15x the diameter of the target.


4.3) BG_Info.txt
---------------------------------------------------------------------------------------------------------------------------------------------------------
Scene Information

1. Number of variables: 16

2. Number of cases/rows: 1344 (NB double geometry data length due to 2x lighting)

3. Variable List: 

	[1] Habitat. the unique habitat ID e.g. 01_grassWildFlowers or 22_estuaryHabitat.
	[2] Background.ID. the photograph code (N=natural, D=diffuse), number equals scene number (0-23) 
					e.g. N_background_0.tif = the natural (direct lighting photo) for the first scene.
	[3] Unique.Background. the unique code for each photograph = Habitat + PhotoName.
	[4] Unique.Location. the unique code for each scene = Habitat + Scene Number (starts from zero, 0-23).
	[5] RecievedShadows. does the target have recieved shadows on it 0/1, no/yes (binary) .
	[6] SmotheredShadow. is the target entirely in shadow 0/1, no/yes (binary).
	[7] Dappled. is there dappled lighting within the scene 0/1, no/yes (binary).
	[8] CastShadow. is the cast shadow of the target visible 0/1, no/yes (binary).
	[9] ExposureIssue. is the image over exposed in dappled regions 0/1, no/yes (binary).
	[10] Anthropogenic. is the scene anthropogenic or does it include manmade objects 0/1, no/yes (binary).
	[11] DominantSubstrate. which substrate makes up the majority of the scene? grass,leaflitter,bare,veg,gravel?
	[12] stones. does the scene have stones 0/1, no/yes (binary).
	[13] leaves. does the scene have leaflitter 0/1, no/yes (binary).
	[14] veg. does the scene have non-grass vegetation 0/1, no/yes (binary).
	[15] grass. does the scene have grass 0/1, no/yes (binary).
	[16] bare. does the scene have bare ground 0/1, no/yes (binary).


4.4) Site_Info.txt
---------------------------------------------------------------------------------------------------------------------------------------------------------
Site Information

1. Number of variables: 12

2. Number of cases/rows: 28 (one for each habitat)

3. Variable List: 

	[1] Habitat. the unique habitat ID e.g. 01_grassWildFlowers or 22_estuaryHabitat.
	[2] Year. the year the photographs were collected.
	[3] Month. the month the photographs were collected.
	[4] Day. the day the photographs were collected.
	[5] Time. the time of day (UK time) that the collection started (collection took ~2 hours)
	[6] TimeUTC. the UTC time.
	[7] Country. the country that the photographs were taken (always UK).
	[8] County. the county the photographs were taken.
	[9] Lat. the latitude of the photography site.
	[10] Long. the longitude of the photography site.
	[11] SunElevation. the elevation of the sun at the start of collection.
	[12] SunAzimuth. the azimuth of the sun at the start of collection.


4.5) LightCamo_BG_2.5.txt
---------------------------------------------------------------------------------------------------------------------------------------------------------
Background Energy Metrics for the local background

1. Number of variables: 152

2. Number of cases/rows: 1344 (two for each background)

3. Variable List: 

	[1] Habitat. the unique habitat ID e.g. 01_grassWildFlowers or 22_estuaryHabitat.
	[2] Background-ID. the photograph code (N=natural, D=diffuse), number equals scene number (0-23) 
					e.g. N_background_0.tif = the natural (direct lighting photo) for the first scene.
	[3] Unique.Background. the unique code for each photograph = Habitat + PhotoName.
	[4] Unique.Location. the unique code for each scene = Habitat + Scene Number (starts from zero, 0-23).
	[5-152] Due to the large number of repeats the variables are broken down by their codes. 
			Output here represents the energy output metrics for different scales, orientations and channels.
			
			LightCamo_2.5, denotes that the metric is for the local background (2.5x target scale).
			
			Next is whether it is luminance (L*, [5-68]) , red-green (A, [69-132]) or blue-yellow (B,[133-196])
			
			Next scale S smaller numbers indicate smaller spatial scales (higher frequency)
				[1,2,3,4,5,6] = [0.878mm, 1.758mm, 3.516mm , 7.031mm, 14.062mm, 28.125mm]
				
			Next is Gabor, this simply indicates that these are metrics from the gabor output.
			
			Then lastly is the specific metric these are:
			contrast, the average energy across all angles for that scale.
			deviation, the standard deviation of contrast for the angles at that scale.
			peak, the contrast at the angle with the highest contrast.
			orthogonal, the contrast at the angle orthogonal to the highest contrast.
			D[0,90,45,135], the contrast at the different angles these are used to calculate the directionality and the V (0) - H (90) measures in the R code.


4.6) LightCamo_BG_15.txt
---------------------------------------------------------------------------------------------------------------------------------------------------------
Background Energy Metrics for the global background

1. Number of variables: 152

2. Number of cases/rows: 1344 (two for each background)

3. Variable List: 

	[1] Habitat. the unique habitat ID e.g. 01_grassWildFlowers or 22_estuaryHabitat.
	[2] Background-ID. the photograph code (N=natural, D=diffuse), number equals scene number (0-23) 
					e.g. N_background_0.tif = the natural (direct lighting photo) for the first scene.
	[3] Unique.Background. the unique code for each photograph = Habitat + PhotoName.
	[4] Unique.Location. the unique code for each scene = Habitat + Scene Number (starts from zero, 0-23).
	[5-152] Due to the large number of repeats the variables are broken down by their codes. 
			Output here represents the energy output metrics for different scales, orientations and channels.
			
			LightCamo_15, denotes that the metric is for the global background (15x target scale).
			
			Next is whether it is luminance (L) , red-green (A),  or blue-yellow (B)
			
			Next scale S smaller numbers indicate smaller spatial scales (higher frequency)
				[1,2,3,4,5,6] = [0.878mm, 1.758mm, 3.516mm , 7.031mm, 14.062mm, 28.125mm]
				
			Next is Gabor, this simply indicates that these are metrics from the gabor output.
			
			Then lastly is the specific metric these are:
			contrast, the average energy across all angles for that scale.
			deviation, the standard deviation of contrast for the angles at that scale.
			peak, the contrast at the angle with the highest contrast.
			orthogonal, the contrast at the angle orthogonal to the highest contrast.
			D[0,90,45,135], the contrast at the different angles these are used to calculate the directionality and the V (0) - H (90) measures in the R code.

4.7) LightCamo_Render.txt
---------------------------------------------------------------------------------------------------------------------------------------------------------
Energy Metrics for the rendered targets

1. Number of variables: 156

2. Number of cases/rows: 42336 (one for each target)

3. Variable List: 

	[1] Habitat. the unique habitat ID e.g. 01_grassWildFlowers or 22_estuaryHabitat.
	[2] Lighting. what lighting condition was the target evolved under [1_Natural = direct only, 2_Diffuse = diffuse only, and 3_Mixed = 50:50 direct and diffuse)
	[3] Generation. which generation 0-20 the target is from.
	[4] Target-ID. the ID code for the target.
	[5] Background-ID. the photograph code (N=natural, D=diffuse), number equals scene number (0-23) 
					e.g. N_background_0.tif = the natural (direct lighting photo) for the first scene.
	[6] Unique-ID. the unique ID for the target habitat, lighting and target
				e.g. 01_grassWildFlowers.1_Natural.Gen0_Mut0_ID10.N_background_14
					Habitat = 01_grassWildFlowers
					Lighting = 1_Natural
					TargetID = Gen0_Mut0_ID10
	[7] Unique-Render-ID. the unique ID for the target habitat, lighting, target, and background
				e.g. 01_grassWildFlowers.1_Natural.Gen0_Mut0_ID10.N_background_14
					Habitat = 01_grassWildFlowers
					Lighting = 1_Natural
					TargetID = Gen0_Mut0_ID10
					Background = N_background_14 (direct lighting, background 14).
	[8-13]  Due to the large number of repeats the variables are broken down by their codes. 
			Output here represents the mean and standard deviation for the different channels.
			e.g. TG_Render_L_mean

			TG_Render, denotes that the metric is for the remdered target.
			
			Next is whether it is luminance (L) , red-green (A),  or blue-yellow (B)

			Lastly is the measure the mean or the stddv

	[14-156] Due to the large number of repeats the variables are broken down by their codes. 
			Output here represents the energy output metrics for different scales, orientations and channels.
			
			TG_Render, denotes that the metric is for the remdered target.
			
			Next is whether it is luminance (L) , red-green (A),  or blue-yellow (B)
			
			Next scale S smaller numbers indicate smaller spatial scales (higher frequency)
				[1,2,3,4,5,6] = [0.878mm, 1.758mm, 3.516mm , 7.031mm, 14.062mm, 28.125mm]
				
			Next is Gabor, this simply indicates that these are metrics from the gabor output.
			
			Then lastly is the specific metric these are:
			contrast, the average energy across all angles for that scale.
			deviation, the standard deviation of contrast for the angles at that scale.
			peak, the contrast at the angle with the highest contrast.
			orthogonal, the contrast at the angle orthogonal to the highest contrast.
			D[0,90,45,135], the contrast at the different angles these are used to calculate the directionality and the V (0) - H (90) measures in the R code.


4.8) LightCamo_Skin.txt
---------------------------------------------------------------------------------------------------------------------------------------------------------
Energy Metrics for the un-rendered target skins

1. Number of variables: 156

2. Number of cases/rows: 42336 (one for each target)

3. Variable List: 

	[1] Habitat. the unique habitat ID e.g. 01_grassWildFlowers or 22_estuaryHabitat.
	[2] Lighting. what lighting condition was the target evolved under [1_Natural = direct only, 2_Diffuse = diffuse only, and 3_Mixed = 50:50 direct and diffuse)
	[3] Generation. which generation 0-20 the target is from.
	[4] Target-ID. the ID code for the target.
	[5] Background-ID. the photograph code (N=natural, D=diffuse), number equals scene number (0-23) 
					e.g. N_background_0.tif = the natural (direct lighting photo) for the first scene.
	
	[6] Unique-ID. the unique ID for the target habitat, lighting and target
				e.g. 01_grassWildFlowers.1_Natural.Gen0_Mut0_ID10.N_background_14
					Habitat = 01_grassWildFlowers
					Lighting = 1_Natural
					TargetID = Gen0_Mut0_ID10
	[7] Unique-Render-ID. the unique ID for the target habitat, lighting, target, and background
				e.g. 01_grassWildFlowers.1_Natural.Gen0_Mut0_ID10.N_background_14
					Habitat = 01_grassWildFlowers
					Lighting = 1_Natural
					TargetID = Gen0_Mut0_ID10
					Background = N_background_14 (direct lighting, background 14).
	[8-13]  Due to the large number of repeats the variables are broken down by their codes. 
			Output here represents the mean and standard deviation for the different channels.
			e.g. TG_Skin_L_mean

			TG_Skin, denotes that the metric is for the remdered target.
			
			Next is whether it is luminance (L) , red-green (A),  or blue-yellow (B)

			Lastly is the measure the mean or the stddv

	[14-156] Due to the large number of repeats the variables are broken down by their codes. 
			Output here represents the energy output metrics for different scales, orientations and channels.
			
			TG_Render, denotes that the metric is for the remdered target.
			
			Next is whether it is luminance (L) , red-green (A),  or blue-yellow (B)
			
			Next scale S smaller numbers indicate smaller spatial scales (higher frequency)
				[1,2,3,4,5,6] = [0.878mm, 1.758mm, 3.516mm , 7.031mm, 14.062mm, 28.125mm]
				
			Next is Gabor, this simply indicates that these are metrics from the gabor output.
			
			Then lastly is the specific metric these are:
			contrast, the average energy across all angles for that scale.
			deviation, the standard deviation of contrast for the angles at that scale.
			peak, the contrast at the angle with the highest contrast.
			orthogonal, the contrast at the angle orthogonal to the highest contrast.
			D[0,90,45,135], the contrast at the different angles these are used to calculate the directionality and the V (0) - H (90) measures in the R code.
