## Supplementary material for "Shining a light on camouflage evolution: using genetic algorithms to determine the effects of geometry and lighting on optimal camouflage": CIELAB_32Bit_to_RGB24_smooth.java

```
/*__________________________________________________________________

	Title: CIE LAB Colour Space (32-bit) to RGB (24-bit)
	''''''''''''''''''''''''''''''''''''''''''''''''''''''''''''''''''''''''''''''''''''''''
	Author: Jolyon Troscianko
	Date: 17/1/2018
............................................................................................................

This code converts 32-bits/channel CIE LAB to 8 bits/channel RGB,
based on the procedure used here:

http://www.easyrgb.com/index.php?X=MATH&H=07#text7

The code requries a 32-bit LAB stack

____________________________________________________________________
*/


import ij.*;
import ij.plugin.filter.PlugInFilter;
import ij.process.*;

public class CIELAB_32Bit_to_RGB24_smooth implements PlugInFilter {

ImageStack stack;
	public int setup(String arg, ImagePlus imp) { 
	stack = imp.getStack(); 
	return DOES_32 + STACK_REQUIRED; 
	}
public void run(ImageProcessor ip) {

float[] L;
float[] A;
float[] B;

int w = stack.getWidth();
int h = stack.getHeight();
int dimension = w*h;

L = (float[]) stack.getPixels(1);
A = (float[]) stack.getPixels(2);
B = (float[]) stack.getPixels(3);

int[] outRGB = new int[dimension];

double var_R = 0.0;
double var_G = 0.0;
double var_B = 0.0;

double var_Y = 0.0;
double var_X = 0.0;
double var_Z = 0.0;

double RefX = 95.047; // 2 degree 1931, D65 values
double RefY = 100.000;
double RefZ = 108.883;


int sR = 0;
int sG =0;
int sB = 0;

for(int i=0; i<dimension; i++){

	var_Y = ( L[i] + 16 ) / 116;
	var_X = A[i] / 500 + var_Y;
	var_Z = var_Y - B[i] / 200;

	//if ( Math.pow(var_Y,3)  > 0.008856 )
		var_Y = Math.pow(var_Y,3);
	//else
	//	var_Y = ( var_Y - 16 / 116 ) / 7.787;

	//if ( Math.pow(var_X,3)  > 0.008856 )
		var_X = Math.pow(var_X,3);
	//else
	//	var_X = ( var_X - 16 / 116 ) / 7.787;

	//if ( Math.pow(var_Z,3)  > 0.008856 )
		var_Z =  Math.pow(var_Z,3);
	//else
	//	var_Z = ( var_Z - 16 / 116 ) / 7.787;

	var_X = (var_X * RefX)/100;
	var_Y = (var_Y * RefY)/100;
	var_Z = (var_Z * RefZ)/100;

	//----temp---------
	//X[i] = (float) var_X;
	//Y[i] = (float) var_Y;
	//Z[i] = (float) var_Z;

	var_R = var_X *  3.2406 + var_Y * -1.5372 + var_Z * -0.4986;
	var_G = var_X * -0.9689 + var_Y *  1.8758 + var_Z *  0.0415;
	var_B = var_X *  0.0557 + var_Y * -0.2040 + var_Z *  1.0570;

	//if ( var_R > 0.0031308 ) 
		var_R = 1.055 * ( Math.pow(var_R , ( 1 / 2.4 ) )) - 0.055;
	//else                     var_R = 12.92 * var_R;

	//if ( var_G > 0.0031308 )
		var_G = 1.055 * ( Math.pow(var_G , ( 1 / 2.4 )) ) - 0.055;
	//else                     var_G = 12.92 * var_G;

	//if ( var_B > 0.0031308 )
		var_B = 1.055 * ( Math.pow(var_B , ( 1 / 2.4 )) ) - 0.055;
	//else                     var_B = 12.92 * var_B;


	sR = (int) (Math.round(var_R * 255));
	sG = (int) (Math.round(var_G * 255));
	sB = (int) (Math.round(var_B * 255));

	if(sR >= 0 && sR <= 255)
		sR = sR;
	else if(sR >= 255)
		sR = 255;
	else sR = 0;


	if(sG >= 0 && sG <= 255)
		sG = sG;
	else if(sG >= 255)
		sG = 255;
	else sG = 0;


	if(sB >= 0 && sB <= 255)
		sB = sB;
	else if(sB >= 255)
		sB = 255;
	else sB = 0;


	outRGB[i] = ((sR << 16) | ((sG << 8) | sB));

}//i


ImageStack outStack = new ImageStack(w, h);
outStack.addSlice("RGB", outRGB);
new ImagePlus("Output", outStack).show();


}
}
```
