## Supplementary material for "Shining a light on camouflage evolution: using genetic algorithms to determine the effects of geometry and lighting on optimal camouflage": Reaction_Diffusion_Pattern_Generator.java

```
/*__________________________________________________________________

	Title: Gray-Scott Reaction Diffusion Model
	''''''''''''''''''''''''''''''''''''''''''''''''''''''''''''''''''''''''''''''''''''''''
	Author: Jolyon Troscianko
	Date: 8/1/2018
............................................................................................................

This code generates Reaction Diffusion patterns.

The algorithm joins opposite edges, meaning the patterns can be tiled, and the growth
is effectively on a torus-type surface.

The starting condition is: all "A" concentrations to 1, and all "B" concentrations to 0,
except for a randomly located vertical line 1 px thick, where "B" concentrations are 1.

The concentrations of A and B are limited between 0 and 1.

Random noise is added to A and B, which helps stabilise pattern growth with low values,
and at high values breaks up the patterns

The patterns grow from their random start column to their surroundings. Once the
patterns reach the opposite end of the image the growth continues for a length
of time dependant on the width of the image. Small images require additional extra
growth time for the patterns to start to stabilise. An exponential decay is used to 
calculate the number of additional steps y = 5000*0.996^w where w is image width px.

Useful descriptions:
http://www.algosome.com/articles/reaction-diffusion-gray-scott.html
http://www.mrob.com/pub/comp/xmorphia/

Useful example code:
https://codepen.io/eskimoblood/pen/sFKkc


____________________________________________________________________
*/

import ij.*;
import ij.plugin.PlugIn;
import ij.process.*;
import ij.gui.*;


public class Reaction_Diffusion_Pattern_Generator implements PlugIn {
public void run(String arg) {

IJ.showStatus("Reaction Diffusion");

int w = 100;
int h = 100;
int nSteps = 10000; // maximum number of steps
double randomFeed = 0.01;

double f =  0.025;
double k = 0.077;
double Da = 0.2;
double Db = Da/2;

/*

Keeping k around 0.77 works well.
f = 0.020 = fail
f = 0.0215 = spots (homogeneous, fails with small images)
f = 0.022 = spots (wobbly)
f = 0.023 = stripes (less connected)
f = 0.024 = stripes (quite connected)
f = 0.025 = inverse spots (only form when image is large enough, around 200)
f = 0.026 = chaotic markings
f = 0.027 = mottle (due to random values)
above with noise around 0.01, values below this don't seem to make a difference
highest noise should  be around 0.2, 0.05 is intermediate/high, 0.01 is intermediate, 0.001 is low

*/


GenericDialog gd = new GenericDialog("Reaction Diffusion Pattern Generation");
		gd.addMessage("Pattern Generation Values");
		gd.addNumericField("f Feed rate", f, 5);
		gd.addNumericField("k Kill rate", k, 5);
		gd.addNumericField("Da Diffusion rate", Da, 5);
		gd.addNumericField("Db Diffusion rate", Db, 5);
		gd.addNumericField("noise", randomFeed, 5);
		gd.addNumericField("Max steps", nSteps, 0);
		gd.addMessage("Image Dimensions");
		gd.addNumericField("w Width", w, 0);
		gd.addNumericField("h Height", h, 0);
gd.showDialog();
	if (gd.wasCanceled())
		return;

	
f = gd.getNextNumber();
k = gd.getNextNumber();
Da = gd.getNextNumber();
Db = gd.getNextNumber();
randomFeed = gd.getNextNumber();
nSteps = (int) Math.round( gd.getNextNumber());
w = (int) Math.round( gd.getNextNumber());
h = (int) Math.round( gd.getNextNumber());


//---------------------Set up environment------------------

int dimension = w*h;

float[] outA = new float[dimension];
float[] outB = new float[dimension];
float[] A = new float[dimension];
float[] B = new float[dimension];

// initialise image
for(int i=0; i<dimension; i++){
	A[i] = 1;
	B[i] = 0;
}

// draw a vertical line in a random column of the image as a starting point
int midRow = (int) (Math.round(Math.random()*(w-1)));
for(int y=0; y<h; y++)
	B[(y*w)+midRow] = 1;

if(midRow >= Math.round(w/2)) // prepare midRow for finding the opposite side of the image (i.e last pixels to be filled)
	midRow = midRow - Math.round(w/2);
else midRow = midRow + Math.round(w/2);

double Aval = 0.0;
double Bval = 0.0;

double convCount = 0.0; // used to work with image edges
double cA = 0.0; // convolution sum A
double cB = 0.0; // convolution sum A

double maxB = 0.0;
double minB = 10E10;
double xBSum = 0.0;
int rowsFilled = 0;
int finalFlag = 0;

int pxid = 0;
double d2 = 0.0;
double randVal = 0.0;


for(int j=0; j<nSteps; j++){


for (int y=0; y<h; y++)
for (int x=0; x<w; x++){

	//Laplacian Convolution

	cA = 0.0;
	cB = 0.0;
	pxid = (y*w)+x;
	convCount = 0.0;

	//TOP
	if(y>0){
		cA += A[pxid - w];
		cB += B[pxid - w];
	} else{
		cA += A[pxid + (w*h) - w];
		cB += B[pxid + (w*h) - w];
	}

	//LEFT
	if(x>0){
		cA += A[pxid - 1];
		cB += B[pxid - 1];
	} else{
		cA += A[pxid + w - 1];
		cB += B[pxid + w - 1];
	}


	//RIGHT
	if(x<w-1){
		cA += A[pxid + 1];
		cB += B[pxid + 1];
	} else{
		cA += A[pxid - w + 1];
		cB += B[pxid - w + 1];
	}

	//BOTTOM
	if(y<h-1){
		cA += A[pxid + w];
		cB += B[pxid + w];
	} else{
		cA += A[pxid - (w*h) + w];
		cB += B[pxid - (w*h) + w];
	}


	// CENTRE
	Aval = (double) A[pxid];
	Bval = (double) B[pxid];
	
	d2 = Aval*Bval*Bval;
	
	outA[pxid] = (float) (           Aval + ((Da* ( cA - 4 * Aval) - d2) + f * (1 - Aval))                );
	outB[pxid] = (float) (           Bval + ((Db* ( cB - 4 * Bval) + d2) - k*Bval)              );
	
	
	
	
}

xBSum = 0.0;

for (int y=0; y<h; y++){
for (int x=0; x<w; x++){

	pxid = (y*w)+x;

	if(outA[pxid] > 1)
		A[pxid] = 1;
	else if(outA[pxid] < 0)
		A[pxid] = 0;	
	else A[pxid] = outA[pxid];

	if(outB[pxid] > 1)
		B[pxid] = 1;
	else if(outB[pxid] < 0)
		B[pxid] = 0;	
	else B[pxid] = outB[pxid];

	A[pxid] += (float) (randomFeed*(Math.random()-0.5));
	if(j>nSteps-1) // add noise to B channel except fot the last go
		B[pxid] += (float) (randomFeed*(Math.random()-0.5));

	xBSum += B[pxid];

}//x
}//y

if(xBSum/dimension < 0.01/dimension) // stop when the pattern growth has failed
	j = nSteps;

//--------------is image filled yet?---------------
if(finalFlag == 0){
	xBSum = 0.0;
	for (int y=0; y<h; y++)
		xBSum += B[(y*w)+midRow];

	xBSum = xBSum/h;

	if(xBSum > 0.1){ // final row filled
		// use equation for exponential decay to increase the numebr of additional passes
		// with smaller images requiring additional extra passes.
		nSteps = (int) (j + Math.round(    Math.pow(0.996,w)*5000));

		finalFlag = 1;	
	}

}// final flag

IJ.showProgress((float) j/nSteps);

}//j steps

IJ.showProgress((float) 1);


ImageStack outStack = new ImageStack(w, h);
outStack.addSlice("B", B);
new ImagePlus("f"+ f + " rand" + randomFeed, outStack).show();


} // void
} //class
```
