## Supplementary material for "Shining a light on camouflage evolution: using genetic algorithms to determine the effects of geometry and lighting on optimal camouflage": Simple Normalisation.txt

selectWindow("background");


greyVals = newArray(nSlices); // the array where we'll store the grey values, one for each image channel
reflectance = newArray(27, 30, 33); // the grey standard reflectances


// Measure the grey standard (selected using an ROI - region of interest) in all image channels (slices)
row = nResults();
for(i=0; i<nSlices; i++){
	setSlice(i+1);
	label = getInfo("slice.label");
	getStatistics(area, mean, min, max, std); // gets the statistics from the current ROI
	setResult(label, row, mean);
	greyVals[i]=mean; // saves the mean value measured, and stores it in the "greyVals" array's slot i.
}

// Any transforms we apply will only happen to the selected areas, so de-select the grey standard:
run("Select None");

// run through all the slices and apply the relevant normalisation transform:
for(i=0; i<nSlices; i++){
	setSlice(i+1);
	scalingFactor = (reflectance[i]/100)*255/greyVals[i]; // use the known reflectance to work out what transform is required
	run("Multiply...", "value=&scalingFactor slice"); // apply the scaling factor to this channel only
	


	// Normally anything between quotation marks is a string of text only, the ampersand symbol tells it to use the variable we've just created instead
}

if (isOpen("glossMat")){

selectWindow("glossMat");

run("Select None");

for(i=0; i<nSlices; i++){
	setSlice(i+1);
	scalingFactor = (reflectance[i]/100)*255/greyVals[i]; // use the known reflectance to work out what transform is required
	run("Multiply...", "value=&scalingFactor slice"); // apply the scaling factor to this channel only
	// Normally anything between quotation marks is a string of text only, the ampersand symbol tells it to use the variable we've just created instead
}
	
}


if (isOpen("glossBkg")){

selectWindow("glossBkg");

run("Select None");

for(i=0; i<nSlices; i++){
	setSlice(i+1);
	scalingFactor = (reflectance[i]/100)*255/greyVals[i]; // use the known reflectance to work out what transform is required
	run("Multiply...", "value=&scalingFactor slice"); // apply the scaling factor to this channel only
	// Normally anything between quotation marks is a string of text only, the ampersand symbol tells it to use the variable we've just created instead
}
	
}


if (isOpen("glossMat")){

selectWindow("glossMat");

for(i=0; i<nSlices; i++){
	setSlice(i+1);
	scalingFactor = (reflectance[i]/100)*255/greyVals[i]; // use the known reflectance to work out what transform is required
	run("Multiply...", "value=&scalingFactor slice"); // apply the scaling factor to this channel only
	// Normally anything between quotation marks is a string of text only, the ampersand symbol tells it to use the variable we've just created instead
}
	
}


if (isOpen("glossBkg")){

selectWindow("glossBkg");

for(i=0; i<nSlices; i++){
	setSlice(i+1);
	scalingFactor = (reflectance[i]/100)*255/greyVals[i]; // use the known reflectance to work out what transform is required
	run("Multiply...", "value=&scalingFactor slice"); // apply the scaling factor to this channel only
	// Normally anything between quotation marks is a string of text only, the ampersand symbol tells it to use the variable we've just created instead
}
	
}
