## Supplementary material for "Shining a light on camouflage evolution: using genetic algorithms to determine the effects of geometry and lighting on optimal camouflage": Spherical_Deform.java

```
/*__________________________________________________________________

	Title: Spherical Image Deform
	''''''''''''''''''''''''''''''''''''''''''''''''''''''''''''''''''''''''''''''''''''''''
	Author: Jolyon Troscianko
	Date: 16/1/2018
............................................................................................................

This script takes a square image and 'wraps' it around a spherical
object, as if it were a 3D ball.

The script applies linear interpolation for minimising artefacts.

To create a 3D-simulated shadow, use a circular selection centered
on the 'top' point of the sphere.

____________________________________________________________________
*/


import ij.*;
import ij.plugin.filter.PlugInFilter;
import ij.process.*;


public class Spherical_Deform implements PlugInFilter {

ImageStack stack;
	public int setup(String arg, ImagePlus imp) { 
	stack = imp.getStack(); 
	return DOES_RGB; 
	}

public void run(ImageProcessor ip) {


int w = stack.getWidth();
int h = stack.getHeight();

int[] px =(int[]) stack.getPixels(1);

int[] outRGB = new int[w*h];
int R = 0;
int G = 0;
int B = 0;

int r = (int) (Math.round(w/2));

double angle = 0.0;
double distance = 0.0;
double hyp = 0.0;

double xxd = 0.0;
double yyd = 0.0;

int xx = 0;
int yy = 0;
int xxp = 0;
int yyp = 0;


double xaR = 0.0;
double xaG = 0.0;
double xaB = 0.0;
double xbR = 0.0;
double xbG = 0.0;
double xbB = 0.0;

double yaR = 0.0;
double yaG = 0.0;
double yaB = 0.0;
double ybR = 0.0;
double ybG = 0.0;
double ybB = 0.0;

	
for(int y=1; y<h-1; y++)
for(int x=1; x<w-1; x++){

	//if(x-r == 0)
	//	angle = 0.0;
	//else angle = Math.atan2( (y-r) / (x-r) ); // angle from centre
	angle = Math.atan2( (y-r) , (x-r) ); // angle from centre
	distance = Math.pow( Math.pow(x-r,2) + Math.pow(y-r,2) ,0.5); // distance from centre
	hyp = (2*r*(Math.asin(distance/r))) / Math.PI;

	xxd = Math.cos(angle)*hyp;
	yyd = Math.sin(angle)*hyp;

	xx = (int) (Math.round(xxd));
	yy = (int) (Math.round(yyd));

	xxd = xxd-xx;
	yyd = yyd - yy;

	if(xxd > 0)
		xxp = xx+1;
	else xxp = xx-1;

	if(yyd > 0)
		yyp = yy+1;
	else yyp = yy-1;

	xx += r; yy += r; xxp += r; yyp += r;

/*
	if(y >= r)
	if(x >= r){xx += r; yy += r; xxp += r; yyp += r;}

	if(y < r)
	if(x >= r){ xx += r; yy += r; xxp += r; yyp += r;}
		
	if(y < r)
	if(x < r){ xx = r-xx; yy = r-yy; xxp = r-xxp; yyp = r-yyp;}
		
	if(y >= r)
	if(x < r){ xx = r-xx; yy = r-yy; xxp = r-xxp; yyp = r-yyp;}
*/

	
	if(    (yy*w)+xx >=0    &&    (yy*w)+xx < w*h){

		xxd = Math.abs(xxd);
		yyd = Math.abs(yyd);
		
		xaR = ((px[(yy*w)+xx]>>16)&0xff) * (1-xxd);
		xbR = ((px[(yy*w)+xxp]>>16)&0xff) * xxd; // left/right

		xaG = ((px[(yy*w)+xx]>>8)&0xff) * (1-xxd);
		xbG = ((px[(yy*w)+xxp]>>8)&0xff) * xxd; // left/right

		xaB = (px[(yy*w)+xx]&0xff) * (1-xxd);
		xbB = (px[(yy*w)+xxp]&0xff) * xxd; // left/right

		yaR = ((px[(yy*w)+xx]>>16)&0xff) * (1-yyd);
		ybR = ((px[(yyp*w)+xx]>>16)&0xff) * yyd; // above/below

		yaG = ((px[(yy*w)+xx]>>8)&0xff) * (1-yyd);
		ybG = ((px[(yyp*w)+xx]>>8)&0xff) * yyd; // above/below

		yaB = (px[(yy*w)+xx]&0xff) * (1-yyd);
		ybB = (px[(yyp*w)+xx]&0xff) * yyd; // above/below
		
		R = (int) (Math.round(     (xaR+xbR+yaR+ybR)/2  ));
		G = (int) (Math.round(     (xaG+xbG+yaG+ybG)/2  ));
		B = (int) (Math.round(     (xaB+xbB+yaB+ybB)/2  ));

		outRGB[(y*w)+x] = ((R << 16) | ((G << 8) | B));
	}
} //xy


ImageStack outStack = new ImageStack(w, h);
outStack.addSlice("RGB", outRGB);
new ImagePlus("Output", outStack).show();


}
}
```
