## Supplementary material for "Shining a light on camouflage evolution: using genetic algorithms to determine the effects of geometry and lighting on optimal camouflage": Background_Image_Normalisation.txt

powerVal = 0.34; // specify the non-linear power function. e.g. 0.5 is square root, 0.4 is more extreme, 1 leaves the contrast unchanged


imDIR = getDirectory("Choose a folder containing images");


fileList = getFileList(imDIR);

imList = newArray(0);

for(i=0; i<fileList.length; i++)
	if(endsWith(fileList[i], ".tif") == 1 && startsWith(fileList[i], "Low") != 1|| endsWith(fileList[i], ".TIF") && startsWith(fileList[i], "Low") != 1)
		imList = Array.concat(imList, fileList[i]);

setBatchMode(false);


for(j=0; j<imList.length; j++){
//setBatchMode(true);


ts = imDIR + imList[j];
open(ts);


run("To ROI Manager");


run("RGB Stack");
run("32-bit");


setSlice(2);
for(i=1; i<=nSlices; i++){
	//run("Linearisation Function", "equation=[JT Linearisation] a=9.883462554 b=45.384996178 c=-0.000242783 d=0.071096231 slice=&i low=10 high=254");
	run("Linearisation Function", "equation=[JT Linearisation] a=9.883462554 b=45.384996178 c=100 d=0.071096231 slice=&i low=10 high=254");
	
	setSlice(1); // seems to not apply/update to currently active slice!
}


nRoi = roiManager("count");
roiNames = newArray(nRoi);

sphereCount = 0;
sphereLoc = 0;
vegCount = 0;
vegLoc = 0;


for(i=0; i<nRoi; i++){	
	roiManager("select", i);
	roiNames[i] = getInfo("selection.name");
	if(endsWith(roiNames[i], "sphere") == true){
		sphereCount++;
		sphereLoc = i;
	}
	if(endsWith(roiNames[i], "vegetation") == true){
		vegCount++;
		vegLoc = i;
	}
}

if(sphereCount > 1)
	exit("Image has more than one sphere!");

if(sphereCount == 0)
	exit("No sphere ROI found");

if(vegCount > 1)
	exit("Image has more than one sphere!");


if(vegCount == 1){
	roiManager("select", newArray(sphereLoc, vegLoc));
	roiManager("AND");
	roiManager("Add");
	roiManager("select", newArray(sphereLoc, nRoi));
	roiManager("XOR");
	roiManager("Add");
	roiManager("select", nRoi+1);
} else {
	roiManager("select", sphereLoc);
}


greyVals = newArray(3); // the array where we'll store the grey values, one for each image channel
reflectance = newArray(	4, 5, 7);  // the grey standard reflectances


for(i=0; i<nSlices; i++){
	setSlice(i+1);
	getStatistics(area, mean, min, max, std); // gets the statistics from the current ROI
	greyVals[i]=mean; // saves the mean value measured, and stores it in the "greyVals" array's slot i.
}

run("Select None");

if(vegCount == 1){ // clean up
	roiManager("select", newArray(nRoi, nRoi+1));
	roiManager("delete");
}

run("Select None");

// run through all the slices and apply the relevant normalisation transform:
setSlice(2);
for(i=0; i<nSlices; i++){
	setSlice(i+1);
	scalingFactor = (reflectance[i]/100)*255/greyVals[i]; // use the known reflectance to work out what transform is required
	run("Multiply...", "value=&scalingFactor slice"); // apply the scaling factor to this channel only
	// Normally anything between quotation marks is a string of text only, the ampersand symbol tells it to use the variable we've just created instead
}


// reduce contrast:

tm = 100 / pow(100, powerVal);
//tm = 1/powerVal;

setSlice(2);
run("Linearisation Function", "equation=Power a=&tm b=&powerVal c=0.000000000 d=0.000000000 slice=1");
setSlice(1);
run("Linearisation Function", "equation=Power a=&tm b=&powerVal c=0.000000000 d=0.000000000 slice=2");
run("Linearisation Function", "equation=Power a=&tm b=&powerVal c=0.000000000 d=0.000000000 slice=3");


//ts = "code=[v=pow(v, " + powerVal + ")] stack";
//run("Macro...", ts);

//run("Macro...", "code=[v=pow(v, 0.5)] stack");


run("Select All");

maxs = newArray(3);
mins = newArray(3);
for(i=0; i<nSlices; i++){
	setSlice(i+1);
	getStatistics(area, mean, min, max, std); 
	maxs[i]=max;
	mins[i]=min;
}

Array.getStatistics(maxs, minmax, maxmax, mean, std);
Array.getStatistics(mins, minmin, maxmin, mean, std);

middleMax = Array.copy(maxs);
middleMax = Array.sort(middleMax);
middleMax = middleMax[1];

middleMin = Array.copy(mins);
middleMin = Array.sort(middleMin);
middleMin = middleMin[1];

run("Make Composite", "display=Composite");
/*
Array.show("Values",mins,maxs);

print("minmax = " + minmax);
print("maxmax = " + maxmax);

print("minmin = " + minmin);
print("maxmin = " + maxmin);
*/

for(i=0; i<nSlices; i++){
	setSlice(i+1);
	setMinAndMax(maxmin, minmax); // setting the maximum to the lowest of all three channel's maximum values, and the minimum to the highest of the three channel's minima
}


run("RGB Color");

roiManager("Deselect");
run("From ROI Manager");

while(roiManager("count") > 0){
	roiManager("select", 0);
	roiManager("delete");
}


/*
rename("Shadow = " + powerVal);
close("0.tif");
*/

/*
setBatchMode("show");

ts =  imDIR + "LowishContrast_" + imList[j];


saveAs("Tiff", ts);

run("Close All");

}//j
