## Supplementary material for "Shining a light on camouflage evolution: using genetic algorithms to determine the effects of geometry and lighting on optimal camouflage": ButtonGame_README.txt

NB, because of when the game was made the majority of the code here can not be re-integrated into CamoEvo. 

If you wish to run a similar online game using CamoEvo, please contact George Hancock,
