## Supplementary material for "Shining a light on camouflage evolution: using genetic algorithms to determine the effects of geometry and lighting on optimal camouflage": DoG_ROI_bandpass_smooth.java

```
import ij.*;
import ij.process.*;
import ij.gui.*;
import java.awt.*;
import ij.plugin.*;

import ij.measure.ResultsTable;

/*

	Title: DoG ROI Bandpass
	Author: Jolyon Troscianko
	Date: 17/9/2021

.................................................................................................................

Description:
'''''''''''''''''''''

This script applies a bank of DoG filters to an ROI. The script has been designed to
work with ROIs of any (non-rectangular) shape by applying a 2D convolution and
weighting all measurements by the degree to which the kernel is inside the ROI
i.e. it re-normalises the kernel with each convolution.

The script efficiently applies the DoG filter to a series of different spatial scales
by effectively scaling the size of the image (with no interpolation). This makes
it incredibly fast at measuring the larger spatial frequencies that would otherwise
be incredibly slow to process (e.g. by using a larger sigma value).

The script then applies a bicubic interpolation (Catmull-Rom) to the output. This
interpretation assumes that pixels outside the ROI have a value of zero, which
makes sense given the convolution is normalised to have average output of zero.
This is different from a standard image, where out-of-bounds pixels cannot be assumed
to have a particular value.

The result is a very fast convolution, with increasing processing efficiency at larger
spatial scales, combined with smooth output. Any systematic errors introduced by
the scaling and interpolation will be applied equally at all spatial scales.


Input Requirements:
'''''''''''''''''''''
Requires a 32-bit image. If an ROI is selected it will use it, otherwise it will
process the whole image.

The size of the selection must be larger than 2^nOctaves otherwise
the entire image will be a single pixel at larger scales. If the SD ("energy")
is zero at a given scale then avoid using that scale or greater, and probably
avoid using the first smaller scale.

*/
public class DoG_ROI_bandpass_smooth implements PlugIn {
public void run(String arg) {


	ImagePlus imp = IJ.getImage();

	// ---------------------- Calculate Gabor Filter Stuff-------------------------------

	double sigma1 = 2.0;
	double sigma2 = 3.2;
	int nOctaves = 8;
	String imTitle = imp.getTitle();

	GenericDialog gd = new GenericDialog("DoG Convolution");
		gd.addMessage("Gaussian Filter Settings");
		gd.addNumericField("Sigma1", sigma1, 2);
		gd.addNumericField("Sigma2", sigma2, 2);
		gd.addNumericField("Number_of_octaves", nOctaves, 0);
		gd.addMessage("Output Images");
		gd.addCheckbox("DoG Kernel", false);
		gd.addStringField("Label", imTitle, 20);

	gd.showDialog();
	if (gd.wasCanceled())
		return;


	sigma1 = gd.getNextNumber();
	sigma2 = gd.getNextNumber();
	nOctaves = (int) Math.round(gd.getNextNumber());

	boolean outputKernel = gd.getNextBoolean();

	imTitle = gd.getNextString();

// sigma 2 should be larger than sigma1
if(sigma1 > sigma2){
	double td = sigma2;
	sigma2 = sigma1;
	sigma1 = td;
}

//----------------calculate size of kernel---------------

double gVal = (1/Math.pow(2*Math.PI*Math.pow(sigma2,2),0.5)) * Math.exp(-1*(Math.pow(0,2)/(2*Math.pow(sigma2,2) ) ));
double minVal = 0.001; // this is the cutoff as a proportion of the peak at which it is assumed the Guassian curve is small enough to have no influence
minVal = minVal / ( (1/Math.pow(2*Math.PI*Math.pow(sigma2,2),0.5)) * Math.exp(-1*(Math.pow(0,2)/(2*Math.pow(sigma2,2) ) ))); // minVal as a proportaion of start val
int cutoff = 1;

while(gVal > minVal){
	gVal = (1/Math.pow(2*Math.PI*Math.pow(sigma2,2),0.5)) * Math.exp(-1*(Math.pow(cutoff,2)/(2*Math.pow(sigma2,2) ) ));
	cutoff++;
}


// Gaussian Kernel
int mapW = 2*cutoff +1;
double[] kernelArray = new double[mapW*mapW]; 
double rad = 0.0;

// create normalised kernel where mean = 0
double cSum = 0.0; // centre sum
double sSum = 0.0; // surround sum
double csSD = 0.0; // sum of squares  for SD

//---- Calculate the area under the curve for centre and surround Gaussian sigma value for normalisation

for(int y=0; y<mapW; y++)
for(int x=0; x<mapW; x++){
	rad = Math.pow( Math.pow(x-cutoff,2) + Math.pow(y-cutoff,2) ,0.5);
	sSum += (1/Math.pow(2*Math.PI*Math.pow(sigma2,2),0.5)) * Math.exp(-1*(Math.pow(rad,2)/(2*Math.pow(sigma2,2) ) ));
	cSum += (1/Math.pow(2*Math.PI*Math.pow(sigma1,2),0.5)) * Math.exp(-1*(Math.pow(rad,2)/(2*Math.pow(sigma1,2) ) ));
}


for(int y=0; y<mapW; y++)
for(int x=0; x<mapW; x++){
	rad = Math.pow( Math.pow(x-cutoff,2) + Math.pow(y-cutoff,2) ,0.5);
	kernelArray[(y*mapW)+x] = -((1/Math.pow(2*Math.PI*Math.pow(sigma2,2),0.5)) * Math.exp(-1*(Math.pow(rad,2)/(2*Math.pow(sigma2,2) ) )))/sSum;
	kernelArray[(y*mapW)+x] += ((1/Math.pow(2*Math.PI*Math.pow(sigma1,2),0.5)) * Math.exp(-1*(Math.pow(rad,2)/(2*Math.pow(sigma1,2) ) )))/cSum;
	csSD += Math.pow(kernelArray[(y*mapW)+x] , 2);
}

// ---- normalise Kernel so that SD=1---------

csSD = Math.pow(csSD/(mapW*mapW), 0.5);

for(int y=0; y<mapW; y++)
for(int x=0; x<mapW; x++){
	kernelArray[(y*mapW)+x] = kernelArray[(y*mapW)+x] / csSD;
}


	if(outputKernel == true){
		ImageStack kernelStack = new ImageStack(mapW, mapW);
		float[] kernelOutputArray = new float[mapW*mapW];
		for(int i=0; i<mapW*mapW; i++)
			kernelOutputArray[i] = (float) kernelArray[i];
		kernelStack.addSlice("s" + sigma1 + "_" +sigma2, kernelOutputArray);
		new ImagePlus("DoG Kernel", kernelStack).show();
	}


	// --------------------- Get ROI Mask and Outline---------------------------------


	Roi roi = imp.getRoi();
	if (roi!=null && !roi.isArea()) roi = null;
	ImageProcessor ip = imp.getProcessor();
	int w = ip.getWidth();
	int h = ip.getHeight();
	int roiW = 0;
	int roiH = 0;
	int roiX = 0;
	int roiY = 0;

	//ImageProcessor mask = new ByteProcessor(roiW, roiH);

	if (roi==null) // in case there is no ROI selected use the entire image
		roi = new Roi(0, 0, w, h);
	
	Rectangle r = roi.getBounds();
	ImageProcessor mask = roi.getMask();

	//int roiType = roi.getType();
	//if (roiType==Roi.RECTANGLE){
	if(mask==null){
		//IJ.log("rectangle");
		mask = new ByteProcessor(r.width, r.height);
		mask.invert();
	}

	roiW = r.width;
	roiH = r.height;
	roiX = r.x;
	roiY = r.y;


ImageStack outStack = new ImageStack(roiW, roiH);
double kernelSum = 0.0; // the kernel needs re-normalising with each pass
int kernelCount = 0;

float[] pixels = new float[roiW*roiH];

for (int y=0; y<roiH; y++)
for (int x=0; x<roiW; x++)
if(mask.getPixel(x,y) != 0){
	pixels[x+ y*roiW] = ip.getPixelValue(x+roiX, y+roiY);
}


ResultsTable rt =  ResultsTable.getResultsTable();
rt.incrementCounter();
rt.addValue("Title", imTitle);


//---------------Bicubic interpolation coefficients----------------
double[][] M = {             // Catmull-Rom basis matrix
	{-0.5,  1.5, -1.5, 0.5},
	{ 1  , -2.5,  2  ,-0.5},
	{-0.5,  0  ,  0.5, 0  },
	{ 0  ,  1  ,  0  , 0  }
};

double[][] C = new double[4][4];
double[][] T = new double[4][4];
double[][] G = new double[4][4];

double ix = 0.0;
double iy = 0.0;


//-------------------------------- downscale pixels-------------------------------------
// sampling at an octave scale

int scaleRad = 1;

for(int k=0; k<nOctaves; k++){

scaleRad = (int) Math.pow(2, k);

/*
int scaleW = 0;
if( (double) (roiW/scaleRad) == (double) Math.round(roiW/scaleRad))
	scaleW = (int) Math.round(r.width/scaleRad);
else scaleW = (int) (Math.floor(roiW/scaleRad) +1);

int scaleH = 0;
if( (double) (roiH/scaleRad) == (double) Math.round(roiH/scaleRad))
	scaleH = (int) Math.round(roiH/scaleRad);
else scaleH = (int) (Math.floor(roiH/scaleRad) +1);
*/

int scaleW = (int) (Math.floor(roiW/scaleRad) +1);
int scaleH = (int) (Math.floor(roiH/scaleRad) +1);

double scaleSum = 0.0;
int scaleCount = 0;


float[] scalePixels = new float[scaleW*scaleH];
int[] scaleMask = new int[scaleW*scaleH];

for(int y=0; y<scaleH; y++)
for(int x=0; x<scaleW; x++){

	scaleSum = 0.0;
	scaleCount = 0;

	for(int y2 = 0; y2<scaleRad; y2++)
	for(int x2 = 0; x2<scaleRad; x2++)
	if(mask.getPixel(x2+x*scaleRad,y2+y*scaleRad) != 0){
		scaleSum += pixels[(x2+x*scaleRad) + (roiW*(y2+y*scaleRad))];
		scaleCount ++;
	} //x2 y2 fi

	if(scaleCount > 0){
		scalePixels[x+y*scaleW] = (float) (scaleSum/scaleCount);
		scaleMask[x+y*scaleW] = 1;
	}
} //xy

//ImageStack scaleStack = new ImageStack(scaleW, scaleH);
//scaleStack.addSlice("scaled pixels " + k, scalePixels);
//new ImagePlus("Scale " + scaleRad, scaleStack).show();


//-----------------Scale convolution-------------

//ImageStack scaleOutStack = new ImageStack(scaleW, scaleH);

//for(int j=0; j<nAngles; j++){

//int j=3;

double[] pixels2 = new double[scaleW*scaleH];
int[] counts = new int[scaleW*scaleH];
float[] outPxs = new float[scaleW*scaleH];

int loc = 0;
int pxloc = 0;
int scaleLoc = 0;

for(int y=0; y<scaleH; y++)
for(int x=0; x<scaleW; x++)
if(scaleMask[x + scaleW*y] == 1){

	pxloc = (y*scaleW)+x;

	// calculate kernel sum (it needs re-normalising at each point around the ROI)
	kernelSum = 0.0;
	kernelCount = 0;

	for(int y2=-cutoff; y2<=cutoff; y2++)  
	for(int x2=-cutoff; x2<=cutoff; x2++){
		//if(mask.getPixel(x+x2,y+y2) != 0){
		if(y+y2>=0 && y+y2< scaleH && x+x2 >=0 &&x+x2 < scaleW && scaleMask[x + x2 + scaleW*(y+y2)] == 1){
			loc = ((y2+cutoff)*mapW) + (x2+cutoff);
			kernelSum += kernelArray[loc];
			kernelCount++;
		}
	}

	kernelSum = kernelSum/kernelCount;

	for(int y2=-cutoff; y2<=cutoff; y2++)  
	for(int x2=-cutoff; x2<=cutoff; x2++){
		//if(mask.getPixel(x+x2,y+y2) != 0){
		if(y+y2>=0 && y+y2< scaleH && x+x2 >=0 &&x+x2 < scaleW && scaleMask[x + x2 + scaleW*(y+y2)] == 1){
			loc = ((y2+cutoff)*mapW) + (x2+cutoff);
			pixels2[pxloc] = pixels2[pxloc] + (scalePixels[x+x2 + (scaleW*(y+y2))] * (kernelArray[loc]-kernelSum));
			counts[pxloc] ++;
		}
	}

	IJ.showProgress((float) y/roiH );
	
} // xy fi


for(int i=0; i<scaleH*scaleW; i++)
	outPxs[i] = (float) (pixels2[i]/counts[i]);

//scaleOutStack.addSlice("output angle" + j, outPxs);


//-----------------Apply to original scale (allows for easier measurement and weighting----------


float[] scaleOutPxs = new float[roiW*roiH];

for(int i=0; i<roiW*roiH; i++)
	scaleOutPxs[i] = Float.NaN;

double td = 0.0;
double pxSum = 0.0;
double nPx = 0.0;

for(int i=0; i<scaleW*scaleH; i++)
	pixels2[i] = pixels2[i]/counts[i];


for(int y=-1; y<scaleH; y++)
for(int x=-1; x<scaleW; x++){

	if(scaleRad > 1){
		for(int yy = -1; yy<3; yy++)
		for(int xx = -1; xx<3; xx++){
			if(y+yy >= 0
				&& y+yy < scaleH
				&& x+xx >= 0
				&& x+xx < scaleW
				&& scaleMask[(x+xx) + scaleW*(y+yy)] == 1)
				G[xx+1][yy+1] = pixels2[x+xx+scaleW*(y+yy)];
			else G[xx+1][yy+1] = 0.0;
	
			C[xx+1][yy+1] = 0.0;
			T[xx+1][yy+1] = 0.0;
		}
	
		for (int ii = 0 ; ii < 4 ; ii++)    // T = G * MT
		for (int jj = 0 ; jj < 4 ; jj++)
		for (int kk = 0 ; kk < 4 ; kk++)
			T[ii][jj] += G[ii][kk] * M[jj][kk];

		for (int ii = 0 ; ii < 4 ; ii++)    // C = M * T
		for (int jj = 0 ; jj < 4 ; jj++)
		for (int kk = 0 ; kk < 4 ; kk++)
			C[ii][jj] += M[ii][kk] * T[kk][jj];
	} //fi scaleRad >1


	for(int y2 = 0; y2<scaleRad; y2++)
	for(int x2 = 0; x2<scaleRad; x2++)
	if(mask.getPixel(x2+(int) Math.floor(scaleRad/2)+x * scaleRad, y2 + (int) Math.floor(scaleRad/2)+ y*scaleRad) != 0){
		//ix = x2*(0.5/scaleRad); // the scaled XY coordinates within each pixel
		//iy = y2*(0.5/scaleRad);

		if(scaleRad > 1){
			iy = y2/((double)scaleRad); // the scaled XY coordinates within each pixel
			ix = x2/((double)scaleRad);

			td = iy*(iy*(iy*(ix * (ix * (ix * C[0][0] + C[1][0]) + C[2][0]) + C[3][0])	
			                     + (ix * (ix * (ix * C[0][1] + C[1][1]) + C[2][1]) + C[3][1]))
			                  +    (ix * (ix * (ix * C[0][2] + C[1][2]) + C[2][2]) + C[3][2]))
			               +       (ix * (ix * (ix * C[0][3] + C[1][3]) + C[2][3]) + C[3][3]);
			//scaleOutPxs[ x2+ (int) Math.floor(scaleRad/2)+x*scaleRad + roiW*(y2+ (int) Math.floor(scaleRad/2)+y*scaleRad)] = (float) td;
			scaleOutPxs[ x2+ (int) Math.floor(scaleRad/2)+x*scaleRad + roiW*(y2+ (int) Math.floor(scaleRad/2)+y*scaleRad)] = (float) td;

		} else { // no interpolation
			td = pixels2[x+scaleW*y];
			scaleOutPxs[ x2+x*scaleRad + roiW*(y2+y*scaleRad)] = (float) td;
		}

		pxSum += td;
		nPx ++;
	}
}//yx mask

double pxMean = pxSum/nPx;
double pxSSum = 0.0;


for(int y=0; y<roiH; y++)
for(int x=0; x<roiW; x++)
if(mask.getPixel(x, y) != 0)
	pxSSum += Math.pow( scaleOutPxs[x+roiW*y]-pxMean,2);

//IJ.log("scale:" + k+ " angle:" + j + "mean: " + pxMean + " SD:" + Math.pow(pxSSum/nPx, 0.5));

	// ------------------ OUTPUT RESULTS--------------------------

	// if adding new row:
	//rt.incrementCounter();
	//rt.addValue("Title", imTitle);
	//rt.addValue("Scale", k);
	//rt.addValue("Angle", j);
	//rt.addValue("Energy", Math.pow(pxSSum/nPx, 0.5));
	rt.addValue("s"+k, Math.pow(pxSSum/nPx, 0.5));

outStack.addSlice("scale" +k, scaleOutPxs);

//}//j angles

IJ.showProgress((float) 1);

//new ImagePlus("Scaled Gabor output", scaleOutStack).show();

}//k

new ImagePlus("DoG output", outStack).show();

rt.showRowNumbers(true);
rt.show("Results");

}
}
```
