## Supplementary figures and images for "Shining a light on camouflage evolution: using genetic algorithms to determine the effects of geometry and lighting on optimal camouflage"

### eggPatterns.jpg

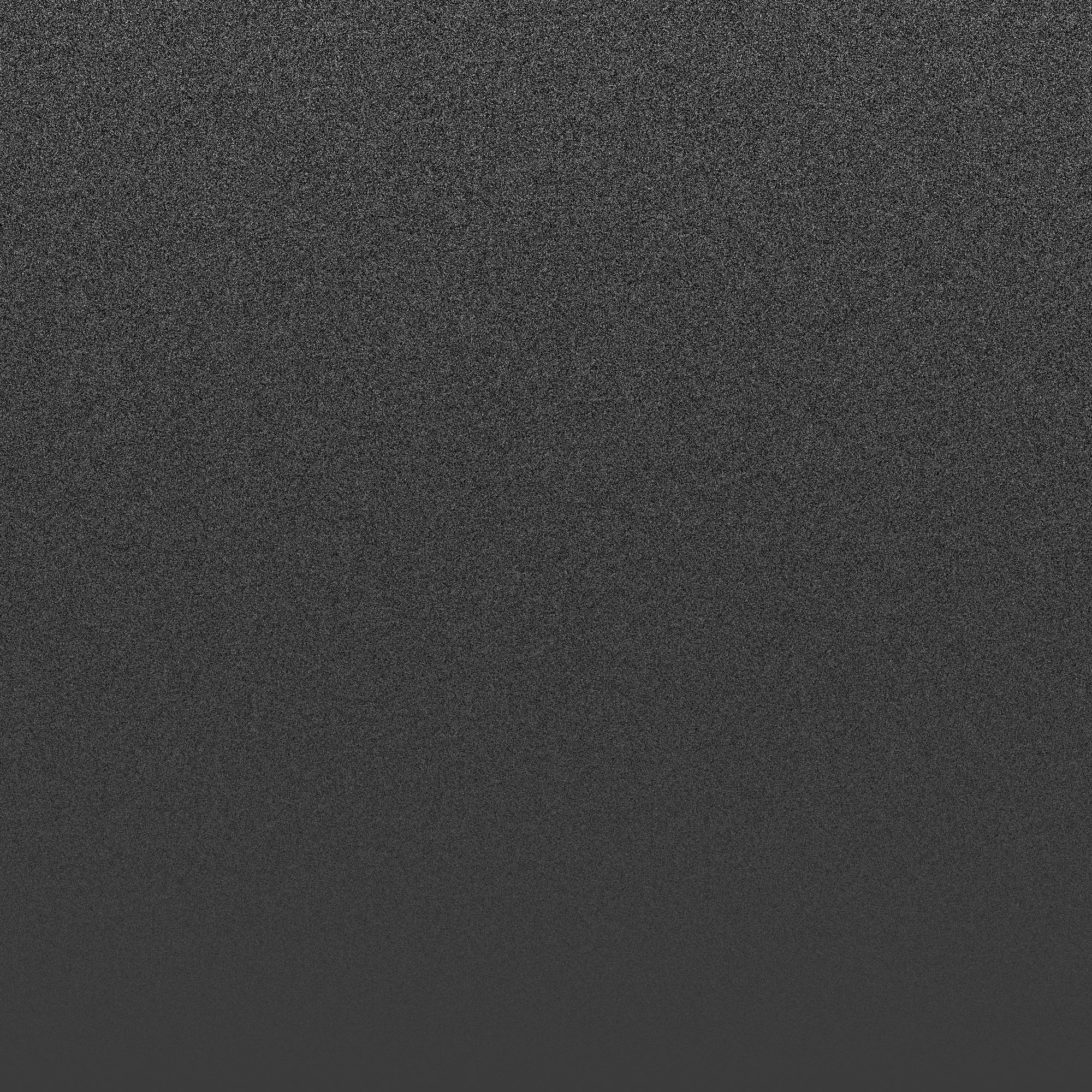

### Gen13_Mut0_ID8.tif

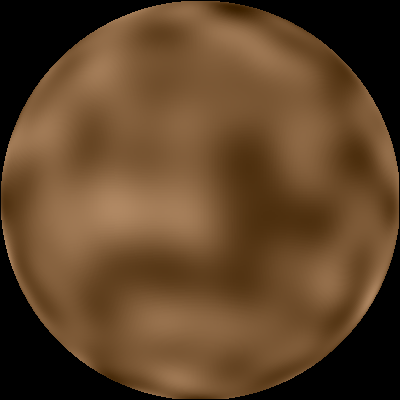

### Gen16_Mut0_ID1.tif

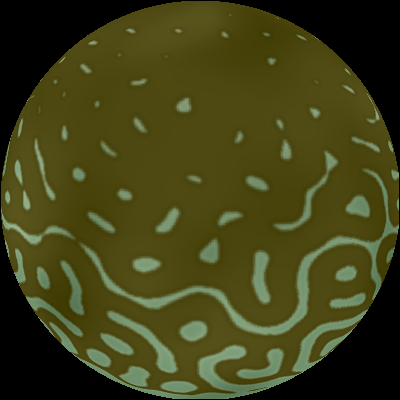

### Gen16_Mut0_ID3.tif

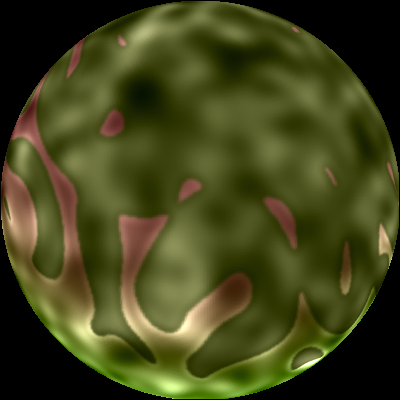

### Gen16_Mut0_ID6.tif

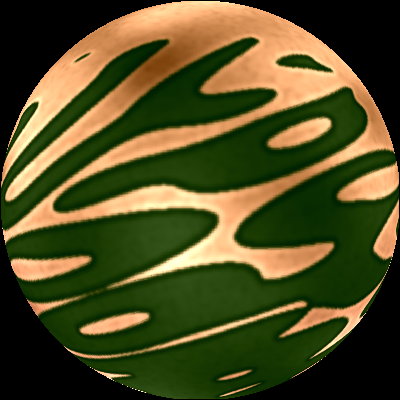

### Gen16_Mut0_ID6.tif

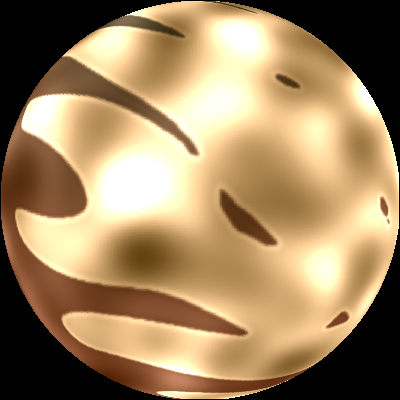

### Gen16_Mut0_ID11.tif

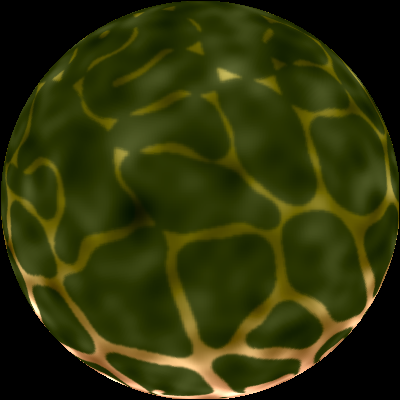

### Gen16_Mut0_IDR1.tif

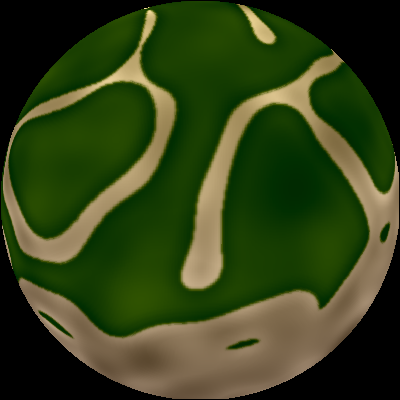

### Gen17_Mut0_ID3.tif

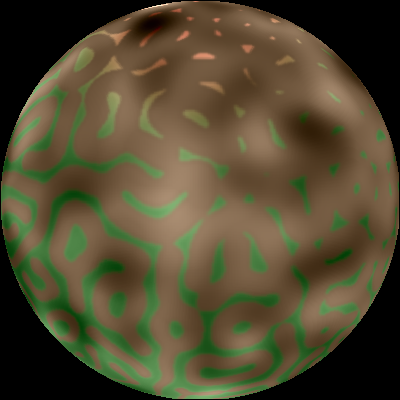

### Gen17_Mut0_ID6.tif

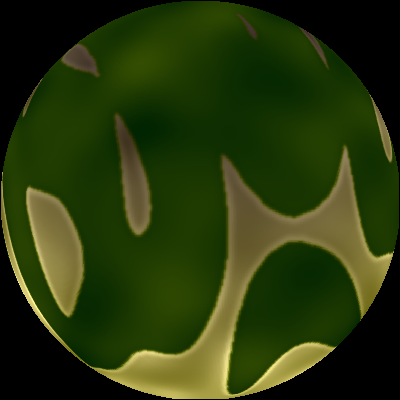

### Gen17_Mut0_ID11.tif

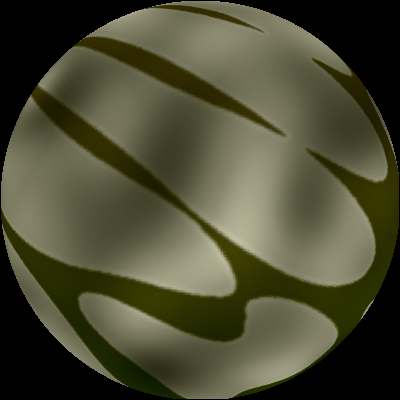

### Gen18_Mut0_ID0.tif

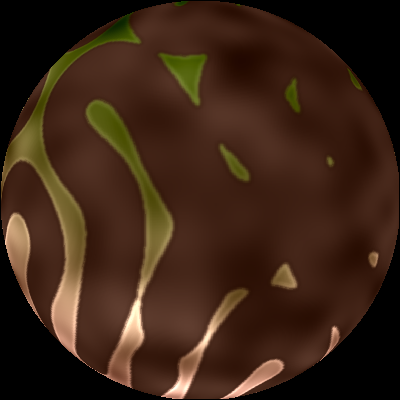

### Gen18_Mut0_ID0.tif

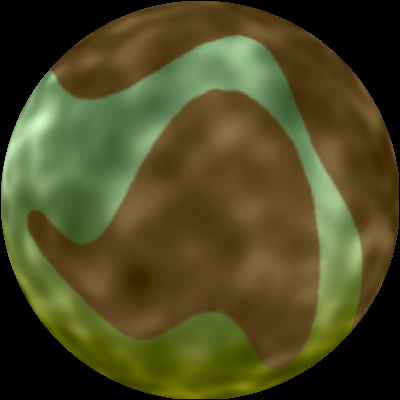

### Gen18_Mut0_ID0.tif

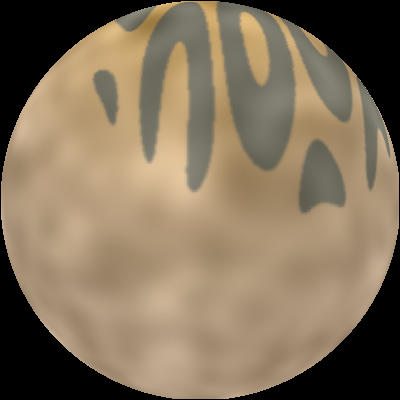

### Gen18_Mut0_ID1.tif

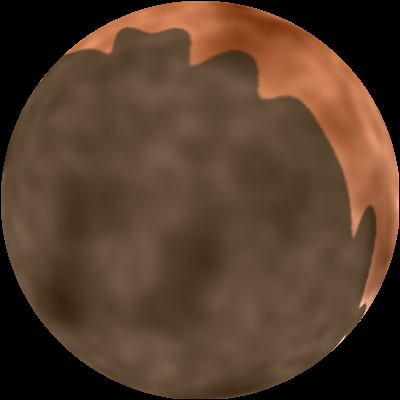

### Gen18_Mut0_ID1.tif

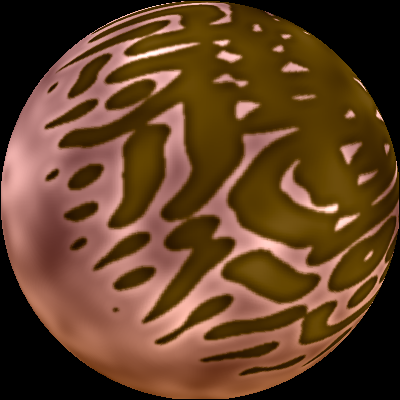

### Gen18_Mut0_ID1.tif

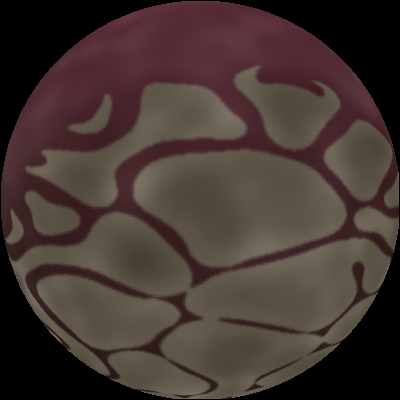

### Gen18_Mut0_ID1.tif

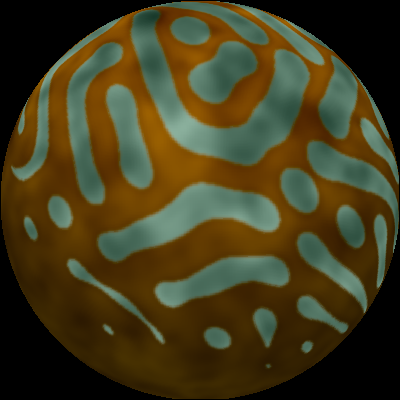

### Gen18_Mut0_ID1.tif

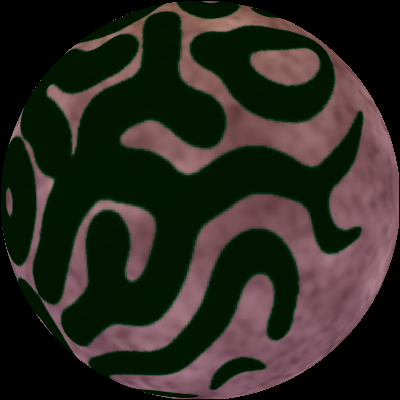

### Gen18_Mut0_ID3.tif

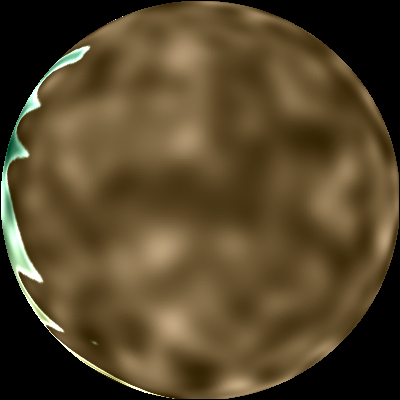

### Gen18_Mut0_ID4.tif

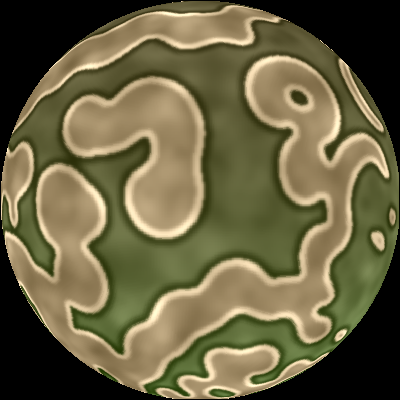

### Gen18_Mut0_ID8.tif

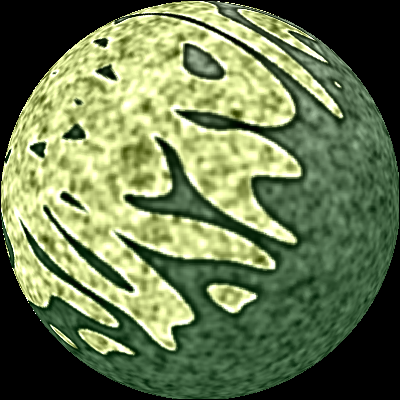

### Gen18_Mut0_ID10.tif

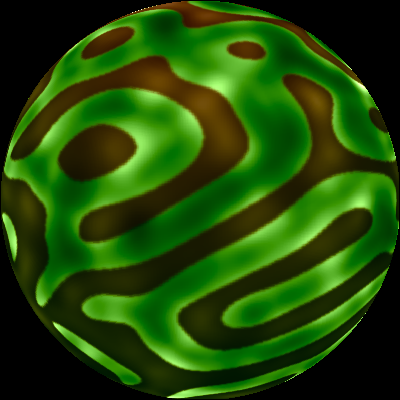

### Gen18_Mut0_ID11.tif

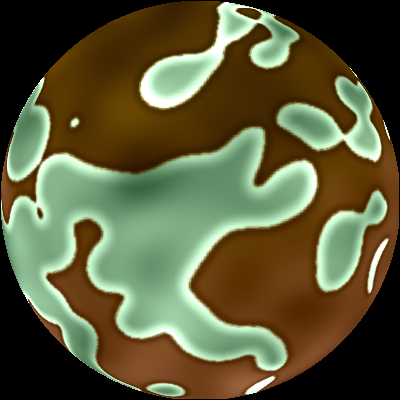

### Gen18_Mut0_IDR1.tif

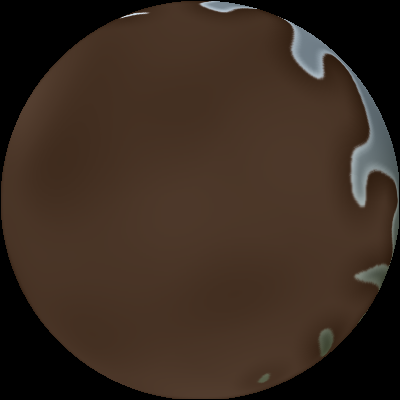

### Gen19_Mut0_ID0.tif

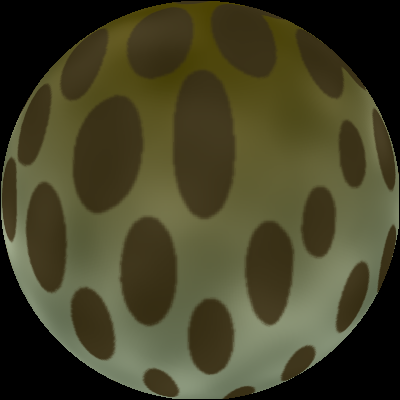

### Gen19_Mut0_ID1.tif

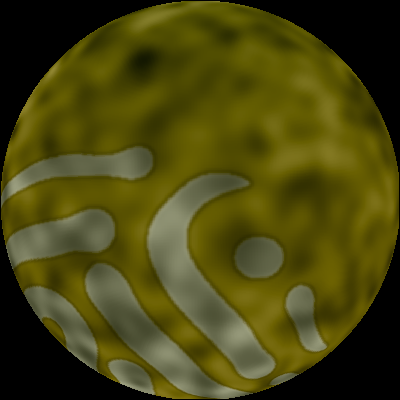

### Gen19_Mut0_ID2.tif

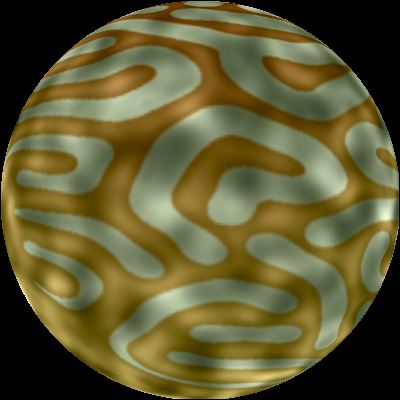

### Gen19_Mut0_ID2.tif

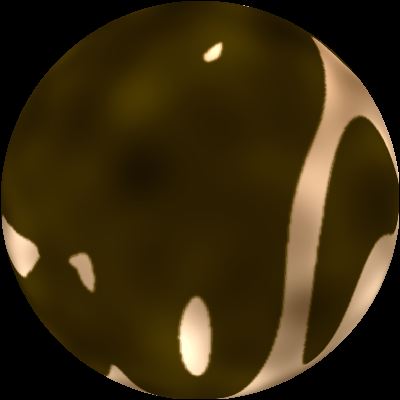

### Gen19_Mut0_ID3.tif

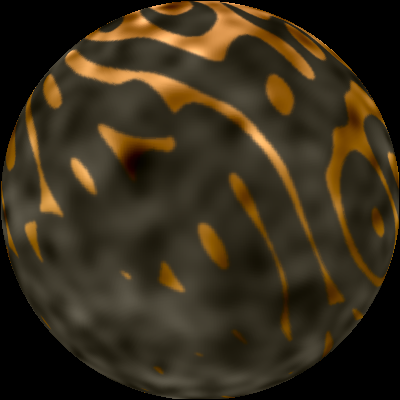

### Gen19_Mut0_ID4.tif

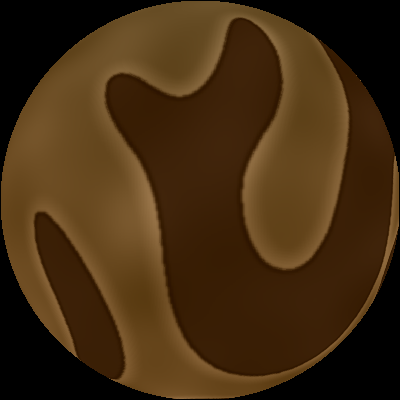

### Gen19_Mut0_ID4.tif

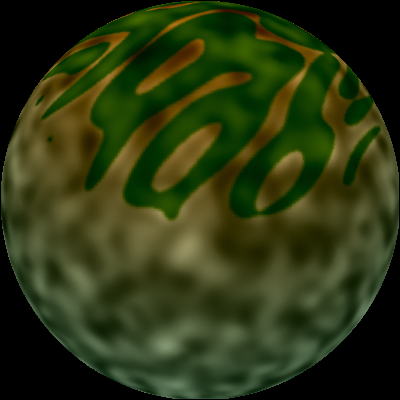

### Gen19_Mut0_ID4.tif

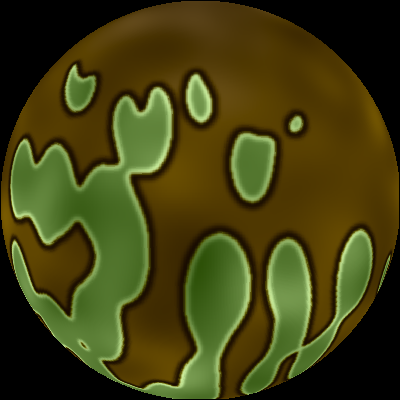

### Gen19_Mut0_ID8.tif

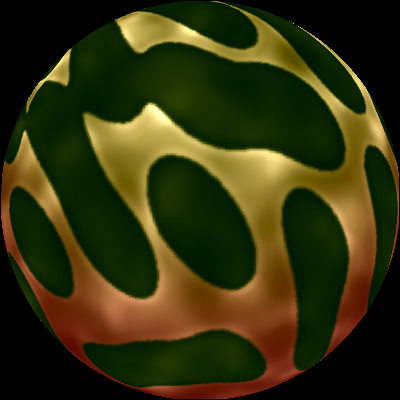

### Gen19_Mut0_ID9.tif

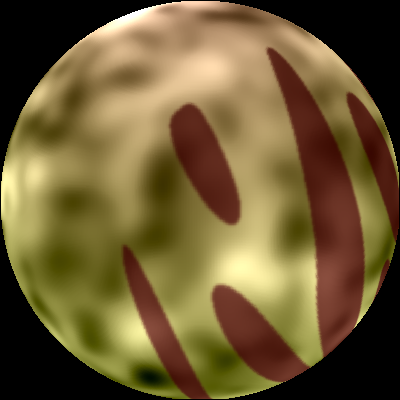

### Gen19_Mut0_ID9.tif

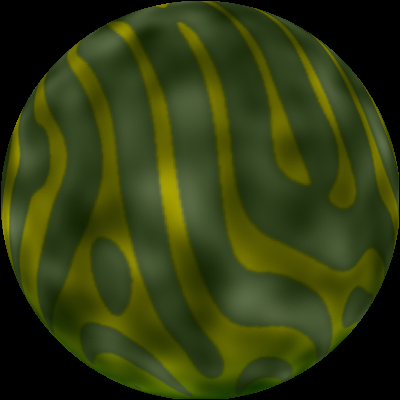

### Gen19_Mut0_ID9.tif

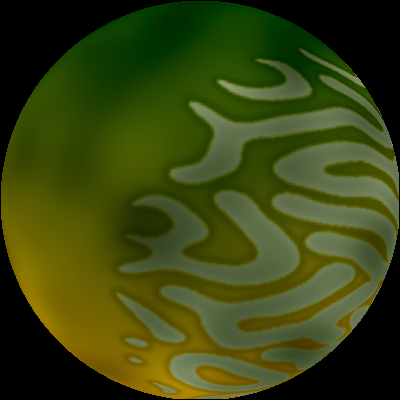

### Gen19_Mut0_ID9.tif

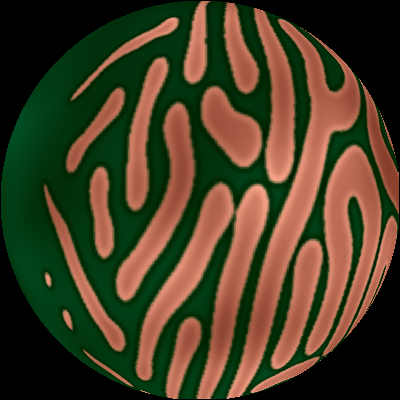

### Gen19_Mut0_ID9.tif

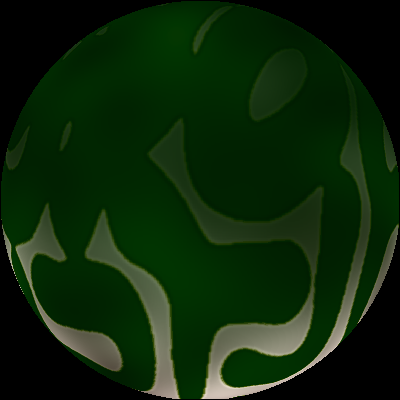

### Gen19_Mut0_ID9.tif

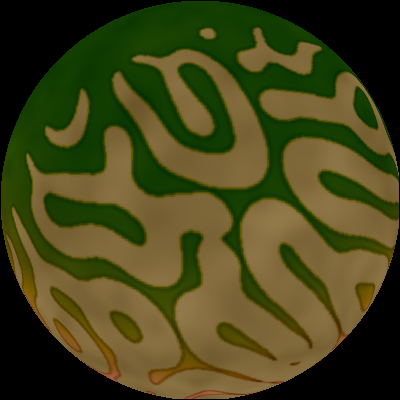

### Gen19_Mut0_ID10.tif

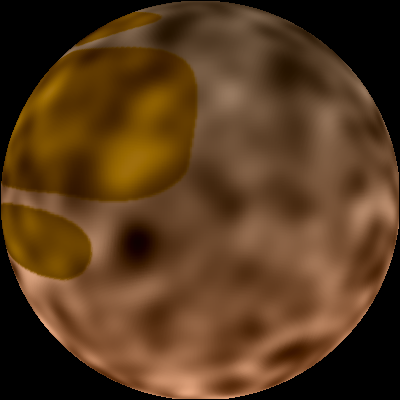

### Gen19_Mut0_IDR0.tif

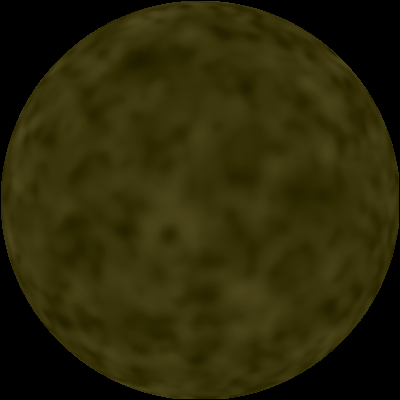

### Gen20_Mut0_ID0.tif

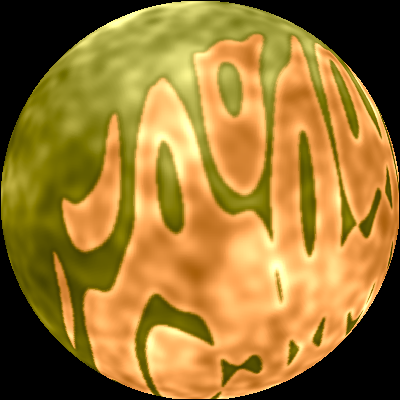

### Gen20_Mut0_ID0.tif

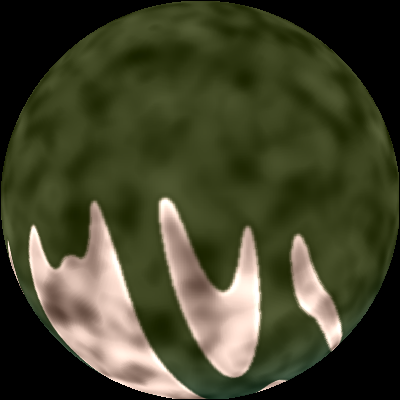

### Gen20_Mut0_ID0.tif

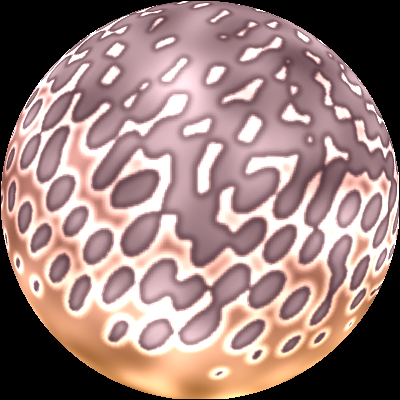

### Gen20_Mut0_ID1.tif

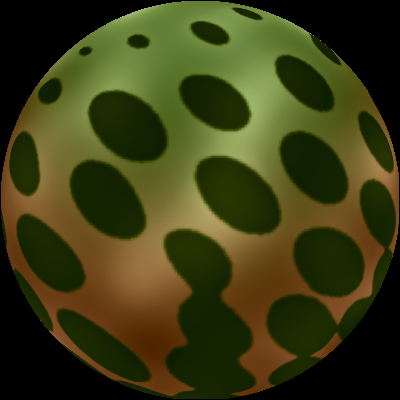

### Gen20_Mut0_ID1.tif

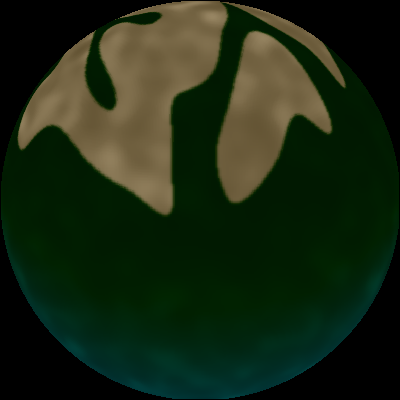

### Gen20_Mut0_ID1.tif

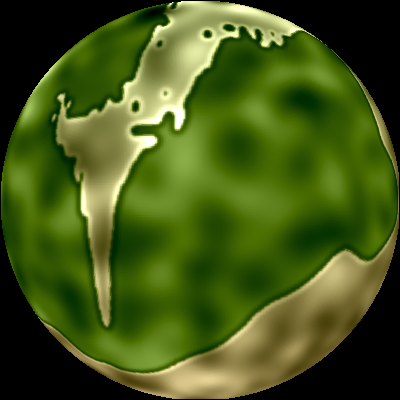

### Gen20_Mut0_ID1.tif

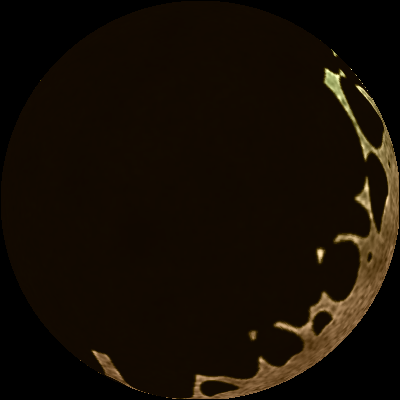

### Gen20_Mut0_ID1.tif

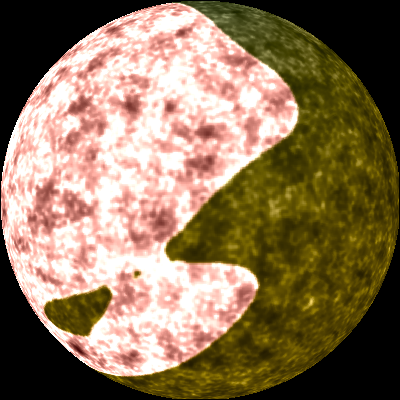

### Gen20_Mut0_ID2.tif

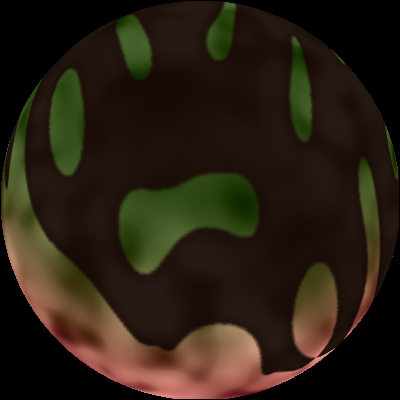

### Gen20_Mut0_ID2.tif

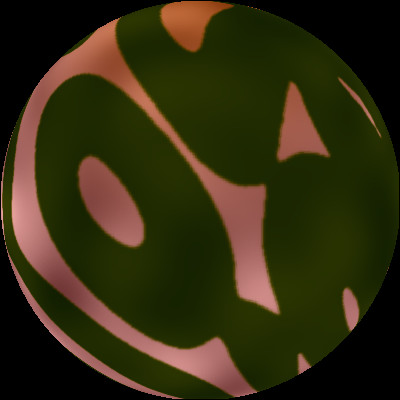

### Gen20_Mut0_ID3.tif

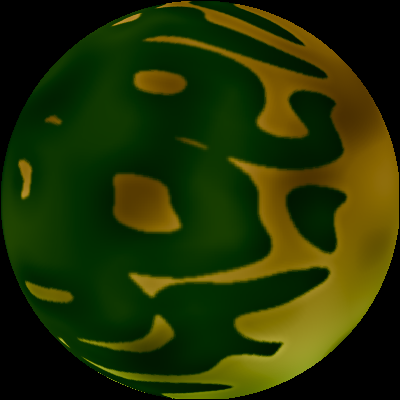

### Gen20_Mut0_ID3.tif

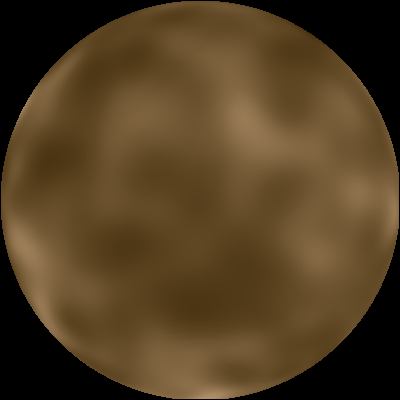

### Gen20_Mut0_ID3.tif

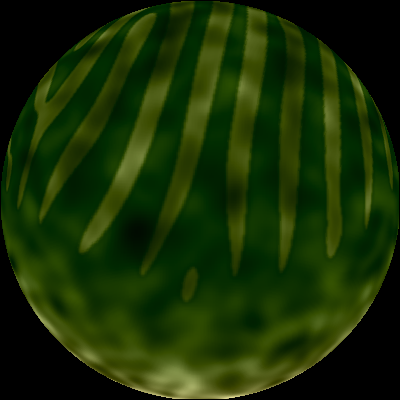

### Gen20_Mut0_ID3.tif

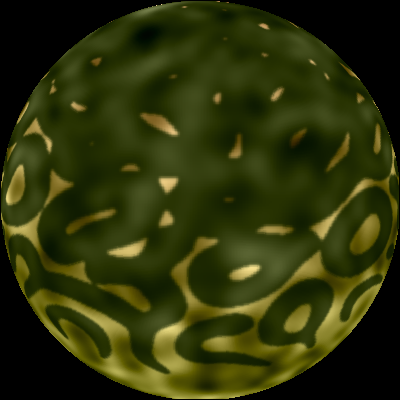

### Gen20_Mut0_ID4.tif

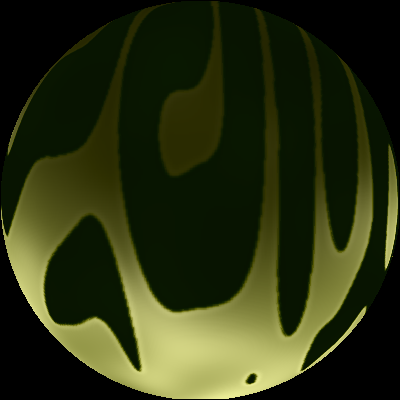

### Gen20_Mut0_ID4.tif

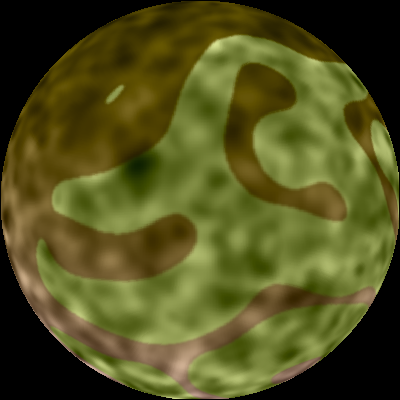

### Gen20_Mut0_ID4.tif

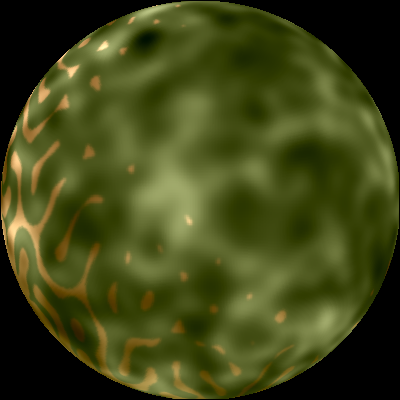

### Gen20_Mut0_ID6.tif

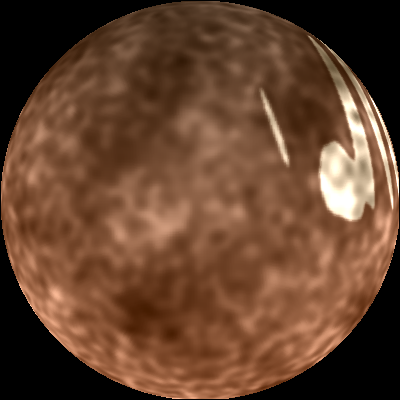

### Gen20_Mut0_ID6.tif

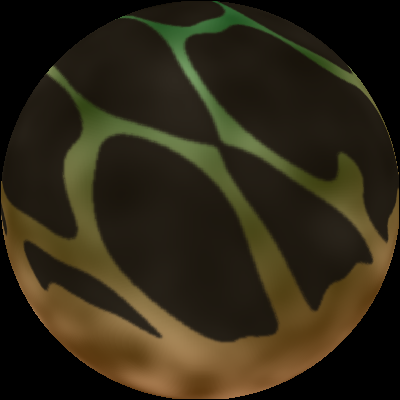

### Gen20_Mut0_ID6.tif

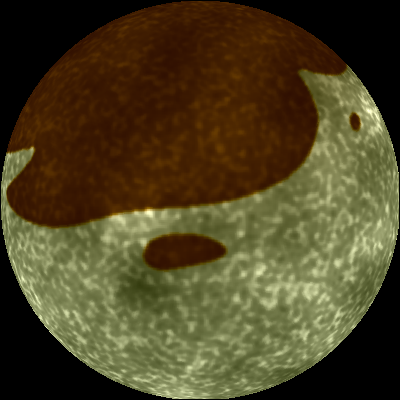

### Gen20_Mut0_ID6.tif

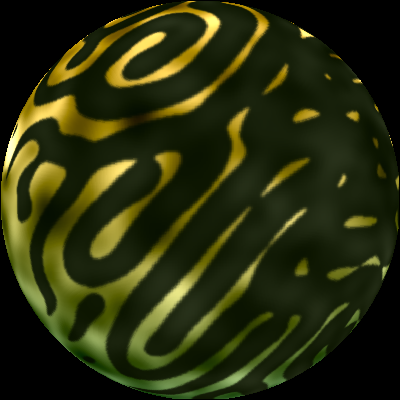

### Gen20_Mut0_ID6.tif

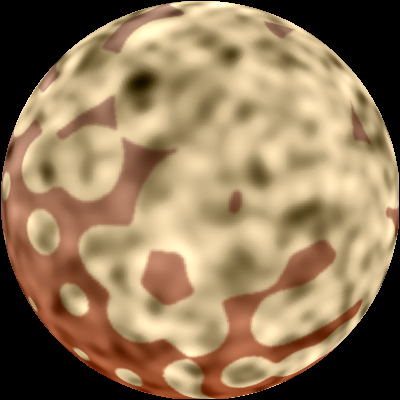

### Gen20_Mut0_ID7.tif

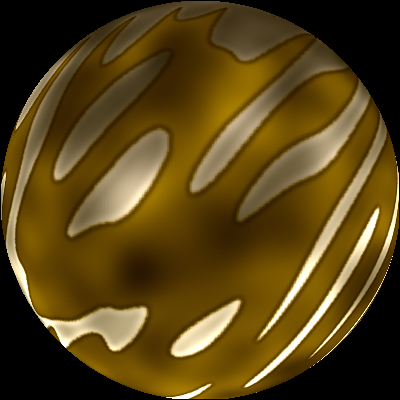

### Gen20_Mut0_ID7.tif

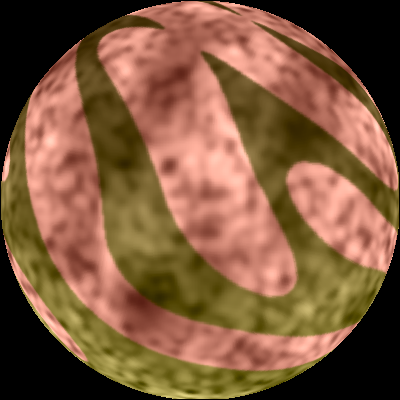

### Gen20_Mut0_ID7.tif

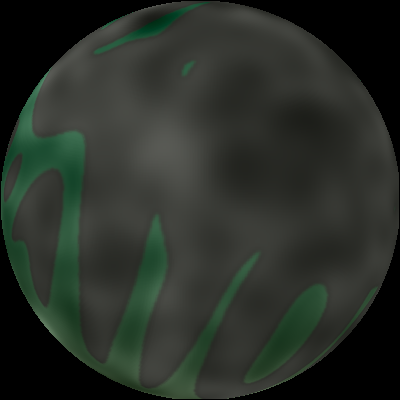

### Gen20_Mut0_ID8.tif

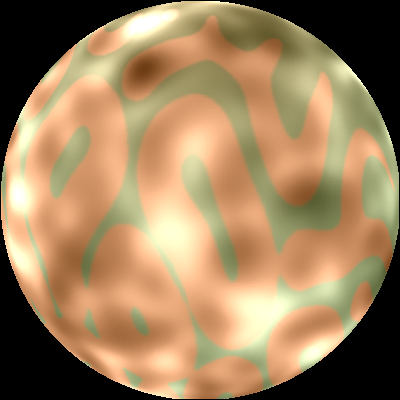

### Gen20_Mut0_ID8.tif

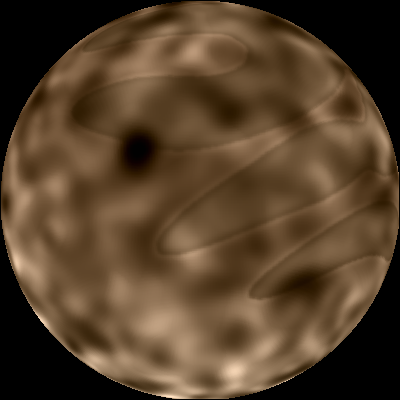

### Gen20_Mut0_ID9.tif

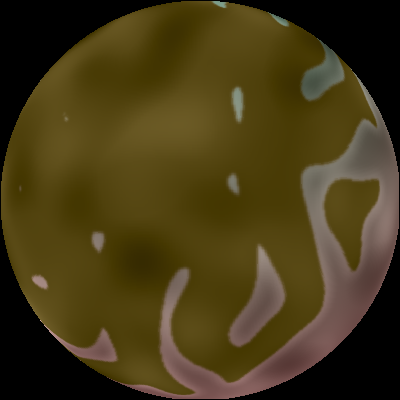

### Gen20_Mut0_ID9.tif

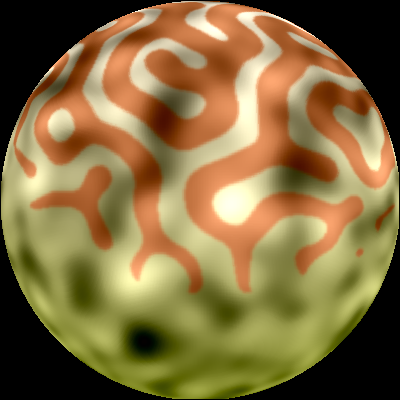

### Gen20_Mut0_ID9.tif

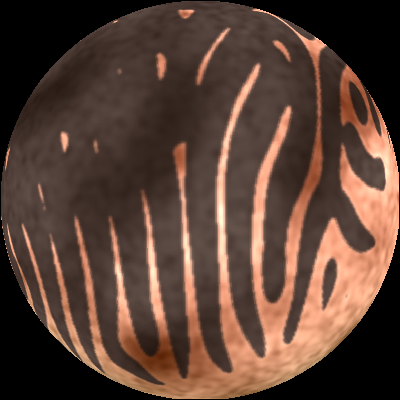

### Gen20_Mut0_ID9.tif

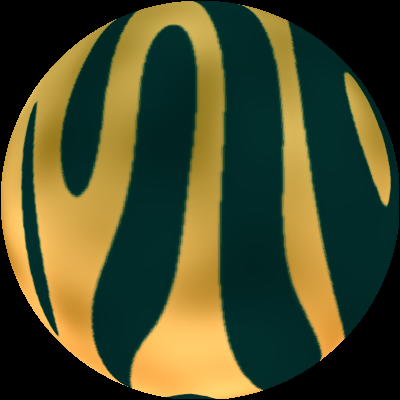

### Gen20_Mut0_ID10.tif

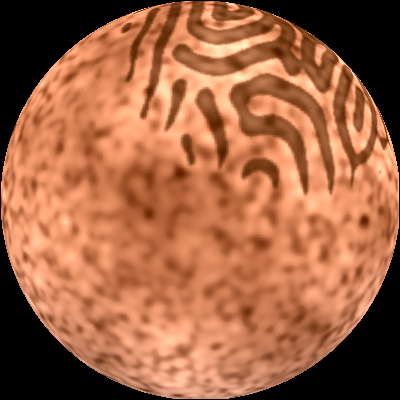

### Gen20_Mut0_ID10.tif

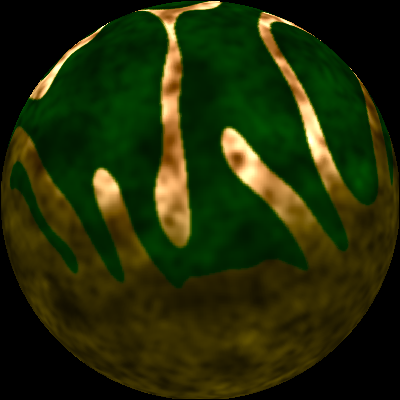

### Gen20_Mut0_ID10.tif

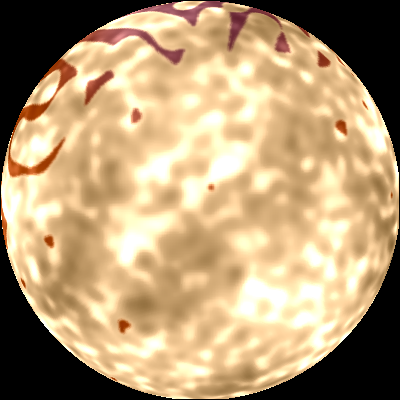

### Gen20_Mut0_ID11.tif

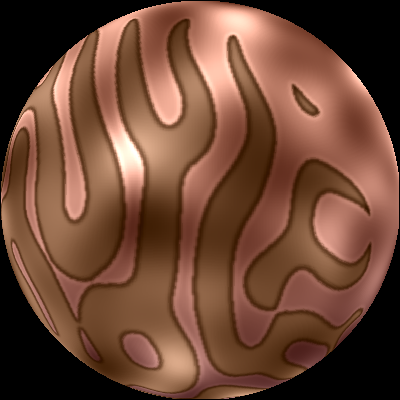

### Gen20_Mut0_ID11.tif

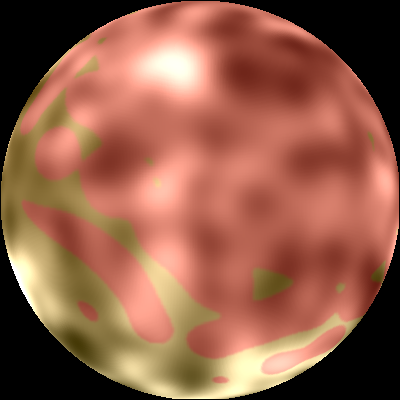

### Gen20_Mut0_ID11.tif

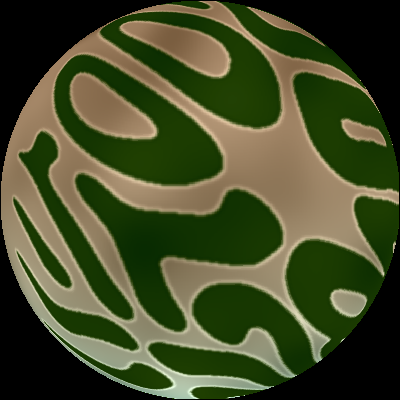

### Gen20_Mut0_IDR0.tif

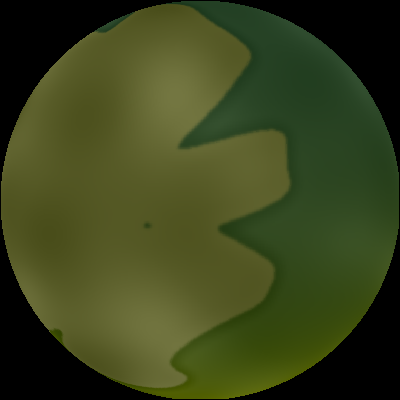

### Gen20_Mut0_IDR0.tif

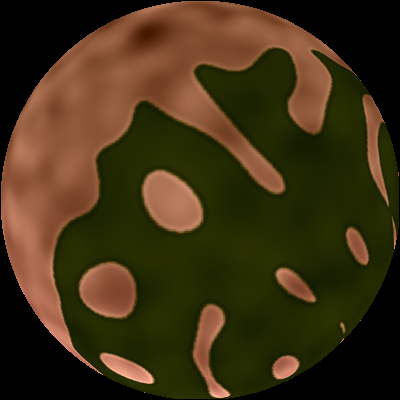

### Gen20_Mut0_IDR1.tif

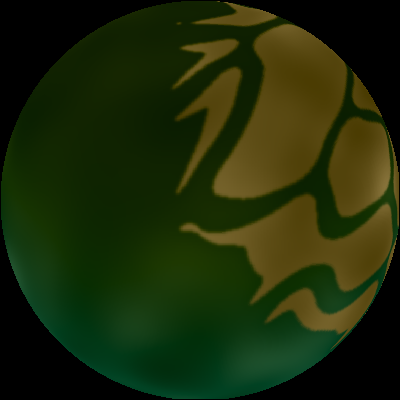

### Gen20_Mut0_IDR1.tif

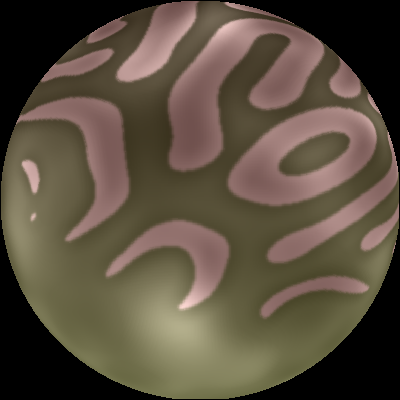

### Gen20_Mut0_IDR2.tif

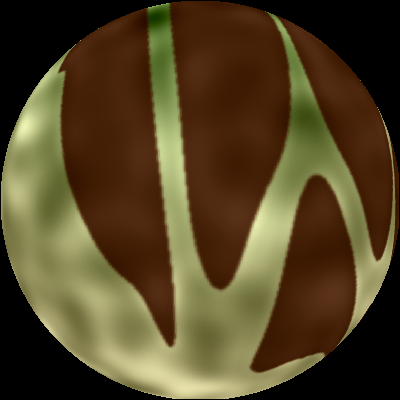

### Gen20_Mut0_IDR3.tif

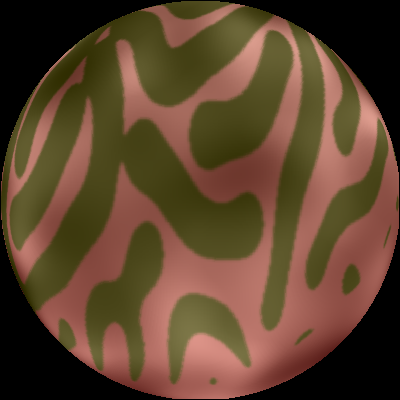

### pattern1.jpg

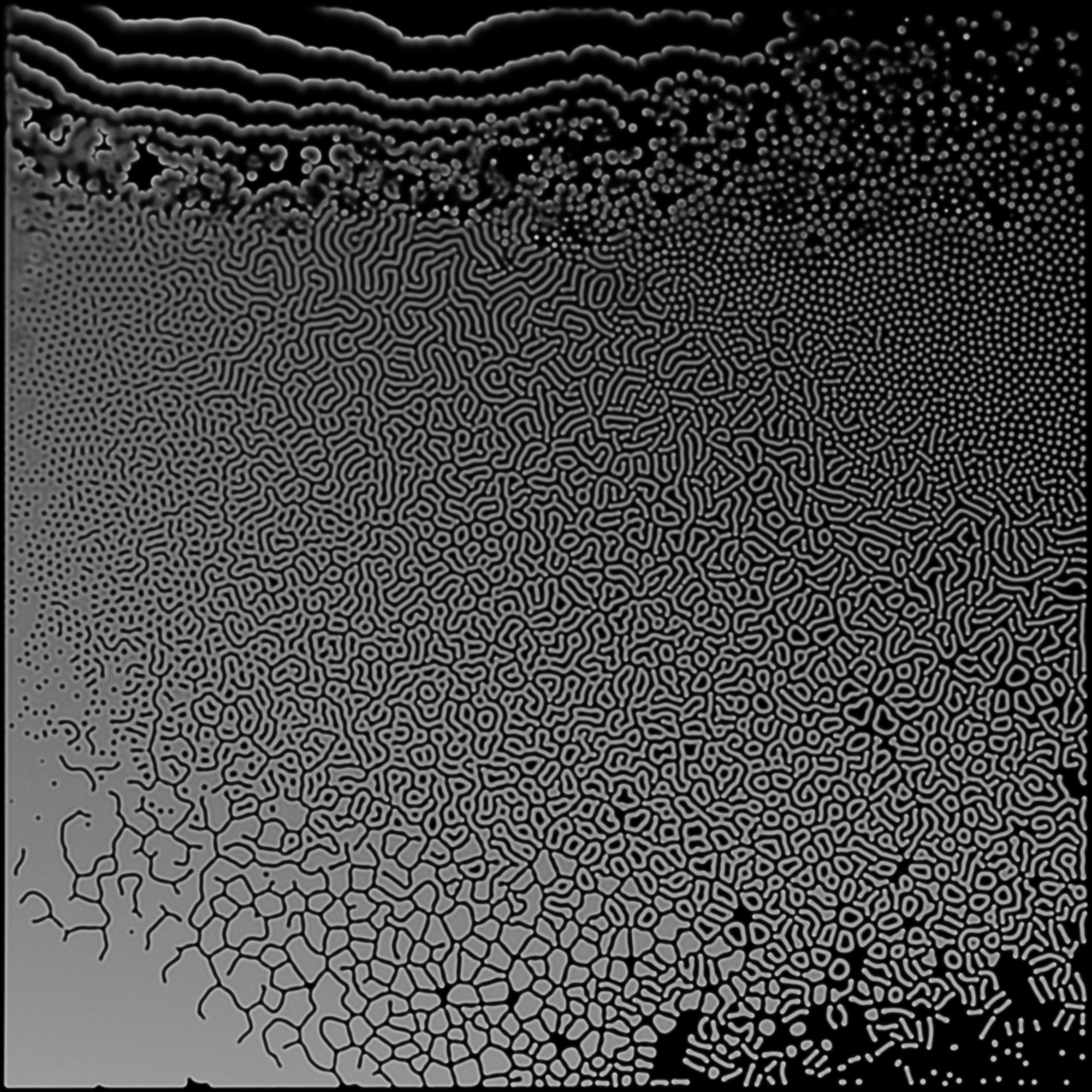
